# Exploration of the proxiOME of large subunit ribosomal proteins reveals Acl1 and Bcl1 as cooperating dedicated chaperones of Rpl1

**DOI:** 10.1101/2025.09.18.677003

**Authors:** Sébastien Favre, Benjamin Pillet, Fabiana Burchert, Devanarayanan Siva Sankar, Alfonso Méndez-Godoy, Stephan Kiontke, Jörn Dengjel, Gert Bange, Dieter Kressler

## Abstract

In eukaryotes, most newly synthesized ribosomal proteins (r-proteins) need to rapidly and safely get into the nucleus to reach their assembly site on pre-ribosomal particles. However, only for few r-proteins tailored support mechanisms involving so-called dedicated chaperones could so far be revealed. Here, with the primary aim of identifying novel dedicated chaperones, we performed TurboID-based proximity labelling with all 46 large subunit r-proteins of *Saccharomyces cerevisiae*, which unveiled the fungi-specific Acl1 and the conserved Bcl1 as candidate dedicated chaperones of Rpl1. We show that the functionally cooperating Acl1 and Bcl1 both directly interact with Rpl1, form a trimeric Acl1-Rpl1-Bcl1 complex, and enable the nuclear import of Rpl1. Moreover, our crystal structure of the minimal Acl1-Rpl1 complex reveals how Acl1’s ankyrin repeat domain shields a positively charged rRNA-binding surface of Rpl1. Our proximity labelling approach also permitted to establish novel interactions between four r-proteins and distinct importins and to illuminate r-protein neighbourhoods on successive pre-60S particles. Additionally, reciprocal proximity labelling with the known dedicated chaperones indicates that almost all appear to be transiently associated with pre-ribosomal particles. Our study provides for the first time comprehensive insight into the physical proximities of large subunit r-proteins along their entire life cycle.

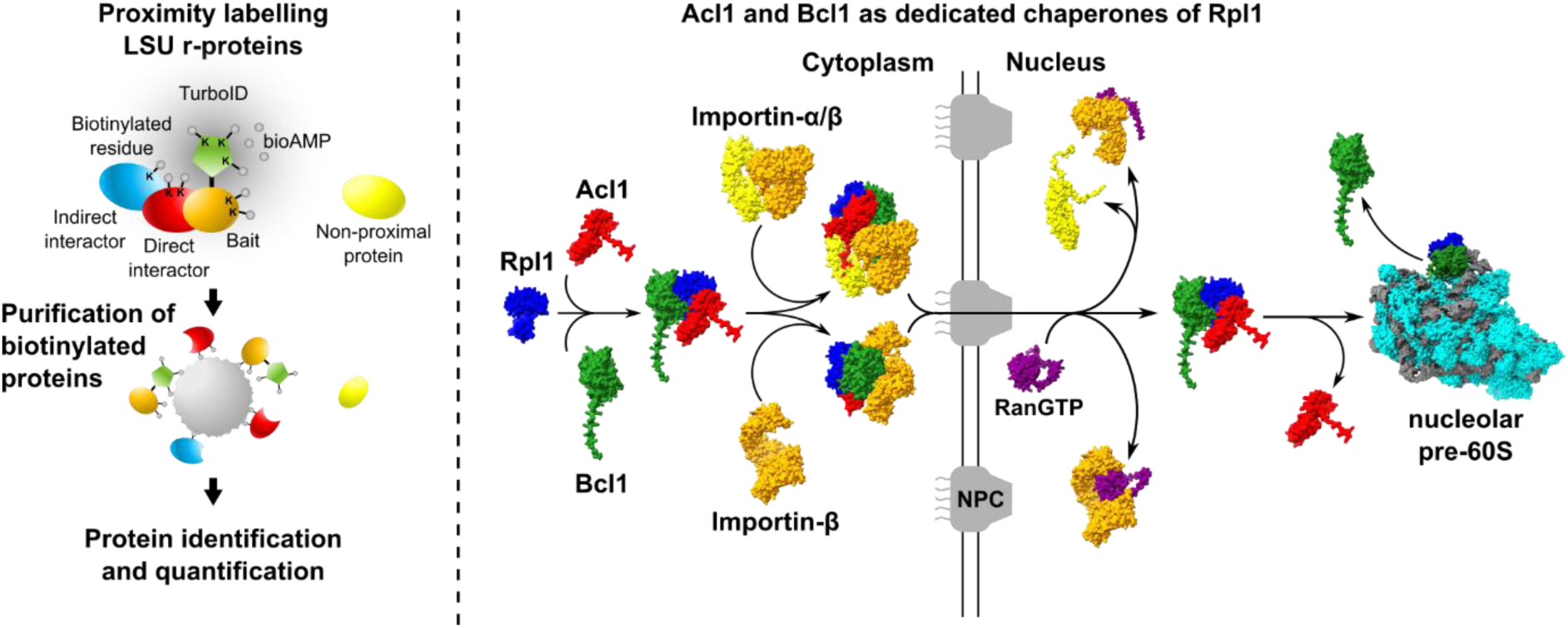
GRAPHICAL ABSTRACT.

## INTRODUCTION

Protein synthesis is an essential process accomplished by ribosomes in all organisms; accordingly, ribosomes of all three domains of life share a conserved functional core [1]. As their overall structure and composition are quite well conserved among eukaryotes, functional and structural studies of the eukaryotic ribosome and its assembly process were and are still often carried out in the budding yeast *Saccharomyces cerevisiae*, notably due to the experimental amenability of this unicellular organism. In eukaryotes, translating 80S ribosomes consist of two asymmetric subunits, referred to as the small 40S (SSU) and large 60S (LSU) ribosomal subunit. In *S. cerevisiae*, the SSU contains 33 ribosomal proteins (r-proteins) and the 18S ribosomal RNA (rRNA), whereas the LSU is composed of 46 r-proteins and three rRNAs, the 5S, 5.8S, and 25S rRNA [1,2]. In order to maintain the cellular amounts of this macromolecular nanomachine, yeast cells in exponential growth phase need to produce approximately 2000 functional ribosomes per minute [3]. This task represents a major challenge, not only to coordinate the production of the ribosomal components but also for their well-orchestrated assembly. Ribosome assembly alone requires more than 200 assembly factors (AFs) and around 80 small nucleolar RNAs (snoRNAs) to provide accuracy and speed to this multistep process, during which the r-proteins get sequentially incorporated into pre-ribosomal particles as the precursor rRNAs (pre-rRNAs) are progressively processed into mature rRNAs and, at the same time, folded into their final conformation [2,4–8].

Most r-proteins, owing to their capacity to interact with rRNA, contain highly basic regions. In unassembled r-proteins, these highly basic regions are prone to engage in non-specific interactions with polyanions, such as cytoplasmic tRNAs, which can result in r-protein aggregation [9]. Accordingly, an excess of unassembled r-proteins, in certain cases even of an individual r-protein, can lead to r-protein aggregation and elicit a proteostatic collapse [10–14]. To limit the detrimental effects of unassembled r-proteins while still producing sufficient amounts of new r-proteins to meet the actual demand of ribosome assembly, cells possess several general as well as highly specific regulation and protection mechanisms. First, the production of r-proteins is already coordinated at the transcriptional level as the synthesis of all r-protein gene (RPG) mRNAs is driven by only three distinct types of co-regulated promoters, which probably most notably differ by their diverging dependence on the TORC1-controlled transcriptional activators Ifh1 and Sfp1, thereby ensuring the provision of roughly equal amounts of each RPG mRNA in different growth conditions and, hence, at different rates of ribosome assembly [15–19]. Second, cells continuously monitor the levels of free r-proteins and respond to their excess presence by two general regulatory mechanisms, the ERISQ (excess ribosomal protein quality control) pathway and a stress response pathway termed RASTR (ribosome assembly stress response) or RPAS (ribosomal protein assembly stress) [10,12,13,16]. The ERISQ pathway selectively eliminates free r-proteins and, thus, enables cells to readily cope with a moderate excess of unassembled r-proteins. To do so, the nucle(ol)ar E3 ubiquitin ligase Tom1 ubiquitinates excess free r-proteins on sites that are only accessible in their unassembled, unprotected state, thereby channelling its target r-proteins to degradation by the proteasome [10,11,20]. The adaptive RASTR or RPAS stress response is activated by the aggregation of excess free r-proteins, which leads to the sequestration of Ifh1 and chaperones in these aggregates and, thereby, to a downregulation of Ifh1-dependent RPG transcription and the upregulation of Hsf1 target genes, altogether lowering the proteostatic burden by decreasing the amounts of newly synthesized r-proteins and by bolstering the cellular protein folding and degradation capacity [12,13,16]. Third, while the above mechanisms operate to regulate r-protein expression at the transcriptional level and to degrade excess unassembled r-proteins that have accumulated in the nuclear compartment, the two ribosome-associated chaperone systems, the ribosome-associated complex (RAC)-Ssb chaperone triad and the nascent polypeptide-associated complex (NAC) [21–24], cooperate during translation of RPG mRNAs to prevent the aggregation of many newly synthesized r-proteins [25]. Fourth, most r-proteins associate with pre-ribosomal particles in the nucle(ol)us and the high rate of ribosome assembly relies on their rapid nuclear accumulation [26,27]; accordingly, studies in mammalian cells have shown that the nuclear import of r-proteins largely depends, despite their (mostly) small size, on active, importin-mediated transport across the central channel of the nuclear pore complex (NPC) [9,28]. As importins also prevent the aggregation of r-proteins, likely by recognizing and shielding the exposed rRNA-binding regions of free r-proteins, they appear to exhibit a dual function as transport receptors and chaperones [9,29]. In yeast, import of r-proteins has been suggested to be mainly mediated by the importin-β Kap123 [30]. In support of this, selective ribosome profiling revealed that Kap123 has the capacity to co-translationally associate with seven r-proteins, whereas, out of the remaining ten importins (nine importin-βs and the importin-α Kap60/Srp1), only Kap104 exhibited co-translational recognition of one r-protein [31]. However, thorough experimental proof, pinpointing both the nuclear localization signal (NLS) and the responsible importin, for the selective import of a given r-protein by a distinct importin has only been reported in very few cases [32,33], presumably also owing to a partial redundancy among importins, the potential presence of multiple NLSs in an r-protein, and the NLS not necessarily having to correspond to a short linear sequence [30,34,35].

Besides the above-mentioned, mostly general regulation, protection, and import mechanisms, several r-proteins rely on tailor-made solutions in order to be furnished in adequate amounts and in an unproblematic, assembly-competent form. In the case of two r-proteins, Rpl40 (eL40) and Rps31 (eS31), their initial production as fusion proteins (Ubi1/2 and Ubi3) bearing an N-terminal ubiquitin moiety, which is cleaved off prior to their incorporation into pre-ribosomal particles, is necessary to ensure their efficient production [36–40], suggesting that the fused ubiquitin may fulfil the function of a *cis*-acting chaperone. Notably, several r-proteins have been shown to be transiently associated with a selective binding partner; as these ensure, albeit by different means and by employing different folds for specific r-protein recognition, the soluble expression, nuclear transport, and/or efficient assembly of their r-protein clients, they are collectively referred to as dedicated chaperones (DCs) of r-proteins [41,42]. Up to now, eight bona fide DCs that beneficially act on nine different r-proteins have been identified [42–45], with one of these, Syo1, being able to simultaneously interact with the r-proteins Rpl5 (uL18) and Rpl11 (uL5) [46]. By doing so, the transport adaptor Syo1 not only enables the Kap104-mediated co-import of Rpl5 and Rpl11 but also their subsequent assembly with the 5S rRNA into an assembly-competent 5S ribonucleoprotein (RNP) precursor [46–48]. Interestingly, six of the eight DCs, namely Rrb1, Acl4, Syo1, Sqt1, Tsr4, and Yar1, have the capacity to co-translationally capture, almost exclusively by recognizing an N-terminally located binding site, the r-protein Rpl3 (uL3), Rpl4 (uL4), Rpl5, Rpl10 (uL16), Rps2 (uS5), and Rps3 (uS3), respectively [34,41,44,45], indicating that such an early encounter may in many instances be crucial for an optimal protection and/or folding of the r-protein client. In contrast, the DCs Bcp1 and Tsr2 have been suggested to only associate with Rpl23 (uL14) and Rps26 (eS26), respectively, once the r-protein has entered the nucleus, where they would act, by releasing Rpl23 and Rps26 from redundant import receptors and mediating their safe transfer to the pre-ribosomal assembly site, as nuclear escortins [43,49]. Finally, and in line with the necessity for a tight regulation of the abundance of unassembled r-proteins, the expression levels of several r-proteins are fine-tuned through autoregulatory feedback mechanisms. Some r-proteins, when excessively present, bind to their own (pre-)mRNA and thereby, for example, repress its translation, inhibit its splicing, and/or promote its degradation [50–56]. In the case of Rpl3 and Rpl4, their excess presence indirectly downregulates *RPL3* or *RPL4* mRNA levels by reducing the availability of Rrb1 and Acl4 to bind in a timely manner to nascent Rpl3 and Rpl4, respectively, and thereby enabling instead the recruitment of a regulatory machinery consisting of the NAC and the Caf130-associated Ccr4-Not complex [14]. Strikingly, deregulated expression of Rpl3 and/or Rpl4 in cells lacking Tom1 leads to their massive aggregation, a perturbation of overall proteostasis, and lethality, underscoring the importance of the ERISQ pathway and the DCs Rrb1 and Acl4 for protecting cells from the potentially detrimental effects of their surplus production [14].

We have a long-standing interest in understanding how r-proteins safely reach their site of pre-ribosomal incorporation. With the primary aim of identifying novel DCs, we have performed a TurboID-based proximity labelling screen with all *S. cerevisiae* LSU r-proteins. Despite the relatively short 1-h duration of induced r-protein expression and biotinylation, chosen to preferentially capture transient interactions of r-proteins prior to their assembly into pre-ribosomal particles, this approach nevertheless permits, as illustrated for selected examples, a spatiotemporal resolution of r-protein neighbourhoods on successive pre-60S particles. More importantly, and as aimed for, our large-scale proximity labelling screen has proven to be well suited for the detection of transient interactions of unassembled r-proteins. Besides unveiling potential interactions with distinct importins, which we confirmed in the case of four r-proteins by yeast two-hybrid (Y2H) assays, our approach also allowed to uncover established and potential DCs among the enriched proteins. In this study, we have focused on two previously uncharacterized, non-essential proteins, Acl1 (Ycr051w) and Bcl1 (Ynl035c), which could be identified in the proxiOME of the universally conserved Rpl1 (uL1). Our genetic, cell biological, biochemical, and structural data indicate that the ankyrin repeat-containing Acl1 and the predicted WD-repeat β-propeller protein Bcl1 function as cooperating DCs that mediate the safe and efficient transfer of Rpl1 to its assembly site on nucleolar pre-60S subunits. While Acl1 appears to be a fungi-specific protein, proximity labelling with human RPL10A, the orthologous protein of Rpl1, in the human HeLa cell line uncovered WDR89, the likely human orthologue of Bcl1, as an RPL10A-proximal protein, suggesting that Bcl1 carries out an evolutionarily conserved function as a DC of uL1.

## MATERIALS AND METHODS

### Plasmids

All recombinant DNA techniques were according to established procedures using *Escherichia coli* DH5α for cloning and plasmid propagation. All cloned DNA fragments generated by PCR amplification were verified by sequencing. More information on the plasmids, which are listed in the Supplementary file 1, is available upon request.

### Yeast genetics

The *S. cerevisiae* strains used in this study are listed in Supplementary file 1 and are, unless otherwise specified, derived from W303 [57]. Deletion disruption as well as C-terminal tagging at the genomic locus was performed as described [58,59]. The preparation of media, yeast transformation, and genetic manipulations were done according to established procedures.

### TurboID-based proximity labelling assay in yeast

The wild-type strain YDK11-5A [60] was transformed with centromeric plasmids (*LEU2* marker) expressing the TurboID-fused bait proteins under the control of the copper-inducible *CUP1* promoter. Transformed cells were grown in exponential phase to an OD_600_ of around 0.5 at 30°C in 100 mL of synthetic complete medium containing 2% glucose as carbon source and lacking leucine (SC-Leu) that was prepared with copper-free yeast nitrogen base (FORMEDIUM). Expression from the *CUP1* promoter was induced by addition of copper sulphate to a final concentration of 500 μM. Simultaneously, biotin (Sigma-Aldrich), from a freshly prepared 100 mM stock solution in dimethyl sulfoxide, was added to a final concentration of 500 μM. Then, cells were grown for an additional hour, harvested by centrifugation at 4000 rpm for 5 min at 4°C, washed once with 50 mL of ice-cold nanopure water, resuspended in 1 mL of ice-cold lysis buffer (LB: 50 mM Tris-HCl (pH 7.5), 150 mM NaCl, 1.5 mM MgCl_2_, 0.1% SDS, and 1% Triton X-100) containing 1 mM PMSF, transferred to 2 mL safe-lock tubes, and pelleted by centrifugation. Cell pellets were rapidly frozen in liquid nitrogen and stored at – 80°C. Prior to their lysis, cells were resuspended in 400 μL lysis buffer containing 0.5% sodium deoxycholate and 1 mM PMSF (LB-P/D). After addition of glass beads, cells were lysed with a Precellys 24 homogenizer (Bertin Technologies) set at 5000 rpm for three 30 s lysis cycles alternating with 30 s breaks at 4°C. The resulting lysates were transferred to a 1.5 mL tube. For complete extract recovery, 200 μL LB-P/D were added to the glass beads and, after brief vortexing, combined with the already transferred lysate. Cell lysates were centrifuged for 10 min at 13,500 rpm at 4°C and supernatants were transferred to a new 1.5 mL tube. Total protein concentration was determined with the Pierce™ BCA Protein Assay Kit (Thermo Scientific) using a BioTek 800 TS microplate reader. 2 mg of total protein in a final volume of 800 μL of LB-P/D was added to the equivalent of 100 μL of Pierce^TM^ High-Capacity Streptavidin Agarose Resin (Thermo Scientific) slurry (corresponding to 50 μL of settled beads), which had been blocked by incubation with 1 mL LB containing 3% BSA for 1 h at RT and washed four times with 1 mL LB. For affinity purification of biotinylated proteins, lysates were incubated for 1 h on a rotating wheel at RT. Then, beads were washed once with 1 mL of wash buffer (50 mM Tris-HCl (pH 7.5), 2% SDS), five times with 1 mL LB, and five times with 1 mL ABC buffer (100 mM ammonium bicarbonate (pH 8.2)). Purified proteins were eluted from the beads by addition of 30 μL 3x SDS sample buffer containing 10 mM biotin and 20 mM dithiothreitol (DTT) and incubation for 10 min at 75°C. This step was repeated a second time, and the two eluates were combined and stored at –20°C. Upon reduction (incubation for 10 min at 75°C with an additional 1 μL of 100 mM DTT in a thermoshaker set at 1000 rpm) and alkylation (10-min incubation in the dark at RT after addition of 1 μL 550 mM iodoacetamide (IAA) in a thermoshaker set at 1000 rpm), the entire eluates were separated on NuPAGE 4-12 % Bis-Tris gels (Invitrogen) in NuPAGE 1x MES SDS running buffer (Novex) for a total of 12 min at 200 V. Then, the gels were incubated in fixing solution (40% methanol, 7% acetic acid) for 1 h, stained by a 5-min incubation in Brilliant Blue G Colloidal Coomassie (Sigma-Aldrich), rinsed for 5 min in rinsing solution (25% methanol, 5% acetic acid), destained three consecutive times for 40 min with destaining solution (25% methanol), and washed for 3x 5 min in nanopure water. Each lane was cut from the slot to the migration front into three gel pieces of equal lengths (fractions), which were cut into smaller pieces and transferred to low protein-binding 1.5 mL tubes (Sarstedt). From this point on, each fraction was individually treated with HPLC grade solvents or solutions prepared therein. The fractions were washed three times by alternating 10-min incubations at RT in 100-150 μL ABC buffer and 100-150 μL ethanol in a thermoshaker set at 1000 rpm. For in-gel digestion of proteins, gel pieces were covered with 120 μL ABC buffer containing 1 μg Sequencing Grade Modified Trypsin (Promega) and incubated overnight at 37°C with shaking at 1000 rpm. The proteolytic digestion was quenched by addition of 50 μL 2% trifluoroacetic acid (TFA) and incubation for 10 min at RT in a thermoshaker set at 1000 rpm, and the supernatant was transferred into a new tube. The gel pieces were then incubated, again for 10 min at RT with shaking at 1000 rpm, another two times with 100-150 μL ethanol, and these two supernatants were combined with the first supernatant. The organic solvent of the combined supernatants was evaporated by using a SpeedVac. Then, 200 μL of buffer A (0.1% formic acid) was added to the remaining volume, and the samples were centrifuged for 10 min at 13,000 rpm and applied to C18 StageTips [61], which were beforehand washed with 50 μL of buffer B (80% acetonitrile, 0.1% formic acid) and equilibrated twice with 50 μL of buffer A, for desalting and peptide purification. After loading, the StageTips were washed once with 100 μL of buffer A, and the peptides were eluted with 50 μL of buffer B. The elution solvents were completely evaporated using a SpeedVac, and the peptides were resuspended by first adding 3 μL of buffer A* (3% acetonitrile, 0.3% TFA) and then, after brief vortexing, 17 μL of buffer A*/A (30% buffer A*/70% buffer A). The samples were vortexed, quickly centrifuged, and stored at –80°C.

### Mass spectrometry-based proteomic analysis

LC-MS/MS measurements were performed on a Q Exactive HF-X (Thermo Scientific) coupled to an EASY-nLC 1200 nanoflow-HPLC (Thermo Scientific). HPLC-column tips (fused silica) with 75 μm inner diameter were self-packed with ReproSil-Pur 120 C18-AQ, 1.9 μm particle size (Dr. Maisch GmbH) to a length of 20 cm. Samples were directly applied onto the column without a pre-column. A gradient of A (0.1% formic acid in H_2_O) and B (0.1% formic acid in 80% acetonitrile in H_2_O) with increasing organic proportion was used for peptide separation (loading of sample with 0% B; separation ramp: from 5-30% B within 85 min). The flow rate for sample application was 600 nL/min and for sample separation 250 nL/min. The mass spectrometer was operated in the data-dependent mode and switched automatically between MS (max. of 1 x 10^6^ ions) and MS/MS. Each MS scan was followed by a maximum of ten MS/MS scans using a normalized collision energy of 25% and a target value of 1000. Parent ions with a charge state form *z* = 1 and unassigned charge states were excluded for fragmentation. The mass range for MS was *m/z* = 370-1750. The resolution for MS was set to 70,000 and for MS/MS to 17,500. MS parameters were as follows: spray voltage 2.3 kV, no sheath and auxiliary gas flow, ion-transfer tube temperature 250°C.

The MS raw data files were analysed with the MaxQuant software package version 1.6.2.10 [62] for peak detection, generation of peak lists of mass-error-corrected peptides, and database searches. The UniProt *Saccharomyces cerevisiae* database (version March 2016), additionally including common contaminants, trypsin, TurboID, and GFP, was used as reference. Carbamidomethylcysteine was set as fixed modification and protein amino-terminal acetylation, oxidation of methionine, and biotin were set as variable modifications. Four missed cleavages were allowed, enzyme specificity was Trypsin/P, and the MS/MS tolerance was set to 20 ppm. Peptide lists were further used by MaxQuant to identify and relatively quantify proteins using the following parameters: peptide and protein false discovery rates, based on a forward-reverse database, were set to 0.01, minimum peptide length was set to seven, and minimum number of unique peptides for identification and quantification of proteins was set to one. The ‘match-between-run’ option (0.7 min) was used.

For quantification, missing iBAQ (intensity-based absolute quantification) values in the two control purifications from cells expressing either the GFP-TurboID or the NLS-GFP-TurboID (controls for C-terminally TurboID-tagged bait proteins) or the TurboID-GFP or the NLS-TurboID-GFP (controls for N-terminally TurboID-tagged bait proteins) were imputed in Perseus using standard settings [63]. For normalization of intensities in each independent purification, iBAQ values were divided by the median iBAQ value, derived from all nonzero values, of the respective purification. To calculate the enrichment of a given protein compared to its average abundance in the two control purifications, the normalized iBAQ values were log2-transformed and those of the control purifications were subtracted from the ones of each respective bait purification. In the case of the TurboID series with the DC baits, the enrichment of a given protein was calculated compared to its median abundance in the two control purifications and the remaining DC purifications, except for the Acl1 and Bcl1 TurboID assays where these two were reciprocally excluded. For graphical presentation, the normalized iBAQ value (log10 scale) of each protein detected in a bait purification was plotted against its log2-transformed enrichment compared to the control purifications (fold change). For the bar graph representation, the log2-transformed enrichment of a given protein in the indicated TurboID assays is represented by bars, while its median normalized log2-transformed enrichment across the different TurboID assays is shown as a dotted line.

### MiniTurbo-based proximity labelling in HeLa cells

The DNA sequence coding for the *Homo sapiens* RPL10A protein was PCR-amplified from plasmid pADH111-HsRPL10A (pDK10427), generated by cloning the PCR-amplified *RPL10A* coding sequence (template pNTI194 (Addgene plasmid #84266)) into the *Nde*I/*Bam*HI-restricted plasmid pADH111-LTV1 (pDK3331), and cloned by Gibson assembly (NEB, M5510A) between the *Nhe*I and *Pst*I restriction sites of the lentiviral donor vector pSKP-32, a pCW57.1-derived plasmid bearing the MND-Blasticidin resistance cassette instead of the hPGK-Puromycin resistance cassette [64], to generate plasmid pDS79 containing the *RPL10A* gene under the transcriptional control of a doxycycline-inducible promoter and fused at its 3’ end to sequences encoding the V5 tag, the miniTurbo (MT) biotin ligase, and the HA tag. As controls, pSKP-32 plasmids encoding HA-MT-V5-EGFP (pDS48) and EGFP-V5-MT-HA-NLS (pDS82), respectively, were constructed. Replication-defective lentiviral particles were produced as previously described [65] by co-transfecting, using the JetPRIME transfection reagent (Polyplus, 114-75), the above lentiviral donor plasmids with packaging and envelope expressing plasmids psPAX2 (Addgene plasmid #12260) and pMD2.G (Addgene plasmid #12259) into HEK293T cells seeded the night before. Transfection medium was changed 12 h post transfection, and lentiviral supernatants were harvested 24 h later, filter-sterilized through 0.2 μm syringe filters, supplemented with 8 μg/mL polybrene (Sigma-Aldrich, H9268), and stored in aliquots at –80°C. These lentiviral particles expressing the different fusion constructs were, at different viral dilutions (1:2 to 1:100) prepared in Dulbecco’s Modified Eagle Medium (DMEM, PAN-Biotech, P04-04510) containing 8 μg/mL polybrene, used to infect HeLa cells (ATCC CCL-2), which had been validated by genotyping (Microsynth) and negatively tested for mycoplasma. HeLa cells were grown in a humidified incubator in DMEM supplemented with 10% Fetal Bovine Serum (FBS; Biowest, S181B-500) and 1% penicillin-streptomycin (PAN-Biotech, P06-07100) at 37°C and 5% CO_2_. Infected cells were selected 24 h post infection in medium (DMEM, 10% FBS, and 1% penicillin-streptomycin) containing 4 μg/mL Blasticidin (InvivoGen, ant-bl-1) until selection was complete. Selected cells were tested by Western blotting using anti-HA antibodies (mouse monoclonal anti-HA antibody (6E2); Cell Signaling, #2367) to assess expression of the HA-tagged fusion proteins and to determine the optimal viral dilution. For the proximity labelling experiment, HeLa cells (two 15 cm dishes per replicate) were infected in three experimental replicates with the lentiviral particles expressing the RPL10A-V5-MT-HA fusion protein and the HA-MT-V5-EGFP and EGFP-V5-MT-HA-NLS control fusion proteins. Expression was induced with 2 mg/mL doxycycline for 24 h and biotinylation by incubation in the presence of 400 μM biotin for 90 min. Then, cells were washed five times with ice-cold PBS, scraped, harvested by centrifugation at 3000 rpm for 2 min, snap frozen in liquid nitrogen, and stored at –80°C. Cell pellets were lysed in 1.2 mL of modified RIPA buffer (50 mM Tris-HCl (pH 7.5), 150 mM NaCl, 1 mM EDTA, 1 mM EGTA, 1% Triton X-100, 0.5% sodium deoxycholate, and 0.1% SDS), containing Benzonase (1:5000 dilution), cOmplete^TM^, EDTA-free Protease Inhibitor Cocktail (Roche, 05-056-489-001), and 1 mM PMSF, by a 30-min incubation on ice with periodical vortexing (four times 15 s). Then, cell lysates were clarified by centrifugation at 14,000 rpm for 10 min at 4°C and transferred to a new tube. Total protein concentration in the clarified cell extracts was determined with the Pierce^TM^ BCA Protein Assay Kit (Thermo Scientific, 23225) using a microplate reader (BioTek 800 TS). For affinity purification of biotinylated proteins, 100 μL of Pierce^TM^ High Capacity Streptavidin Agarose Resin (Thermo Scientific, 20361) slurry, corresponding to 50 μL of settled beads, were washed twice with modified RIPA buffer L (lacking sodium deoxycholate, SDS, protease inhibitors, and PMSF), and equal amounts of proteins (3.5 mg) in a total volume of 1.3 mL were incubated with the washed beads for 90 min on a rotating wheel at RT. After binding, beads were washed five times with the modified RIPA buffer, three times with a HEPES buffer (50 mM HEPES-KOH (pH 8), 100 mM KCl, 2 mM EDTA, 0.1% Triton X-100, and 10% glycerol), and two times with ABC buffer (100 mM ammonium bicarbonate (pH 8.2)).

Bound proteins were processed by a filter-aided sample preparation (FASP) protocol [66]. After the last wash, biotinylated proteins bound to the streptavidin beads were denatured with 8 M urea in a 50 mM Tris-HCl (pH 8) buffer containing 1 mM DTT for 5 min at RT. The denatured proteins and the beads were transferred to a 10 kDa molecular weight cut-off filter centrifugal unit (Vivacon 500, 10,000 MWCO; Sartorius, VN01H02), followed by reduction with 1 mM DTT and alkylation with 5.5 mM iodoacetamide for 10 min in the dark at RT. The samples were centrifuged at 10,000 x g for 30 min and then washed three times with 50 mM Tris-HCl (pH 8) buffer with the same centrifugation conditions. On-filter digestion of proteins was performed overnight at 37°C in a thermoshaker set to 550 rpm in 200 μL 50 mM Tris-HCl (pH 8) buffer containing 1.5 μg Sequencing Grade Modified Trypsin (Promega, V5113). Tryptic peptides were eluted by centrifugation, purified over C18 StageTips, and analysed by LC-MS/MS as described above.

The MS raw data files were analysed with the MaxQuant software package version 1.6.2.10 [62], using the UniProt *Homo sapiens* database (version 2016), additionally including common contaminants, trypsin, miniTurbo, and GFP, as reference. Perseus (software version 1.6.2.3) [63] was used for further data analysis. For quantification, missing iBAQ values in the control purifications from cells expressing either the HA-MT-V5-EGFP or the EGFP-V5-MT-HA-NLS bait were imputed in Perseus.

For normalization of intensities in each independent purification, iBAQ values were divided by the median iBAQ value, derived from all nonzero values, of the respective purification. Then, the average of the median-normalized iBAQ values obtained for the three experimental RPL10A bait replicates, excluding the missing values (Not a Number (NaN) values), and the six experimental control replicates were calculated. To calculate the enrichment of a given protein compared to its average abundance in the control purifications, the normalized, averaged iBAQ values were log2 transformed and those of the control purifications were subtracted from the ones of the RPL10A bait purification. For graphical representation, the normalized, averaged iBAQ value (log10 scale) of each protein detected in the RPL10A bait purification was plotted against its log2-transformed enrichment compared to the control purifications (fold change).

### Yeast two-hybrid (Y2H) interaction assay

The Y2H reporter strain PJ69-4A [67] was co-transformed with two plasmids expressing the bait protein fused to the Gal4 DNA-binding domain (G4BD, *TRP1*) and the prey protein fused to the Gal4 activation domain (G4AD, *LEU2*). Transformed cells were selected on synthetic complete plates lacking leucine and tryptophan (SC-LT). The protein-protein interactions were documented by spotting representative transformants in 10-fold serial dilution steps onto SC-LT, SC-His-Leu-Trp (SC-HLT, *HIS3* reporter), and SC-Ade-Leu-Trp (SC-ALT, *ADE2* reporter) plates, which were incubated for 3 days at 30°C. Growth on SC-HLT plates is indicative of a weak/moderate interaction, whereas only relatively strong interactions also permit growth on SC-ALT plates.

### In vitro binding assays

For in vitro binding assays between binary combinations of full-length or truncated variants of Acl1, Bcl1, and Rpl1, the two proteins were co-expressed from pETDuet-1 (Novagen) in Rosetta(DE3) (Novagen) *E. coli* cells. Cells were grown in 200 mL of lysogeny broth (LB) medium containing ampicillin and chloramphenicol at 37°C and protein expression was induced at an OD_600_ of around 0.5 to 0.8 by the addition of IPTG to a final concentration of 0.5 mM. After 5 h of growth at 23°C, cells were harvested and stored at –80°C. Cells were resuspended in 25 mL lysis buffer (50 mM Tris-HCl (pH 7.5), 200 mM NaCl, 1.5 mM MgCl_2_, 5% glycerol) and lysed with a M-110L Microfluidizer (Microfluidics). The lysate (30 mL volume) was adjusted by the addition of 300 μL 10% NP-40 to 0.1% NP-40 (note that from here onwards all buffers contained 0.1% NP-40). An aliquot of 50 μL of total extract (sample T) was taken and mixed with 50 μL of 6x SDS sample buffer. The total extract was then centrifuged at 4°C for 20 min at 14,000 rpm. The soluble extract was transferred to a 50 mL Falcon tube and, as above, an aliquot of 50 μL of soluble extract (sample S) was taken and mixed with 50 μL of 6x SDS sample buffer. The insoluble pellet fraction (sample P) was resuspended in 3 mL of lysis buffer and 5 μL thereof were mixed with 45 μL of lysis buffer and 50 μL of 6x SDS sample buffer. The soluble extract (30 mL) was adjusted to 15 mM imidazole by adding 180 μL 2.5 M imidazole pH 8.

Upon addition of 250 μL of Ni-NTA Agarose slurry (Qiagen), samples were incubated for 2 h on a turning wheel at 4°C. Then, the Ni-NTA Agarose beads were pelleted by centrifugation at 4°C for 2 min at 1800 rpm, resuspended in 2 mL of lysis buffer, and transferred to a 2 mL Eppendorf tube. The Ni-NTA Agarose beads were first washed five times with 1 mL of lysis buffer containing 15 mM imidazole and then two times for 5 min, by rotation on a turning wheel at 4°C, with 1 mL lysis buffer containing 50 mM imidazole. Elution of bound proteins was carried out by incubation of the Ni-NTA Agarose beads with 1 mL of lysis buffer containing 500 mM imidazole for 5 min on a turning wheel at 4°C. The eluate (sample E) was transferred to a 1.5 mL Eppendorf tube and 50 μL thereof were mixed with 50 μL of 6x SDS sample buffer. Protein samples (5 μL of samples T, P, S, and E) were separated on Bolt 4-12% Bis-Tris Plus 15-well gels (Invitrogen), run in Bolt 1x MES SDS running buffer, and subsequently stained with Brilliant Blue G Colloidal Coomassie (Sigma-Aldrich). For Western analysis, appropriate dilutions of the above samples were separated on Bolt 4-12% Bis-Tris Plus 15-well gels, run in 1x MES SDS running buffer, and proteins were subsequently blotted onto nitrocellulose membranes (GE Healthcare).

For purification of the trimeric Acl1-Rpl1-Bcl1 complex, Rpl1 and either C-terminally Flag-tagged Acl1 or untagged Acl1 (negative control purification) were co-expressed from pETDuet-1 and C-terminally (His)_6_-tagged Bcl1.N366 was expressed from pET-24d. Rosetta(DE3) cells were co-transformed with the two plasmids and grown in 600 mL of LB medium containing ampicillin, kanamycin, and chloramphenicol. The co-expressed proteins were purified by a two-step affinity purification approach, first via Ni-NTA and then via anti-FLAG agarose beads. The Ni-NTA purification was done as described above, but bound proteins were eluted two consecutive times with 500 μL of lysis buffer containing 500 mM imidazole for 5 min on a turning wheel at 4°C. An aliquot of 50 μL of the first imidazole eluate (sample E1) was mixed with 50 μL of 6x SDS sample buffer. Then, the two eluates were combined and incubated for 2 h with 30 μL of Anti-FLAG M2 Affinity Gel slurry (Sigma-Aldrich). Subsequently, the bead-containing solution was transferred to Mobicol “Classic” columns (Mo Bi Tec), and beads were washed three consecutive times with 1 mL of lysis buffer. Bound proteins were eluted with 50 μL of lysis buffer containing 454 μg/mL 1xFLAG peptide (Sigma-Aldrich) for 30 min on a turning wheel at 4°C. The entire FLAG eluate (∼50 μL) was mixed with 50 μL of 6x SDS sample buffer (sample E2). 5 μL of eluate samples E1 and E2 were, as described above, separated on gels and analysed by Coomassie staining and Western blotting.

### Fluorescence microscopy

To assess their subcellular localization, proteins were tagged with yeast codon-optimized mNeonGreen (referred to as ymNeonGreen or ymNG) ([68]; Allele Biotechnology) or a single (yEGFP) or triple yeast-enhanced GFP (3xyEGFP) [69] and expressed, as indicated in the figure legends, either from the genomic locus or from plasmids under the control of their cognate promoter or the *ADH1* promoter. The nucleolar marker protein Nop58-yEmCherry was either expressed from its genomic locus or from plasmid under the control of its cognate promoter. Transformed cells were grown in the indicated liquid media at 30°C and inspected in exponential growth phase. Live yeast cells were imaged by fluorescence microscopy using a VisiScope CSU-W1 spinning disk confocal microscope (Visitron Systems GmbH). The ImageJ software was used to process the images.

### Sucrose gradient analysis and fractionation

Cell extracts for polysome profile analyses were prepared as previously described [70] and eight A_260_ units were layered onto 10-50% sucrose gradients that were centrifuged at 39,000 rpm in a TH-641 swinging bucket rotor (Thermo Scientific) at 4°C for 2 h 45 min. Sucrose gradients were analysed using an ISCO UA-6 system with continuous monitoring at A_254_. For the fractionation experiments, five A_260_ units were subjected to sucrose gradient centrifugation and 20 fractions of around 500 μL were collected and processed as described [60]. Precipitated proteins were resuspended in 50 μL 3x SDS sample buffer and 5 μL each fraction was separated on NuPAGE 4-12% Bis-Tris 26-well gels (Novex), run in 1x MES running buffer, and subsequently analysed by Western blotting. As an input control, 0.05 A_260_ units of total cell extract was run alongside the fractions.

### Preparation of total yeast protein extracts and Western analysis

Total yeast protein extracts were prepared as previously described [71]. Western blot analysis was carried out according to standard protocols. The following primary antibodies were used in this study: anti-Rpl1 (1:5000; obtained from the laboratory of J. de la Cruz, University of Sevilla, Sevilla, Spain [72]), anti-Rpl4 (1:10,000; L. Lindahl, University of Baltimore, Baltimore, USA), anti-Rps3 (1:15,000; M. Seedorf [73]), mouse monoclonal anti-FLAG (1:2000 – 1:10,000; Sigma), anti-His_6_ (1:500; Roche). Secondary goat anti-mouse or anti-rabbit horseradish peroxidase-conjugated antibodies (Bio-Rad) were used at a dilution of 1:10,000. For detection of TAP-tagged proteins, the peroxidase anti-peroxidase (PAP) Soluble Complex antibody produced in rabbit (Sigma-Aldrich) was used at a dilution of 1:20,000. Immobilized protein-antibody complexes were visualized by using enhanced chemiluminescence detection kits (WesternBright Quantum and Sirius; Advansta) and a Fusion FX7 Edge imaging system (Vilber). Images were processed with ImageJ [74] or the Evolution-Capt Software (Vilber). Intensities of Rpl1 and Rpl4 signals in sucrose gradient fractions were quantified using the Evolution-Capt Software for the two experimental replicates (n=2). The normalized abundance of Rpl1 and Rpl4 in the individual fractions was calculated by dividing their signal intensity in each fraction by their average signal intensity in two polysomal fractions (fractions 13 and 14). The normalized Rpl1/Rpl4 ratios of each fraction were calculated and plotted on the y-axis against the corresponding fraction number on the x-axis.

### Protein expression and purification for X-ray crystallography

For the expression and purification of the Rpl1(63-158)-Acl1.N127 complex, *E. coli* BL21(DE3) cells (Novagen) were transformed with pDK9667 (pETDuet-1/RPL1B(63-158)-(His)_6_--ACL1.N127) and plated on LB agar containing 100 μg/mL ampicillin. Following overnight incubation at 37°C, expression cultures were inoculated directly from the plate. Protein production was carried out in LB medium containing 100 μg/mL ampicillin under auto-induction conditions (1% (w/v) D(+)-lactose-monohydrate). The cultures were incubated for 16 h at 30°C with shaking at 180 rpm. Cells were harvested by centrifugation (4000 rpm, 15 min, 4°C; Fiberlite F9-6 x 1000 LEX fixed angle rotor (Thermo Scientific)) using a Sorvall LYNX 6000 centrifuge (Thermo Scientific) and resuspended in 20 mL of buffer A (20 mM HEPES (pH 8.0), 250 mM NaCl, 20 mM MgCl₂, 20 mM KCl, 40 mM imidazole). Cell lysis was performed using a M-110L Microfluidizer (Microfluidics) and the lysate was clarified by high-speed centrifugation (20,000 rpm, 20 min, 4°C; LYNX A27-8 x 50 fixed angle rotor (Thermo Scientific)) using a Sorvall LYNX 6000 centrifuge. Protein purification was then accomplished in a two-step process via immobilized metal ion affinity chromatography (IMAC) with nickel-nitrilotriacetic acid (Ni-NTA) followed by size exclusion chromatography (SEC): The clarified lysate was first applied to a 5 mL HisTrap Fast Flow column (Cytiva). Immobilized target proteins were washed with ten column volumes (CV) of buffer A and eluted with seven CV of buffer B (20 mM HEPES (pH 8.0), 250 mM NaCl, 20 mM MgCl₂, 20 mM KCl, 500 mM imidazole). The Rpl1(63-158)-Acl1.N127 complex was then further purified by SEC using a HiLoad 16/600 Superdex 200 column (Cytiva) equilibrated with SEC buffer (20 mM HEPES (pH 7.5), 200 mM NaCl, 20 mM MgCl₂, 20 mM KCl). Finally, peak fractions containing the Rpl1(63-158)-Acl1.N127 complex, as verified by Coomassie-stained SDS-PAGE, were pooled and concentrated.

### Crystallization and structure determination

Crystallization screens were performed by the sitting drop method in MRC 2-well crystallization plates (SWISSCI) at 20°C, with 0.5 μL drops composed of protein and precipitant solutions mixed in 1:1 and 1:2 ratios. Precipitation solutions were acquired commercially (NeXtal JCSG Core Suite I-IV, each comprising 96 crystallization conditions). The Rpl1(63-158)-Acl1.N127 complex crystallized at a concentration of 39 mg/mL within four weeks in 0.2 mM ammonium acetate, 0.1 mM sodium citrate (pH 5.6), and 30% (w/v) PEG 4000.

Prior to data collection, protein crystals were flash-frozen in liquid nitrogen using a cryo-solution consisting of the mother liquor supplemented with 20% (v/v) glycerol. Diffraction data was collected at 100 K on beamline ID30A-3 at the European Synchrotron Radiation Facility (ESRF; Grenoble, France). X-ray data was processed, via the XDSAPP3 interface [75], with *XDS* [76]. For determining the final experimental structure of the Rpl1(63-158)-Acl1.N127 complex, initial phases were obtained by molecular replacement using the *phenix.phaser* module of the Phenix suite [77] with an AlphaFold3 model [78] of the Rpl1(63-158)-Acl1.N127 complex as a search model (TFZ = 23.7; LLG = 399). Following *phenix.autobuild*, the structure was finalized through iterative cycles of model building in *Coot* [79] and refinement with *phenix.refine*. Data collection and refinement statistics are summarized in Supplementary Table S1. Coordinates are deposited in the Protein Data Bank (PDB) under the accession code 9T3L.

### Analysis of experimental and predicted structures

Analysis and image preparation of three-dimensional structures, downloaded from the PDB archive or predicted with AlphaFold-Multimer or AlphaFold3 [78,80,81], were carried out with the ChimeraX [82,83] software.

### Identification of orthologous proteins and sequence alignments

Orthologous proteins of *S. cerevisiae* Acl1 (UniProt: P25631) were found using the default settings of the BLAST function of the UniProtKB [84], which also indicates the phylogenetic tree (e.g., kingdom, subkingdom, phylum, and subphylum) of the identified orthologous proteins. Multiple sequence alignments of orthologous proteins were independently generated for subkingdoms, phyla, or subphyla in the ClustalW output format with T-Coffee using the default settings of the EBI website interface [85]. According to the Alliance of Genome Resources (https://www.alliancegenome.org; Version: 8.2.0), *H. sapiens* WDR89 (UniProt: Q96FK6) is the orthologous protein of *S. cerevisiae* Bcl1 (UniProt: P53962).

### Determination of co-translational capturing by qRT-PCR

Co-translational association of Acl1-TAP, Bcl1-TAP, TAP-Bcp1, Bcp1-TAP, TAP-Rrb1, and Acl4-TAP with nascent r-proteins was assessed by IgG-Sepharose pull-down and real-time quantitative reverse transcription PCR (real-time qRT-PCR) as previously described [41,44] with the following modifications: (i) after the last wash, the beads were resuspended in 200 μL of lysis buffer-PC containing 1 mM of DTT; then, 2 μL of RiboLock and 5 μL of home-made TEV protease were added and the samples were incubated overnight on a rotating wheel at 4°C; (ii) for RNA isolation, 50 μL of total cell extracts were diluted with 150 μL of lysis buffer, and RNA was purified from total extracts and TEV eluates using the Quick-RNA Microprep Kit (Zymo Research) and eluted in 15 μL of RNase/DNase free water; then, purified total RNAs were diluted to 5 ng/μL and the volume of the TEV-eluate RNA extracts was increased to 120 μL.

The following oligonucleotide pairs were used for the specific amplification of DNA fragments, corresponding to the *RPL1A*/*B*, *RPL3*, *RPL4A*/*B*, and *RPL23A*/*B* mRNAs, from the input cDNAs: RPL1-forward: 5’-ACCCCAGTTTCTCACAACGA-3’, RPL1-reverse: 5’-GCAACAGCCAAACACAAGA-3’ (amplicon size 98 base pairs (bp)), RPL3-forward: 5’-ACTCCACCAGTTGTCGTTGTTGGT-3’, RPL3-reverse: 5’-TGTTCAGCCCAGACGGTGGTC-3’ (amplicon size 86 bp), RPL4-forward: 5’-ACCTCCGCTGAATCCTGGGGT-3’, RPL4-reverse: 5’-ACCGGTACCACCACCACCAA-3’ (amplicon size 72 bp), RPL23-forward: 5’-AGGGTAAGCCAGAATTGAGAAA-3’, RPL23-reverse: 5’-ACACCGTCTCTTCTTCTCCAAG-3’ (amplicon size 82 bp).

## RESULTS

### Establishing the proxiOME of LSU r-proteins by TurboID-based proximity labelling

DCs transiently interact with free r-proteins before these get incorporated into pre-ribosomal subunits and then become integral components of the long-lived mature ribosomal subunits. Accordingly, it has turned out to be challenging and laborious to efficiently identify DCs in classical affinity purifications of their tagged r-protein clients [34,44,86]. With the aim of facilitating the identification of new potential DCs, we considered harnessing the beneficial characteristics of the TurboID-based proximity labelling approach (Fig. 1A). TurboID is a promiscuous, highly efficient biotin protein ligase derived from the *E. coli* BirA protein [87]. Biotin protein ligases convert biotin into a reactive biotinoyl-5’-AMP (bioAMP) intermediate, which is bound with lower affinity by promiscuous variants, such as BioID or TurboID, than by wild-type BirA [87–89], thereby leading to the formation of a bioAMP concentration gradient around these enzyme variants [90]. The released bioAMP then reacts with surface-exposed primary amines, i.e., mostly the ε-amino group of lysine residues, of proximal proteins; thus, not only leading to the covalent biotinylation of the enzyme-fused bait protein but also of neighbouring proteins that are situated within a certain radius, up to ∼10 nm in the case of BioID [91], around the promiscuous biotin protein ligase [90]. However, the actual labelling radius depends on a number of different parameters, such as the used biotin protein ligase variant and its activity in a given subcellular compartment, the labelling time, the intracellular concentration of biotin, the half-life of the reactive bioAMP, and the duration of the proximity between the enzyme-fused bait protein and its protein neighbourhood(s). Compared to the first-generation BioID variant, which requires long labelling times (up to 24 h) and shows optimal activity at 37°C [87,89,92], TurboID exhibits much faster labelling kinetics, yielding almost the same extent of protein biotinylation in 10 min as BioID in 18 h after biotin addition [87]. Importantly, TurboID also displays high activity at 30°C; thus, permitting to successfully implement the TurboID-based proximity labelling approach in yeast [87,93–95]. After the in vivo proximity labelling and cell lysis, the biotinylated proteins, i.e., the enzyme-fused bait protein and its proximal proteins as well as any non-specifically labelled proteins (background of non-proximal biotinylated proteins that is specific for the cellular compartment(s) where the enzyme-fused bait protein is present) and the four endogenously Bpl1-biotinylated proteins (Acc1/Hfa1, Arc1, Dur12, and Pyc1/Pyc2 [95,96]), can be efficiently purified with streptavidin-coupled beads under stringent conditions and, subsequently, identified and quantified by liquid chromatography-tandem mass spectrometry (LC-MS/MS) (Fig. 1A; see Materials and Methods). As our main goal was to identify new DCs, which are expected to only transiently interact with newly synthesized r-proteins until these get incorporated into pre-ribosomal particles [42], we decided to express the TurboID-fused r-proteins from a centromeric plasmid under the control of the copper-inducible *CUP1* promoter in exponentially growing wild-type cells for 1 h. In this setting, expression and biotinylation can be simultaneously induced by simply adding copper and biotin to the culture medium. To minimize as much as possible persistent biotinylation arising prior induction from the continuous low-level expression of the TurboID-tagged bait r-proteins, due to basal activation of the *CUP1* promoter by low copper concentrations in the standard culture media, and from the utilization of endogenous biotin, which could lead to a disproportionately high labelling of the long-lived r-protein neighbourhoods on mature 60S subunits, we cultured cells in synthetic complete medium prepared with copper-free yeast nitrogen base.

**Figure 1.**
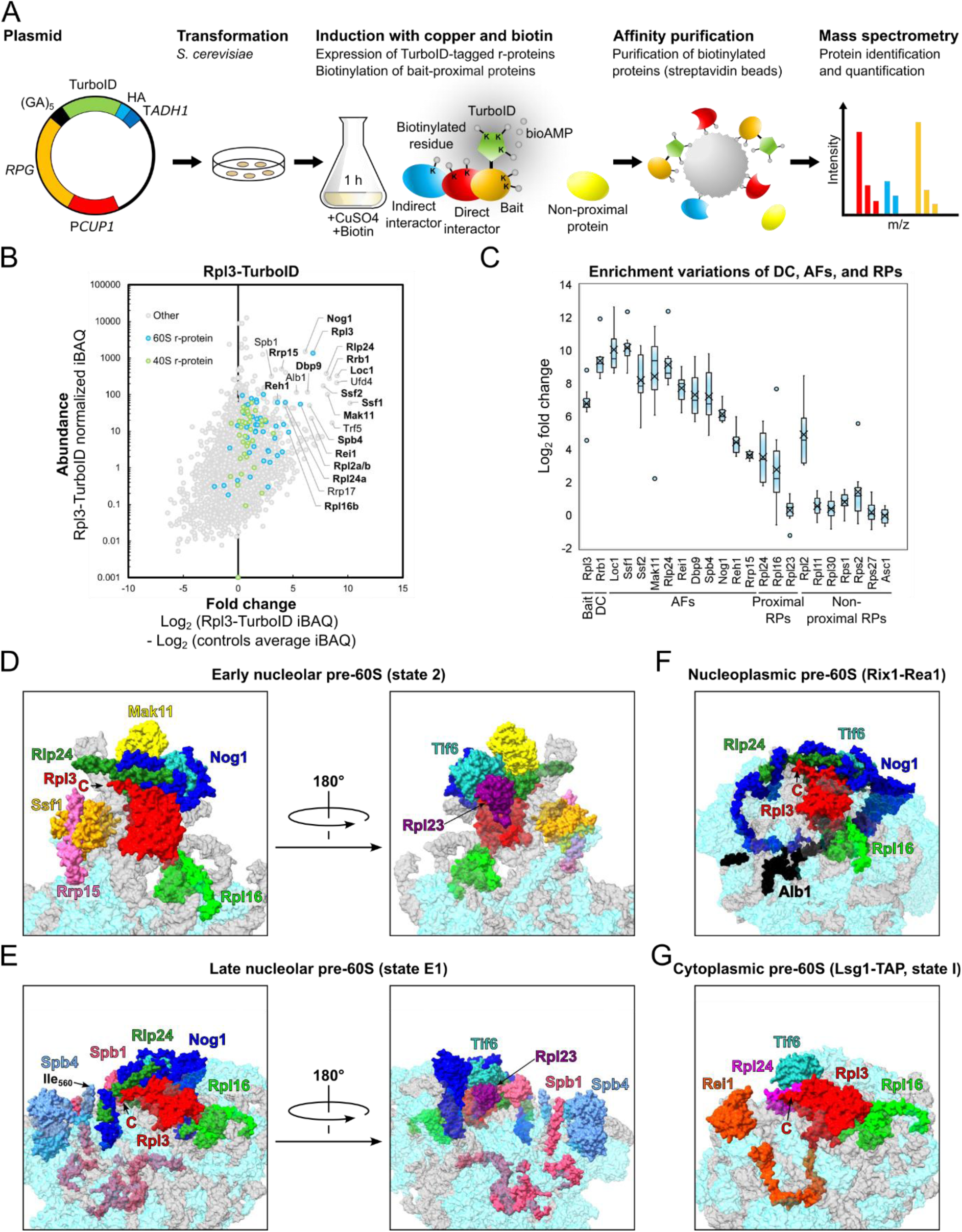
Establishing the proxiOME of LSU r-proteins by TurboID-based proximity labelling. (**A**) Schematic overview of our TurboID-based proximity labelling approach. Wild-type yeast cells (*S. cerevisiae*) are transformed with a centromeric plasmid (*LEU2* marker) containing an r-protein gene (*RPG*, coloured in orange) fused in frame to the DNA sequence encoding the TurboID protein biotin ligase (TurboID, green), which is expressed under the transcriptional control of the copper-inducible *CUP1* promoter (P*CUP1*, red). The C-terminal TurboID tag is separated from the r-protein by inclusion of a (Gly-Ala)_5_ linker (black) and additionally contains a 2xHA tag (blue) at its C-terminal end. For transcription termination, the coding sequence is followed by the *ADH1* terminator (T*ADH1*, dark blue). Expression of the TurboID-tagged r-protein and biotinylation are induced for 1 h by the simultaneous addition of copper and biotin to exponentially growing cultures. TurboID catalyses the conversion of biotin into a reactive biotinoyl-5’-AMP (bioAMP) intermediate, which can covalently modify surface-exposed lysine (K) residues of the TurboID moiety and the bait r-protein as well as of proximal proteins, such as direct (red) and indirect (blue) interaction partners of the TurboID-tagged bait r-protein (orange). Biotinylated proteins are purified from total cell extracts by affinity purification with streptavidin-coupled beads. Purified proteins are then identified and quantified by bottom-up label-free mass spectrometry. **(B)** TurboID results obtained with C-terminally TurboID-tagged Rpl3 (Rpl3-TurboID). The abundance of each detected protein is displayed as its normalized intensity-based absolute quantification (iBAQ) value on the vertical axis, and its relative enrichment (fold change) on the horizontal axis as the log2 fold change of its normalized iBAQ abundance between the bait sample and the average of the GFP-TurboID and NLS-GFP-TurboID controls, from which missing values were imputed according to the normal distribution of the intensities within a sample. The bait r-protein, its known DC, and selected enriched proteins are written in bold. **(C)** Representation of the variation of the enrichment of the Rpl3 bait, its DC Rrb1, proximal AFs, and abundantly detected r-proteins between the experimental replicates (n=8); r-proteins that are close to or far away from Rpl3, as indicated by available structures of pre-60S particles, mature 60S subunits, and 80S ribosomes, are referred to as proximal or non-proximal RPs. The variation of the enrichment is represented by boxes highlighting the interquartile range of each data set, while the whiskers indicate the minimal and maximal limits of the distribution; outliers are shown as dots, the median by a line, and the average by a cross. **(D-G)** Location of Rpl3 and its neighbouring proteins on different pre-60S intermediates: **(D)** early nucleolar pre-60S (state 2) (PDB: 6C0F [97]), **(E)** late nucleolar pre-60S (state E1) (PDB: 7R7A [98]), **(F)** nucleoplasmic pre-60S (Rix1-Rea1) (PDB: 6YLH [99]), **(G)** cytoplasmic pre-60S (Lsg1-TAP, state I) (PDB: 6RZZ [100]). The indicated proteins have been highlighted in different colours; other r-proteins and AFs are coloured in light cyan and (pre-)rRNAs in light grey.

By employing this approach, we have already shown in a previous study that Rrb1 is strongly enriched and highly abundant in the streptavidin pull-down of cells expressing the C-terminally TurboID-tagged Rpl3 bait [94], indicating that our TurboID-based proximity labelling approach should in principle be well suited for the identification of new candidate DCs. Therefore, and initially focusing on the LSU r-proteins, we decided to directly perform TurboID assays with all 46 LSU r-proteins. Since 35 of the 46 different LSU r-proteins are encoded by two paralogous genes (referred to as the A and B copy), yielding identical or nearly identical r-proteins (denoted by an a or b after the r-protein name), we selected in almost all cases the paralogous r-protein whose absence was shown to be lethal or to confer the stronger growth defect [101], which largely correlates with their higher estimated abundance [102], as bait for the assay. Considering that five of the eight established DCs have been shown to recognize the N-terminal extremity of their r-protein client [41,44–46,103,104], we chose, reasoning that an N-terminally fused TurboID moiety might potentially affect the capacity of an r-protein bait to interact with its prospective DC, to first conduct the TurboID-based proximity labelling screen with the C-terminally TurboID-tagged LSU r-proteins. Nevertheless, to maximize the discovery of novel DCs, we also analysed in a second series the proxiOMEs of the N-terminally TurboID-tagged LSU r-proteins. As Rpl40 (eL40) is naturally expressed from the paralogous *UBI1*/*UBI2* genes as identical fusion proteins (Ubi1/2) containing an N-terminal ubiquitin moiety [36], which enhances the stable expression of Rpl40 and is rapidly, normally prior to the incorporation of Rpl40 into cytoplasmic pre-60S particles, removed from the Ubi1/2 precursors [38,39,100,105], we performed the proximity labelling with the Ubi1-TurboID and the TurboID-Rpl40a baits. By doing so, the natural context of ubiquitin release from the Ubi1 precursor is preserved for the Ubi1-TurboID bait, whereas the generation of Rpl40 lacking the N-terminal TurboID moiety is avoided by directly fusing TurboID to the N-terminal end of Rpl40. In each independent experimental set of the screen with the C-terminally TurboID-tagged LSU r-proteins, consisting for practical reasons of maximally nine different r-protein baits, we included Rpl3-TurboID as an internal quality control to validate the successful realization of the multi-step experiment. As most r-proteins exhibit a transient nuclear localization, we used two different C-terminally TurboID-tagged yEGFP fusion proteins, the mainly cytoplasmic yEGFP-TurboID and the nuclear NLS-yEGFP-TurboID (Supplementary Fig. S2A), as controls for determining the non-specific background labelling in the two cellular compartments containing unassembled and (pre-)ribosome-incorporated r-proteins. To calculate the specific enrichment of the purified proteins detected upon induced expression of a given TurboID-tagged r-protein, we divided the normalized abundance intensity (intensity-based absolute quantification (iBAQ) values) of each protein by the average of its normalized iBAQ abundance intensity, with prior imputation of missing values, in the two control purifications (see Materials and Methods). For graphical representation, we plotted the normalized iBAQ abundance (y-axis) of each protein detected in the streptavidin pull-down from cells expressing a given TurboID-tagged bait r-protein against its specific enrichment (x-axis); thus, displaying strongly enriched and highly abundant proteins in the upper right zone of these plots (see for example Fig. 1B). For the screen with the N-terminally TurboID-tagged r-proteins, we used in each independent experimental set TurboID-Rpl1b as the positive control, which enriches the novel DC Acl1 as well as several AFs that are in the proximity of Rpl1 on successive pre-60S particles (see below and Fig. 3A), and the mainly cytoplasmic TurboID-yEGFP and nuclear NLS-TurboID-yEGFP fusion proteins as background controls (Supplementary Fig. S2A). Having at disposal eight and six experimental replicates of the Rpl3-TurboID and the TurboID-Rpl1b experiment, respectively, enabled us to evaluate the reproducibility of our TurboID-based proximity labelling approach. Importantly, both the DC and the same set of enriched AFs and r-proteins could be detected at similar enrichment in all replicates of the Rpl3-TurboID or the TurboID-Rpl1b assays (Fig. 1C and Supplementary Fig. S10A). Given the high reproducibility and relatively low enrichment variability, we considered it fully sufficient to perform the TurboID assays of all LSU r-proteins in single replicates, even more so because we utilized this approach as a large-scale screen to identify candidate interactors and select promising ones for further validation. Moreover, the highly reproducible replicates encouraged us to confidently explore and evaluate, on the basis of the Rpl3-TurboID and TurboID-Rpl1b proxiOMEs, whether our TurboID-based proximity labelling approach was also suitable for the identification of r-protein neighbourhoods on pre-60S particles and mature 60S subunits (see below). To display the enormous amount of generated data in an easily accessible manner, the r-protein baits as well as the prominently enriched proteins, mainly corresponding to AFs and r-proteins, are labelled with their protein names in the graphical representations of the complete TurboID data set of all N– and C-terminally TurboID-tagged LSU r-proteins (Supplementary Fig. S1).

### TurboID-based proximity labelling reveals neighbourhoods of r-proteins on pre-ribosomal particles

While we have not experimentally assessed the capacity of the TurboID-tagged r-proteins to get assembled into pre-60S particles and mature 60S subunits, the identification of AFs or r-proteins that are known, based on the available cryo-EM and X-ray structures of pre-60S and mature 60S subunits, to be in the neighbourhood of a given TurboID-tagged r-protein can be used as a proxy for its successful incorporation. A closer inspection of the Rpl3-TurboID bait proxiOME revealed that, besides its DC Rrb1, several known early-associating (Dbp9, Loc1, Mak11, Nog1, Rlp24, Rrp15, Spb4, and Ssf1/2) and late-associating (Reh1 and Rei1) pre-60S AFs were reproducibly and quite specifically enriched (Fig. 1B and C and Supplementary Fig. S3), indicating that the Rpl3-TurboID fusion protein gets efficiently incorporated into early nucleolar pre-60S particles and remains associated with pre-60S particles along their entire maturation path. In line with this, Rpl3-TurboID, when expressed from a centromeric plasmid under the transcriptional control of its cognate promoter, complemented the absence of endogenous Rpl3 reasonably well on synthetic growth medium (Supplementary Fig. S2B). Importantly, and in support of the observed proximity labelling being due to a true physical proximity, most of these highly enriched AFs are indeed located in close proximity of the C-terminus of Rpl3, from which the TurboID moiety emerges at the LSU surface, on pre-60S particles. Five of the early-associating AFs (Mak11, Nog1, Rlp24, Rrp15, and Ssf1) could be visualized in the cryo-EM structure of the early nucleolar state 2 pre-60S particle [97], purified via the Nsa1 and Nop2 baits, and parts of Nog1, Rlp24, Rrp15, and Ssf1 are in immediate vicinity of Rpl3’s C-terminal end, while the β-propeller domain of Mak11 is located behind the Rpl3-proximal regions of Nog1 and Rlp24 (Fig. 1D). The extent of Nog1’s proximity becomes even more apparent on the late nucleolar state E1 pre-60S intermediate (Fig. 1E), where two additional α-helices within the C-terminal region of Nog1 can for the first time be seen in atomic models of pre-60S cryo-EM structures [98]. In the case of Spb4, which likely associates with intermediate nucleolar pre-60S particles (state D) and can be visualized both on state D and the late nucleolar state E pre-60S particles [98,106,107], its only partially resolved C-terminal extension projects from the LSU surface and thereby gets into proximity of the TurboID moiety attached to Rpl3’s C-terminal end (Fig. 1E; [98]). On the cytoplasmic state I pre-60S intermediate, purified via the Lsg1-TAP bait, the C-terminal end of Rei1 plunges deep into the polypeptide exit tunnel (PET), and the preceding Rei1 segments meander across the solvent-side surface of the LSU, leaving the unmodelled N-terminal part of Rei1 (residues 1-144) in close proximity of Rpl3’s C-terminal end (Fig. 1G; [100]). In subsequent snapshots of cytoplasmic, Lsg1-purified pre-60S subunits (states II-VI), Rei1 has been replaced by Reh1; however, the atomic model of Reh1 is limited to its C-terminal region that is inserted into the PET. Notably, the cryo-EM maps contain additional density that corresponds to the N-terminal parts of Rei1 and Reh1 in direct contact with the AF Tif6 [100]. In support of this, AlphaFold3 predictions revealed that the highly conserved C2H2-type zinc finger domain located at the N-terminal end of Rei1 (residues 1-45) and Reh1 (residues 1-44) has the capacity to interact with Tif6 [78]. Importantly, superposition of the AlphaFold models with the cryo-EM structures of Rei1– or Reh1-bound pre-60S subunits indicated that these predicted interactions between Tif6 and Rei1 or Reh1 could in principle also occur on pre-60S subunits (Supplementary Fig. S4D). In line with these in silico analyses, Y2H assays revealed a quite robust interaction of the N-terminal 66 residues of Rei1 (N66 construct; i.e., starting at the N-terminus and ending at residue 66) or the N-terminal 62 residues of Reh1 (N62 construct) with full-length Tif6 (Supplementary Fig. S4E). Tif6, however, which is in proximity of Rpl3’s C-terminal end almost throughout the entire pre-60S maturation path (Fig. 1D-G and Supplementary Fig. S4A and B), could only be detected via one non-biotinylated peptide in one of the eight experimental replicates of the TurboID assays with the Rpl3-TurboID bait. At first glance, this non-enrichment of Tif6 seems surprising, but it can easily be explained by the fact that Tif6 contains only two lysine residues (K18 and K93), with the K18 side chain engaging in a backbone hydrogen bond at the interior and only the K93 side chain being surface-exposed and in reasonable proximity of Rpl3’ C-terminal end; thus, limiting the identification of this long-lasting pre-60S neighbourhood by our biotinylation-dependent proximity labelling approach. Similarly, while the direct Rpl3 neighbour Rpl24 (eL24), which gets incorporated into early cytoplasmic pre-60S particles upon Drg1-mediated release of the closely related AF Rlp24 [108–110], was reproducibly enriched, the Rpl3-proximal Rpl23 (from early nucleolar (state B/state 2) pre-60S particles onwards) showed a similarly low enrichment as non-proximal r-proteins (Fig. 1B and C and Supplementary Fig. S4C). Based on the inspection of pre-60S, mature 60S, and 80S structures, this non-enrichment is likely due to Rpl23 being efficiently shielded from the bioAMP cloud that is generated around the TurboID moiety fused to Rpl3’s C-terminal end by Rpl3, Rlp24/Rpl24, Nog1, Tif6, and helix H95 as well as by engaging in two intersubunit bridges (Fig. 1D and E, Supplementary Fig. S4A-C, and [111]).

While the specific enrichment of the above-mentioned AFs can be readily explained by their immediate proximity to Rpl3’s C-terminal end in the available pre-60S cryo-EM structures, the TurboID assays with the Rpl3-TurboID bait also indicated potential proximities with the reproducibly and highly enriched AFs Dbp9 and Loc1 (Fig. 1B and C and Supplementary Fig. S3). The DEAD-box RNA helicase Dbp9 is a substoichiometric component of primordial and early nucleolar pre-60S particles but is not visible in the cryo-EM structures of early nucleolar, Nsa1-defined pre-60S particles (state 2 and states A, B, and C) [97,106,112,113]; however, as Dbp9 and Rpl3 are functionally connected by a synthetic lethal relationship and as crosslinking MS revealed a potentially direct interaction between Dbp9 and the Rpl3-proximal r-protein Rpl9 (uL6) [114,115], the observed enrichment of Dbp9 in the Rpl3-TurboID assay likely reflects a true physical proximity. In further support of this possibility, the proximity labelling with N-terminally TurboID-tagged Rlp24 also revealed a substantial enrichment of Dbp9 (Supplementary Figs. S1 and S3); notably, the C-terminal end of Rpl3 and the N-terminal end of the adjacent Rlp24 project in a similar direction from the pre-60S surface [106]. In the case of Loc1, the available cryo-EM structures of pre-60S particles do not offer a straightforward explanation for its suggested proximity to Rpl3’s C-terminal end. While Loc1 is already substantially enriched in early and/or intermediate nucleolar pre-60S particles sequentially purified via the Nsa1 and Ytm1 baits [113], and a short stretch of Loc1 could recently be visualized on early nucleolar, Ssf1-defined pre-60S particles [116], a reasonable portion of Loc1 only starts to be visible in late nucleolar state E pre-60S particles purified via the Spb4 bait [98,107]. On these state E pre-60S particles, the modelled Loc1 segments (residues 10-63 and 74-148) are located on the opposite side of Rpl3 (Supplementary Fig. S4A); however, it cannot be excluded that parts of Loc1, especially its lysine-rich C-terminal region (residues 149-204), which is not resolved in the state E pre-60S particles and projects away from the pre-60S surface, could be within the radius of the bioAMP cloud generated by the TurboID moiety emanating from Rpl3’s C-terminus on earlier pre-60S particles.

Intriguingly, Rpl2 (uL2) is, after the Rpl3 bait protein, the most enriched r-protein in the Rpl3-TurboID assays (Fig. 1B and C). Rpl2 can for the first time be visualized on the early nucleoplasmic state NE1 pre-60S intermediate (Supplementary Fig. S4B; [98]); at this maturation stage, on subsequent pre-60S particles, and on the mature LSU, Rpl2 appears, however, to be too far away from Rpl3’s C-terminus to explain its substantial enrichment. The visible incorporation of Rpl2, i.e., of its central, well-structured part (residues 25-200), and the neighbouring Rpl43 (eL43) is enabled by and/or coincides with the release of several AFs, including Loc1, Rrp17, Noc2, Noc3, and Spb4, which is part of the major compositional and structural rearrangements occurring during the transition from the late nucleolar state E to the early nucleoplasmic state NE1/NE2 pre-60S particles [98,99]. On the state E2 pre-60S intermediate, where the rRNA-binding sites of the central Rpl2 part within folding domains II and III are occluded by Noc2 and Noc3, respectively, the unresolved rRNA region encompassing helices H65 to H71, which contains the main rRNA-binding sites (H66 and H68) of Rpl2 and participates in intersubunit bridges on mature 80S ribosomes [111], emanates next to the N-terminal part of Rrp17’s long, surface-exposed α-helix from the surface (Supplementary Fig. S4A). We speculate that Rpl2 could already be associated, likely via its extensive interactions with rRNA helices H66 and H68, with the state E2 and/or during the transition to the state NE1 pre-60S intermediate. In this scenario, by being associated with this unresolved and, thus, presumably flexible rRNA region, Rpl2 might get into close enough proximity of Rpl3’s C-terminus to account for its observed enrichment in the TurboID assays.

In support of the suitability of the TurboID-based proximity labelling approach for the detection of r-protein neighbourhoods on pre-60S particles, the single data set TurboID assays revealed transient proximities of LSU r-proteins with directly adjacent AFs that can also be observed in pre-60S cryo-EM structures. Nsa1, for example, was most enriched in the proximity labelling assays of the C-terminally TurboID-tagged Rpl26 (uL24) and the N-terminally TurboID-tagged Rpl35 (uL29), respectively (Supplementary Fig. S5A, left panel); these two r-proteins are in close vicinity of Nsa1 within early nucleolar pre-60S particles (state 2 and states A to D) [97,106], in both cases positioning the fused TurboID moiety on the pre-60S surface in reasonable proximity of Nsa1 (Supplementary Fig. S5A, right panel). Rrp1, another member of the four-component Nsa1 module, was found to be most strongly enriched in the assays with the N-terminally TurboID-tagged Rpl4, Rpl7 (uL30), and Rpl18 (eL18) baits (Supplementary Fig. S5B, upper panel). These three r-proteins are in immediate proximity and even in direct contact with Rrp1 from the earliest structurally characterized pre-60S assembly intermediate (the so-called Noc1-Noc2 RNP), on which Rrp1 can, contrary to Nsa1, already be visualized, until the state D pre-60S intermediate, from which the Nsa1 module gets, presumably concertedly, released by the AAA-ATPase Rix7 [97,106,112,117]. While the N-terminal 14 residues of Rpl18 are not resolved in the cryo-EM structures of these early nucleolar pre-60S intermediates, the surface-exposed N-termini of Rpl4 and Rpl7 are directly adjacent to Rrp1 (Supplementary Fig. S5B, lower panel). Nmd3 and Lsg1 were prominently enriched in the TurboID assays performed with C-terminally TurboID-tagged Ubi1 (Rpl40a) and N-terminally TurboID-tagged Rpl10 (Supplementary Fig. S5C, left panels), two r-proteins that only get assembled during the cytoplasmic pre-60S maturation phase and can first be visualized on the state III and state IV pre-60S intermediate, respectively [100]; thus, recapitulating their observed proximities on late cytoplasmic pre-60S intermediates purified via the Lsg1-TAP bait (Supplementary Fig. S5C, right panel). Intriguingly, Yvh1, which could only be visualized on the state I and state II pre-60S intermediates and, thus, prior to the visible incorporation of Rpl40 and Rpl10 [100], was also substantially enriched in these Rpl40 and Rpl10 TurboID assays (Supplementary Fig. S5D, upper panel), suggesting that Yvh1 is nevertheless simultaneously present with Rpl40 and/or Rpl10 on cytoplasmic pre-60S intermediates. Accordingly, the order of cytoplasmic maturation events might be less hierarchical than deduced from the cryo-EM snapshots of consecutive Lsg1-purified pre-60S intermediates [100]. In agreement with this possibility, cryo-EM structures of late cytoplasmic pre-60S intermediates purified via the Nmd3-TAP bait from cells treated with the Drg1 inhibitor diazaborine revealed the simultaneous presence of Yvh1 and Rpl40 on pre-Lsg1 and Lsg1-engaged pre-60S particles and of Yvh1, Rpl40, and Rpl10 on the Rpl10-inserted pre-60S particle [118] (Supplementary Fig. S5D, lower panel). However, it cannot be excluded that partial inhibition of Drg1 or the fused TurboID moiety, especially in the case of Rpl40, could delay the release of Yvh1. Finally, and as briefly outlined later on, the TurboID assays with Rpl1, especially the ones with N-terminally TurboID-tagged Rpl1 performed in six experimental replicates, revealed transient proximities of Rpl1 to distinct AFs on consecutive pre-60S intermediates as well as its longer-lasting proximities to r-proteins on mature 60S and assembled 80S ribosomes (Fig. 3A and Supplementary Fig. S10).

We conclude that a rather specific and substantial enrichment of a given AF is a strong indication for its immediate proximity to the TurboID-tagged LSU r-protein bait on pre-60S particles. Moreover, it appears that our TurboID-based proximity labelling approach quite selectively highlights the neighbouring proteins (AFs and r-proteins) of LSU r-proteins that are most proximal to the fused TurboID enzyme on pre-60S intermediates and mature 60S subunits.

### TurboID-based proximity labelling is suitable to identify DCs of r-proteins

To fully assess the suitability of our TurboID-based proximity labelling approach for the identification of DCs, we also performed TurboID assays with N– and C-terminally TurboID-tagged Rps2, Rps3, and Rps26. Six of the eight known DCs, namely Rrb1, Syo1, Bcp1, Tsr4, Yar1, and Tsr2, could be well detected and were substantially enriched in one of the two or both TurboID assays of their respective r-protein client Rpl3, Rpl5, Rpl23b, Rps2, Rps3, and Rps26a (Fig. 1B and C and Supplementary Fig. S6). On the other hand, our TurboID-based proximity labelling screen would not have enabled the identification of Acl4 and Sqt1 as candidate DCs of Rpl4 and Rpl10, respectively. While Acl4 could not be detected in the assay with the C-terminally TurboID-tagged Rpl4a bait, it was present at relatively low abundance and weakly enriched in the streptavidin pull-down of cells expressing N-terminally TurboID-tagged Rpl4a. However, as Acl4 was, if at all, only detected at low intensity via few peptides across all LSU r-protein TurboID assays and not always present in the control purifications of the different experimental series, increasing the variability and, thus, potentially resulting in a randomly distributed low-level enrichment, its enrichment was not sufficiently prominent or specific to unequivocally infer a true positive proximity to Rpl4 (Supplementary Fig. S7). This weak enrichment was surprising as N-terminally TAP-tagged Rpl4a gets efficiently incorporated into mature 60S subunits and enables the co-purification of Acl4 [34]. Moreover, by recognizing and embracing Rpl4’s long internal loop (residues 44-113) via its concave surface [20,34], Acl4 is closely connected to Rpl4’s globular domain and, thus, expected to be in close enough proximity of the N-terminally fused TurboID moiety for the biotinylation of its surface-exposed lysine residues to occur. Rpl4’s globular domain is followed by an around 100-residue-long, eukaryote-specific C-terminal extension that spans across more than half the width (∼14 nm) of the solvent-side surface on mature 60S subunits [34,86,111], indicating that the C-terminally fused TurboID might be too far away from Acl4 to enable its biotinylation. In line with this possibility, performing the TurboID assay with C-terminally TurboID-tagged Rpl4 lacking its C-terminal extension (Rpl4a.N264-TurboID construct) resulted in a pronounced enrichment of Acl4 (Supplementary Fig. S6J). Sqt1, which accommodates residues 2-15 of Rpl10 on the negatively charged top surface of its WD-repeat β-propeller domain [41], was not detected in any of the LSU r-protein TurboIDs. According to this mode of interaction, the TurboID moiety fused to Rpl10’s C-terminal end, which is separated from the universally conserved uL16 core by an around 50-residue-long, eukaryote-specific C-terminal extension, might be too far away and/or too flexibly attached to enclose Sqt1 in a well-defined bioAMP cloud. The N-terminally fused TurboID, however, should in principle be appropriately positioned to enable biotinylation of at least some of the 18, mostly surface-exposed, lysine residues of Sqt1. While Y2H assays did not provide evidence for a negative impact of an N-terminally located Gal4 DNA-binding domain (G4BD) on the interaction between Rpl10 and Sqt1 (Supplementary Fig. S2C), we cannot rule out, as Sqt1 was shown to exhibit a twofold higher affinity for an N-terminal Rpl10 peptide lacking the initiator methionine in vitro [41], that the available Sqt1 would preferentially interact with endogenously expressed Rpl10 in vivo. Additionally, the equilibrium might be further shifted towards the recognition of untagged Rpl10 as the N-terminally fused TurboID could lower the efficiency of co-translational Rpl10 capturing by Sqt1 [41]. In the case of Rpl3, whose N-terminal 15 residues are sufficient to mediate a robust Y2H interaction and are predicted to interact with the top surface of Rrb1’s seven-bladed WD-repeat β-propeller [41] (Supplementary Fig. S2D), the N-terminal location of the G4BD abolished the Y2H interaction with Rrb1 (Supplementary Fig. S2C); thus, offering a plausible explanation for the non-enrichment of Rrb1 in the proximity labelling assay with N-terminally TurboID-tagged Rpl3. Similarly, and in line with the much lower and less specific enrichment of Syo1 in the TurboID assay with the TurboID-Rpl5 bait, the Y2H interaction between Rpl5 and Syo1, which accommodates residues 2-20 of Rpl5 within an extended groove on its inner α-solenoid surface [46], was markedly reduced when the Rpl5 bait contained the G4BD at its N-terminal end (Supplementary Fig. S2C). Finally, the TurboID assays with Rpl11, the second binding partner of Syo1 and the 5S rRNA [46,47], revealed some enrichment of Syo1 with the C-terminally TurboID-tagged Rpl11 bait (Supplementary Fig. S6E).

To reciprocally explore the suitability of the TurboID-based proximity labelling approach for the detection of the r-protein clients of DCs and, if possible, to gain further insight into the mode of operation of the different DCs, we also performed TurboID assays with all eight N– and C-terminally TurboID-tagged DCs. In these two experimental series, we calculated the enrichment of each protein detected in the TurboID assay of a given DC by dividing its normalized iBAQ intensity by the median value of its normalized iBAQ intensities, with prior imputation of missing values, in the two control TurboIDs and the TurboIDs of the remaining seven DCs. Except for the TurboID assay with the C-terminally TurboID-tagged Syo1, which only yielded a relatively low enrichment of Rpl5, all other TurboID assays resulted in a prominent and specific enrichment of the respective r-protein client (Supplementary Fig. S8). However, Rpl11 was only very weakly enriched, while nevertheless being the most abundant r-protein, in the TurboID of the Syo1-TurboID bait, but not at all enriched in the assay with N-terminally TurboID-tagged Syo1 (Supplementary Fig. S8C). The reason for this is unclear, especially in the case of the C-terminally fused TurboID moiety, which is, according to the structures of Syo1-containing complexes from the thermophilic filamentous ascomycete *Chaetomium thermophilum*, expected to be in close proximity of Rpl11 [47,48]. Considering that Syo1 is not essential and that its tandem affinity purification co-enriches less Rpl11 than Rpl5 [46], the Syo1-enabled import of Rpl5 and Rpl11 is dispensable for the assembly and pre-60S incorporation of the 5S RNP and may not obligatorily involve the co-import of Rpl11, raising the possibility that Syo1, which is intimately functionally connected to Rpl5 by synthetic lethality and reciprocal dosage rescue [46], primarily acts, at least in *S. cerevisiae*, as a transport adaptor and DC of Rpl5. In line with the importin Kap104 mediating the nuclear import of Syo1, by recognizing, most likely in a co-translational manner [31], its N-terminally located proline-tyrosine NLS (PY-NLS) [46], Kap104 was specifically, albeit relatively weakly, enriched in the TurboID of the Syo1-TurboID bait (Supplementary Fig. S8C). Overall, and as expected for selective binding partners of individual r-proteins, the TurboID-based proximity labelling assays with DC baits only identified a limited number of specifically and highly enriched proteins. Strikingly, however, almost all DC TurboIDs revealed a specific enrichment of at least one AF or non-client r-protein, indicating that DCs are transiently associated with pre-ribosomal particles to coordinate the transfer of their r-protein client to its binding site. For example, the Rrb1 TurboIDs enriched, albeit to different extents, several members of the Npa1 complex [119,120] (Supplementary Fig. S8A), which is part of the primordial pre-60S particle and functionally connected to Rpl3 by synthetic lethal relationships [113,114,119,121,122]. The AF Mak16, whose close proximity to Rpl4 can already be observed on the Noc1-Noc2 RNP [117], was highly abundant and strongly enriched in both Acl4 TurboIDs (Supplementary Fig. S8B). The two AFs Rpf2 and Rrs1, which are part of the hexameric 5S RNP precursor (5S rRNA, Rpl5, Rpl11, Rpf2, Rrs1, and Syo1) and required for the incorporation of the 5S RNP into pre-60S particles [48,123], were noticeably enriched in the Syo1 TurboIDs (Supplementary Fig. S8C). The Rpf2-Rrs1 heterodimer is already associated with early nucleolar pre-60S particles, but can only be visualized in the cryo-EM structure of the early nucleoplasmic, Nog2-purified pre-60S intermediate, where it anchors the 5S RNP in its rotated, immature conformation [106,112,113,123–126]. Thus, the observed enrichment of Rpf2 and Rrs1 in the Syo1 TurboIDs could either be evidence for the in vivo existence of the hexameric 5S RNP precursor in *S. cerevisiae*, which could so far, unlike in *C. thermophilum*, not be experimentally observed under normal growth conditions [46,48], or could alternatively indicate that Syo1 is present during the transfer of the 5S RNP to and/or the completion of 5S RNP assembly at its initial binding site, mainly consisting of the pre-bound Rpf2-Rrs1 complex, on nucleolar pre-60S particles. In line with the transient association of Sqt1 with late cytoplasmic pre-60S subunits for coordinating the stable incorporation of Rpl10 with the Lsg1-mediated release of Nmd3 [41,127,128], Nmd3 was prominently enriched in the proximity labelling assay with N-terminally TurboID-tagged Sqt1, while a moderate enrichment of Lsg1 could be observed with both Sqt1 baits (Supplementary Fig. S8D). Notably, Bcp1 exhibited a more multifaceted proxiOME than the other seven DCs (Supplementary Fig. S8E). On the one hand, the two AFs Rlp24 and Mak11, which are in direct contact with or, respectively, in close proximity of Rpl23 (Fig. 1D), were enriched, strongly suggesting that Bcp1 is transiently associated with early nucleolar pre-60S particles at the time of Rpl23 incorporation. Additionally, the Bcp1 proxiOME also included an already validated binding partner of Rpl23 (Rkm1) and two potential direct interactors of Bcp1 (Kap60/Srp1 and Mss4). The SET domain-containing methyltransferase Rkm1, which is responsible for dimethylation of lysine residues K106 and K110 of Rpl23 but dispensable for optimal growth of yeast cells [129–131], was shown to form a binary complex with Rpl23, but not with Bcp1, and a trimeric complex with Rpl23 and Bcp1 in vitro [131]. Moreover, co-immunoprecipitation experiments have indicated an Rpl23-dependent interaction between Bcp1 and Rkm1 [131]; thus, our proximity labelling data, showing an enrichment of Rkm1 in both the Rpl23 and Bcp1 TurboIDs (Supplementary Figs. S6F and S8E), provide further evidence for the in vivo occurrence of the trimeric Bcp1-Rpl23-Rkm1 complex. Bcp1 contains two predicted classical NLS regions [132], the first (residues 11-20) within its N-terminal extension and the second (residues 219-234) within an internal loop. According to AlphaFold3 predictions [78], positively charged side chains of the first and second NLS region interact in a canonical manner with the minor and major NLS-binding site [133], respectively, of the importin-α Kap60 (Supplementary Fig. S9A, left panels), suggesting that the transport adaptor Kap60, in conjunction with the importin-β Kap95, could mediate the nuclear import of Bcp1. Mss4 is an essential phosphatidylinositol-4-phosphate 5-kinase that almost exclusively localizes to the plasma membrane [134,135]. Interestingly, Mss4 also contains a functional NLS (residues 347-364), and overexpressed C-terminally GFP-tagged Mss4 exhibits, besides its normal location at the plasma membrane, a nuclear localization, which is no longer observed in cells lacking the importin-β Kap123 [135]. Moreover, the C-terminally GFP-tagged Mss4-1 mutant protein, bearing the D127N and L393P substitutions, was shown to almost exclusively accumulate in the nucleus at the non-permissive temperature. Notably, while absence of Kap123 abrogated the nuclear accumulation and conferred a cytoplasmic localization, overexpression of Bcp1 fully restored the plasma membrane localization of Mss4-1-GFP [135]. This data was, by also considering the finding that a temperature-sensitive *bcp1* allele (F241S) conferred a pre-60S export defect, interpreted as Bcp1 being responsible for nuclear export of Mss4 [135]. In line with their physical proximity suggested by the Bcp1 TurboID assays, AlphaFold3 predicted with good confidence the formation of a Bcp1-Mss4 complex (Supplementary Fig. S9A, right panels). According to this model, Bcp1 interacts with the first subdomain (residues 376-555) of the phosphatidylinositol phosphate kinase (PIPK) domain (residues 376-756) as well as with a segment (residues 335-357) that precedes the first PIPK subdomain and notably encompasses part of the NLS region of Mss4. Consistent with this predicted mode of interaction, an Mss4 variant (residues 332-565) consisting essentially of the N-terminally extended first PIPK subdomain showed a robust Y2H interaction with full-length Bcp1 (Supplementary Fig. S9B). Moreover, as an Mss4 variant (residues 371-565) comprising only the first PIPK subdomain exhibited a reduced Y2H interaction, it appears that both predicted Bcp1-binding surfaces of Mss4 are necessary for an optimal interaction. Interestingly, in the structure model of the Bcp1-Mss4 complex, Bcp1’s internal loop, whose first negatively charged residues are predicted to contact a positively charged surface on the first PIPK subdomain, and the NLS region of Mss4 are not in a configuration that would grant access to their respective importin, thereby providing a plausible explanation for how the Bcp1-Mss4 interaction would prevent the nuclear entry of Mss4. Since Bcp1 binding is predicted to efficiently shield all prominent positively charged surfaces of Mss4 (Supplementary Fig. S9C), it is likely that Bcp1 also ensures the safe transfer of Mss4 to the plasma membrane, a scenario that is fully compatible with the previously reported cell biological and genetic data [135].

We conclude that despite certain limitations, such as the distance between the fused TurboID moiety and the interacting DC, the requirement for a native context of N-terminally located DC-binding sites on r-proteins, and a potentially limited number of surface-accessible lysine residues on the predominantly negatively charged surface of DCs, our TurboID-based proximity labelling screen should be a valid approach to unveil hitherto unidentified DCs and other transient interaction partners, including importins (see below), of r-proteins. Moreover, as revealed by the identification of distinct AFs and/or r-proteins in the TurboID-based proximity labelling assays of DCs, a common feature of DCs appears to be their transient association with pre-ribosomal particles in order to facilitate the coordinated incorporation of their r-protein clients.

### The r-protein proxiOMEs also unveil potential interactions with importins

Interestingly, several LSU r-protein TurboIDs showed a noticeable enrichment of distinct importins (Fig. 2A-D, left panels; see also Supplementary Fig. S1). Here, we focused on obtaining evidence in support of a physical interaction between three of these importins, namely Kap114, Kap119 (also called Nmd5), and Kap121 (also called Pse1), and their potential cargo r-proteins Rpl6 (eL6) and Rpl40, Rpl30 (eL30), and Rpl38 (eL38), respectively. To this end, we first performed AlphaFold predictions, initially with AlphaFold-Multimer and then with AlphaFold3 [78,81], revealing that in each case almost the entire LSU r-protein is expected to be accommodated on the inner surface of the importin (Fig. 2A-D, middle panels). In the case of Rpl30 and Rpl40, their interaction with Kap119 and, respectively, Kap114 was predicted to also occur in the presence of GTP-bound Gsp1 (the *S. cerevisiae* counterpart of the mammalian GTPase Ran [136–139]) (Fig. 2D and data not shown), whose binding to importin-βs is generally sufficient to dissociate the bound import cargo in the nucleus [140–144], but may as well, as also observed in the case of Kap114 and Kap119 [145–151], only result in incomplete cargo release in vitro and/or be coupled to cargo transfer to specific chaperones or the cognate nuclear interaction site (reviewed in [152]). To provide first experimental evidence for a direct interaction between the above importins and r-proteins, we next performed Y2H assays. Since our well-functioning Y2H system, optimised for high specificity by expressing the bait and prey (as G4BD and Gal4 activation domain (G4AD) fusion proteins, respectively) from monocopy plasmids under the transcriptional control of the *ADH1* promoter, relies on the detection of interactions that occur at the promoters (*GAL1* and *GAL2*, respectively) of the *HIS3* and *ADE2* reporter genes in the nucleus, it will likely fail, due to the rapid binding of the highly abundant GTP-loaded Gsp1 [153], to reveal RanGTP-sensitive interactions between importin-βs and their r-protein import cargo. According to available co-structures, Gsp1 and Ran interact in their GTP-bound state with the four N-terminal HEAT repeats of all importins in a conserved manner, but they also engage in additional importin-specific contacts that centre mainly around HEAT repeats 7 and 8 or may even involve the four C-terminal HEAT repeats [150,152,154]. On the other hand, and not surprisingly, the observed cargo binding modes of the different importin-βs are quite diverse. However, importin-βs mostly employ the HEAT repeats, as well as interspersed loop regions, that succeed the N-terminal RanGTP-binding site to accommodate, often via negatively charged surfaces and hydrophobic pockets, linear sequences and/or folded domains within their inner concave surface [144,150,152,154–160]. We therefore reasoned that it should be possible, by progressively deleting the N-terminal RanGTP-binding site and, if necessary, adjacent HEAT repeats, to obtain importin-β variants with abolished or, at least, sufficiently reduced RanGTP-binding capacity that can be successfully employed in the classical nuclear Y2H system. In line with this, we have previously shown that the Kap121.302C variant (302C construct; i.e., starting at residue 302 and extending to the C-terminal end), which lacks HEAT repeats 1 to 7 and thus most RanGTP-interacting residues [144,154], engages in a productive Y2H interaction with its r-protein import cargo Rps2 [33]. To control that the full-length importin-β versions, when fused to the G4AD at their C-terminal end, have the capacity to form Y2H interactions, we assessed their interaction with the N-terminally G4BD-tagged Gsp1.Q71L mutant variant, which is the *S. cerevisiae* equivalent of the GTPase-deficient human Ran(Q69L) [161,162]. These Y2H assays revealed that Gsp1.Q71L interacts strongly with Kap121 but only relatively weakly with Kap114 and Kap119 (Fig. 2A-C, lower right panels), which may reflect the previously reported lower affinities of Kap114 and Kap119 for Gsp1.Q71L in vitro [153]. On the other hand, and as expected, the Kap121, Kap119, and Kap114 variants lacking part of or the complete N-terminal RanGTP-binding site no longer exhibited a Y2H interaction with Gsp1.Q71L (Fig. 2A-C, lower right panels). In line with the predicted Rpl38-binding region comprising almost exclusively residues of HEAT repeats 8 to 16, the Kap121.302C variant, fused at its C-terminus to the G4AD, showed a robust Y2H interaction with C-terminally G4BD-tagged Rpl38 (Fig. 2A, upper right panel). Conversely, no interaction between full-length Kap121 and Rpl38 could be observed, altogether suggesting that Rpl38 likely represents an import cargo of Kap121. Similarly, while not interacting with full-length Kap119, C-terminally G4BD-tagged Rpl30 showed a strong Y2H interaction with the Kap119.289C (starting after the predicted HEAT repeat 6) and Kap119.328C (beginning shortly before the predicted second α-helix of HEAT repeat 7) variants (Fig. 2B, upper right panel), which lack the predicted N-terminal binding site (HEAT repeats 1 to 3) of GTP-bound Gsp1 (data not shown), but apparently contain all necessary elements for the productive accommodation of Rpl30. For the Y2H assays with Kap114, we generated N-terminal deletion variants that lack partially (Kap114.75C; deletion of HEAT repeats 1 and 2) or completely (Kap114.159C; deletion of HEAT repeats 1 to 4) the N-terminal binding site (HEAT repeats 1 to 4) of GTP-bound Gsp1 [150]. Consistent with Rpl6 being predicted to extensively and almost exclusively contact Kap114 from HEAT repeat 5 onwards, N-terminally G4BD-tagged Rpl6 engaged in a robust Y2H interaction with the C-terminally G4AD-tagged Kap114.75C and Kap114.159C variants (Fig. 2C, upper right panel). On the other hand, Rpl6 did not show a Y2H interaction with full-length Kap114, indicating that Rpl6 could be an import cargo of Kap114. In support of this notion, a previous study detected Rpl6 via numerous peptides in MS analyses of Kap114 affinity purifications from the post-ribosomal supernatant of yeast cytosol [148]. Interestingly, the C-terminally G4BD-tagged Ubi1, i.e., Rpl40a-G4BD due to the rapid proteolytic removal of the ubiquitin moiety from the ubiquitin-Rpl40a precursor protein Ubi1 [36,39], only interacted with full-length Kap114 but not with the two N-terminal deletion variants (Fig. 2D, right panel), suggesting that the formation of a productive Y2H interaction between Rpl40 and Kap114 may depend on the binding of endogenous GTP-loaded Gsp1. Considering that Rpl40 gets only incorporated into pre-60S subunits in the cytoplasm and that one of the yeast importin-βs, Kap122 (also called Pdr6), has previously been shown to function as a biportin, i.e., a bidirectional transport receptor [100,105,118,154,163], it appears reasonable to speculate that Kap114, when associated with GTP-loaded Gsp1, could be responsible for the export of any Rpl40 that has inadvertently entered the nucleus in order to prevent its premature incorporation into pre-60S particles.

**Figure 2.**
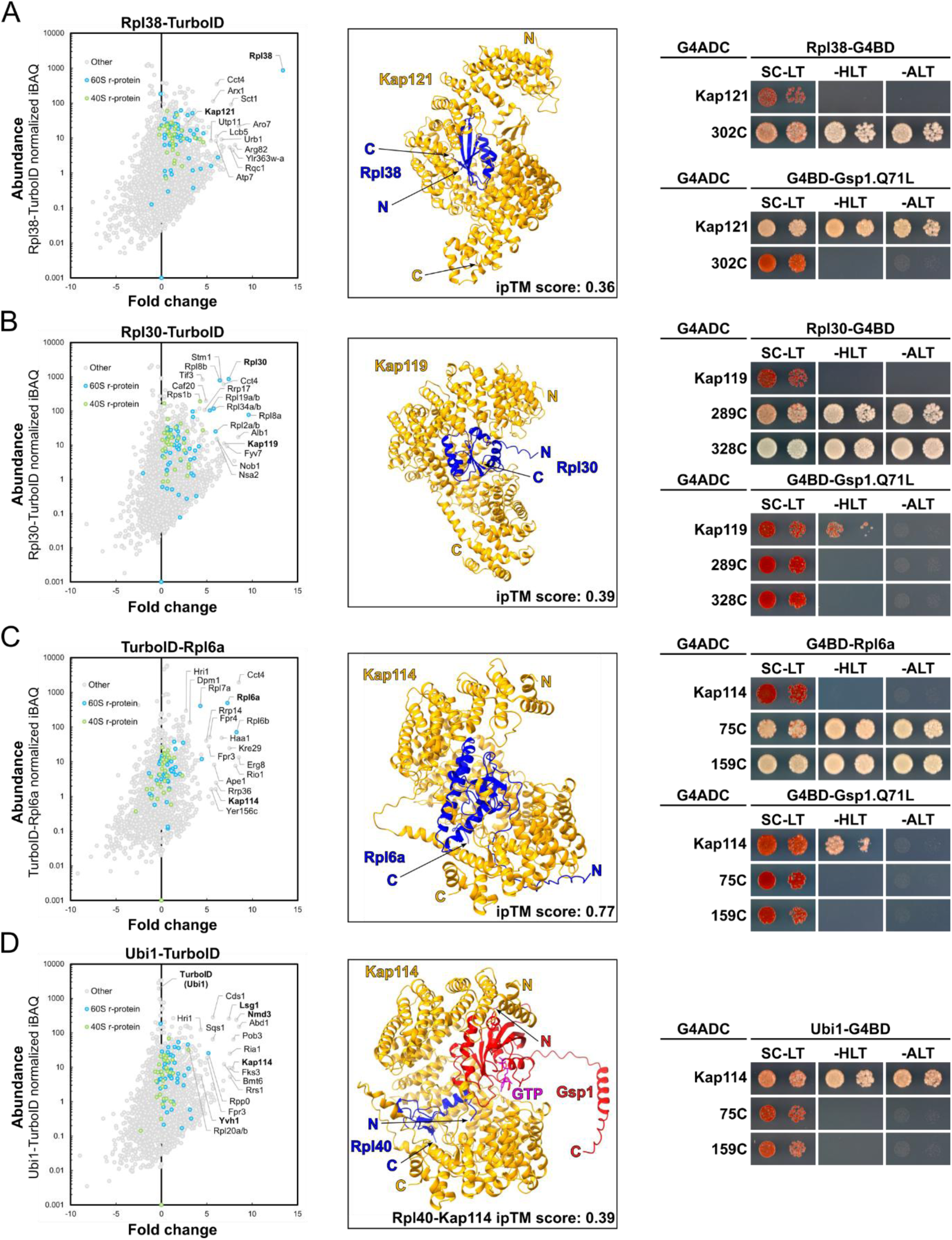
The r-protein proxiOMEs also unveil potential interactions with importins. (**A-D, left panels**) TurboID results obtained with the Rpl38-TurboID **(A)**, Rpl30-TurboID **(B)**, TurboID-Rpl6a **(C)**, and Ubi1-TurboID **(D)** r-protein baits. The bait r-proteins, the enriched importins, and, in the case of the Ubi1-TurboID, the proximal AFs Nmd3, Lsg1, and Yvh1 are written in bold. **(A-D, middle panels)** AlphaFold3 models of the Kap121-Rpl38 **(A)**, Kap119-Rpl30 **(B)**, Kap114-Rpl6a **(C)**, and Kap114-Rpl40-Gsp1-GTP **(D)** complexes. The respective ipTM scores are indicated. **(A-D, right panels)** Y2H interaction assays between Rpl38 **(A)**, Rpl30 **(B)**, Rpl6a **(C)**, or Ubi1 **(D)** and the indicated importins or their N-terminally truncated variants **(upper right panels)**, and between these importins or their N-terminally truncated variants and the Gsp1.Q71L mutant variant **(lower right panels)**. Plasmids expressing the indicated G4BD-tagged bait and G4AD-tagged prey proteins were co-transformed into the Y2H reporter strain PJ69-4A. Transformed cells were spotted in 10-fold serial dilution steps onto SC-Leu-Trp (SC-LT), SC-His-Leu-Trp (SC-HLT), and SC-Ade-Leu-Trp (SC-ALT) plates, which were incubated for 3 days at 30°C.

We conclude that our TurboID-based proximity labelling approach is also suitable to reveal transient interactions of r-proteins with distinct importins. However, as Kap123 and Kap121, suggested to be the main and partially redundant transport receptors for import of r-proteins [30], were not noticeably enriched, except for Kap123 with both Rpl8 (eL8) baits, in the TurboIDs of their supposed r-protein cargos [30,31,33,43,49] (Supplementary Fig. S1), we suspect that not all importins form sufficiently long-lived interactions with their r-protein cargos to warrant their identification by TurboID-based proximity labelling. Notably, and in contrast to the predominant recognition of short linear sequences by Kap121 and Kap123 [144,152,158,164,165], the four LSU r-proteins that enabled the TurboID-based identification of distinct importins are predicted to be almost entirely accommodated by the respective importin. This, together with their presumably not too short-lived association with the importins, may point to a necessity for being chaperoned by these importins and/or to a coupling of their release with the transfer to their pre-ribosomal binding site. Clearly, future experiments are required to address the in vivo relevance of the here unveiled interactions between the importins Kap114, Kap119, and Kap121 and their potential cargo r-proteins Rpl6 and Rpl40, Rpl30, and Rpl38.

### Identification of Acl1 and Bcl1 as potential interaction partners of Rpl1

The proximity labelling assays with N– and C-terminally TurboID-tagged Rpl1b enriched several r-proteins and AFs as well as the previously uncharacterized protein Ycr051w (Fig. 3A and Supplementary Fig. S10A), which we have named Acl1 for <u>A</u>nkyrin repeat <u>c</u>haperone of Rp<u>l1</u> (see below). In support of a highly specific biotinylation of proximal proteins by the two Rpl1b baits, both assays revealed a strong enrichment of the AF Nop2, whose globular domain is in direct contact with the concave surface of the L1 stalk on the late nucleolar state E pre-60S particles [98] (Supplementary Fig. S10B), and of the LSU r-protein Rpl13 (eL13), which is, especially its lysine-rich C-terminal region (residues 165-199), in proximity of the N– and C-terminal end of Rpl1 once the L1 stalk has moved from its pre-mature position in the state E pre-60S particles to its mature-like position on nucleoplasmic (from state NE2 onwards) pre-60S particles [99] (Supplementary Fig. S10C-H). The long-lasting proximity between Rpl1 and Rpl13, from nucleoplasmic pre-60S particles to mature 60S subunits, likely accounts for the strong enrichment and high abundance of Rpl13 in the streptavidin pull-downs of cells expressing TurboID-tagged Rpl1b. Moreover, the proximity labelling with N-terminally TurboID-tagged Rpl1b highlighted proximities with additional AFs that sequentially occur on successive pre-ribosomal particles. For example, parts of Loc1 (modelled residues 74-148 in state E1/E2 pre-60S structures) are located near the base of the L1 stalk on late nucleolar pre-60S particles [98] (Supplementary Fig. S10B), and, notably, the lysine-rich C-terminal region of Loc1 (residues 149-204), which is not visible in the cryo-EM structures of state E pre-60S particles, very likely emanates from the last modelled residue towards the upper part of the L1 stalk and is therefore expected to get into the bioAMP cloud around the Rpl1-fused TurboID. On nucleoplasmic pre-60S particles purified via the Nog2 bait, in which the 5S RNP is stably bound in a rotated, pre-mature orientation [126,166,167], the 5S RNP-anchoring Rpf2-Rrs1 heterodimer is facing the L1 stalk (Supplementary Fig. S10D), whose tip region including the bound Rpl1 is, however, not resolved, indicating that the L1 stalk still exhibits some flexibility at this assembly stage. On the subsequent Rix1-Rea1 pre-60S particle [99], the HEAT repeat domain of Sda1 contacts both the L1 stalk, with parts of Sda1 being directly adjacent to Rpl1, and the almost mature 5S RNP-containing central protuberance (CP), while its non-resolved C-terminal extension (residues 516-767) projects away from the pre-60S surface in an area that is surrounded by Rpl1 and Rpl5 (Supplementary Fig. S10E). From this maturation stage onwards, with the 5S RNP being in its mature conformation, Rpl5 is in proximity of Rpl1 (Supplementary Fig. S10E-H); however, the N– and C-terminal end of Rpl1 do not point in direction of Rpl5 on NPC-trapped pre-60S and mature 60S subunits [168,169], which agrees with the relatively low enrichment of Rpl5 in the Rpl1b TurboIDs. Rpl42 (eL42) can first be visualized on the late nuclear (LN) pre-60S intermediate [118], where it is located in proximity of the partially resolved L1 stalk. In structures with an atomic model of Rpl1 (e.g., NPC-trapped pre-60S), the N-terminal end of Rpl1 is in very close proximity of the surface-exposed β-strand domain of Rpl42 (Supplementary Fig. S10F-H). However, Rpl42 could only be identified in two of the six TurboID-Rpl1b replicates, likely because trypsin digestion of Rpl42 only yields very few peptides that are suitable for LC-MS/MS analysis. Finally, Rps25 (eS25) faces the rRNA side of the L1 stalk on the 80S ribosome [168] (Supplementary Fig. S10H), and its lysine-rich N-terminal extension, which is not resolved in the crystal structure of the 80S ribosome, could potentially come into close proximity of Rpl1.

Notably, both Rpl1 TurboIDs revealed a strong enrichment and a relatively high abundance of Acl1 (Fig. 3A), a protein of hitherto unknown function, suggesting that Acl1 could be a direct interaction partner, possibly a DC, of Rpl1. To obtain further indications that would reinforce this possibility, we first compared the features of Acl1 with the ones of the already established DCs. Acl1 is a relatively small protein consisting of 222 amino acids, and its N-terminal part (residues 1-126) is predicted to contain four ankyrin repeats [80,170,171] (AlphaFold model AF-P25631-F1, see Fig. 5B), which are expected to serve as a scaffold for mediating a protein-protein interaction (PPI). In this respect, Acl1 is highly similar to Yar1, the DC of Rps3 [172]. Yar1 also contains four N-terminal ankyrin repeats, and these were shown to mediate the interaction with the N-terminal α-helix of Rps3 [103,104]. Moreover, Acl1’s calculated isoelectric point (pI) of 4.25 is similar to the average pI (4.26; range 3.87-4.83) of all eight known DCs. Finally, comparative analysis of gene expression profiles, using the SPELL search engine on the *Saccharomyces* Genome Database (SGD) website [173,174], indicates that *ACL1* is transcriptionally co-regulated with genes belonging to the largely overlapping rRNA and ribosome biosynthesis (RRB) and ribosome biogenesis (RiBi) regulons [175,176]. In line with the nuclear pre-60S assembly stage of Rpl1 [98,99,177], Acl1-GFP, as indicated by a high-throughput localization study [178], accumulates in the nucleus but also displays some cytoplasmic localization; such a dual steady-state localization has been previously observed for the DCs Acl4, Bcp1, and Syo1 [34,46,86,135,178]. Contrary to most known DCs (see SGD website and [42]), Acl1 does not have a clearly identifiable human orthologue; moreover, our database searches only revealed clear orthologues in the Fungi kingdom, suggesting that Acl1 is likely a fungi-specific protein (see below and Supplementary Fig. S11). According to systematic analyses of gene deletion phenotypes (see SGD website and [179]), *ACL1* is a non-essential gene whose absence is not annotated to confer any severe growth defects. Furthermore, only very few potential physical and genetic interactions, mainly derived from high-throughput studies, have been reported for Acl1 (see SGD website). Notably, however, a large-scale screen for genetic interactions identified a weak negative genetic interaction (SGA score – 0.167) between the Δ*rpl1a* and Δ*acl1* null mutants [180]. Based on the strong enrichment of Acl1 in the Rpl1b TurboID assays and the above-mentioned supportive indications, we considered Acl1 as a promising candidate DC for the basic Rpl1 (pI 10.38) and, thus, selected it for further analysis. To validate the proximal relationship between Rpl1 and Acl1 and, at the same time, explore the proxiOME of Acl1, we first performed the reciprocal proximity labelling assays with both N– and C-terminally TurboID-tagged Acl1 (Fig. 3B). Not unexpectedly and like most known DCs (Supplementary Fig. S8), Acl1 exhibits a very limited proximal network. In strong support of their close proximity and suggestive of a direct physical interaction, Rpl1 was the most abundant of the few substantially enriched proteins in both Acl1 TurboID assays. Besides Rpl1, only the uncharacterized Ynl035c, which we have named Bcl1 for <u>B</u>eta-propeller <u>c</u>haperone of Rp<u>l1</u> (see below), was noticeably enriched in both assays, especially in the one of the N-terminally TurboID-tagged Acl1 bait. A closer inspection of the Rpl1b TurboID assays revealed that Bcl1 could also be detected among the enriched proteins, albeit only in four of the six experimental replicates and at relatively low abundance, in the proximity labelling with the N-terminally TurboID-tagged Rpl1b bait, raising the possibility that Bcl1 could be a direct interactor of Rpl1 or Acl1. Bcl1 is an acidic protein (pI 5.09) of 389 amino acids, which is predicted to fold into a typical seven-bladed WD-repeat β-propeller (residues 1-322) that is followed by a largely unstructured C-terminal extension ([181]; AlphaFold model AF-P53962-F1, see also Fig. 7B). In this respect, Bcl1 is similar to the known DCs Rrb1 and Sqt1, which mainly consist of a predicted seven-bladed or an experimentally-determined eight-bladed WD-repeat β-propeller that mediates the interaction with the N-terminal residues of Rpl3 or Rpl10 [41], and may likely engage in a PPI via its WD-repeat domain [182]. According to database searches (e.g., OrthoDB [183]), Bcl1 is conserved throughout eukaryotes and has a predicted human orthologue called WDR89. *BCL1* is a non-essential gene, and no pronounced phenotypes have been observed for Δ*bcl1* null mutant cells in large-scale surveys (see SGD website). Comparative analysis of gene expression profiles performed with the SPELL search engine on the SGD website reveals that the transcriptional co-regulation of *BCL1* with genes belonging to the RRB and RiBi regulons is less pronounced than in the case of *ACL1*, nevertheless several AF-encoding genes exhibit an expression profile that is similar to the one of *BCL1*. So far, only very few potential physical interactions have been reported for Bcl1 and these as well as its genetic interaction network, which is almost exclusively based on high-throughput data, do not provide any indications for a function in ribosome biogenesis that could be related to Rpl1. Finally, Bcl1-GFP, as indicated by a high-throughput study [178], exhibits nuclear localization.

**Figure 3.**
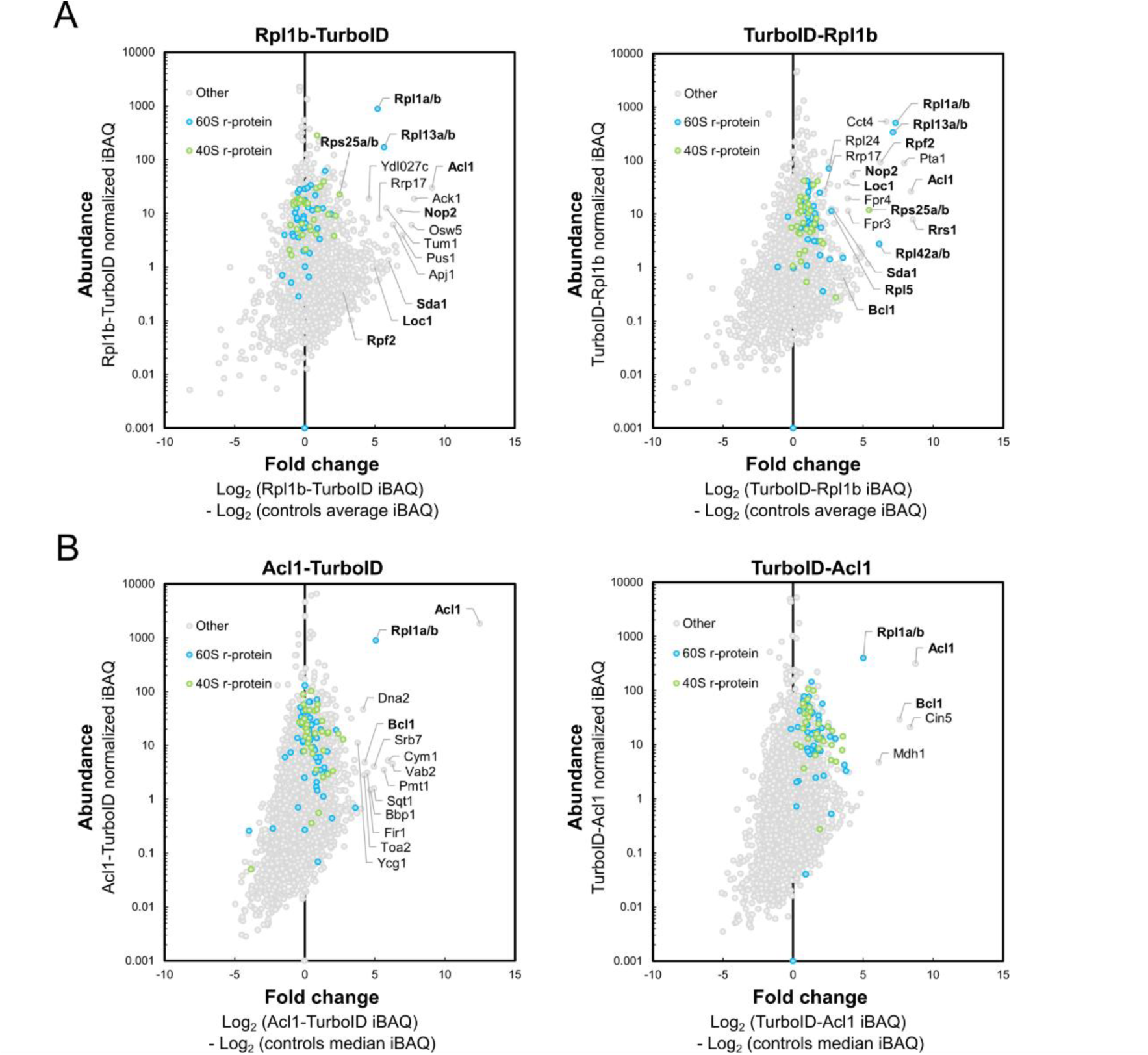
ProxiOMEs of Rpl1 and Acl1. **(A)** TurboID results obtained with the C– and N-terminally TurboID-tagged Rpl1b-TurboID (left panel) and TurboID-Rpl1b (right panel) bait proteins. **(B)** TurboID results obtained with Acl1-TurboID (left panel) and TurboID-Acl1 (right panel). The bait proteins and selected enriched proteins are written in bold.

**Figure 4.**
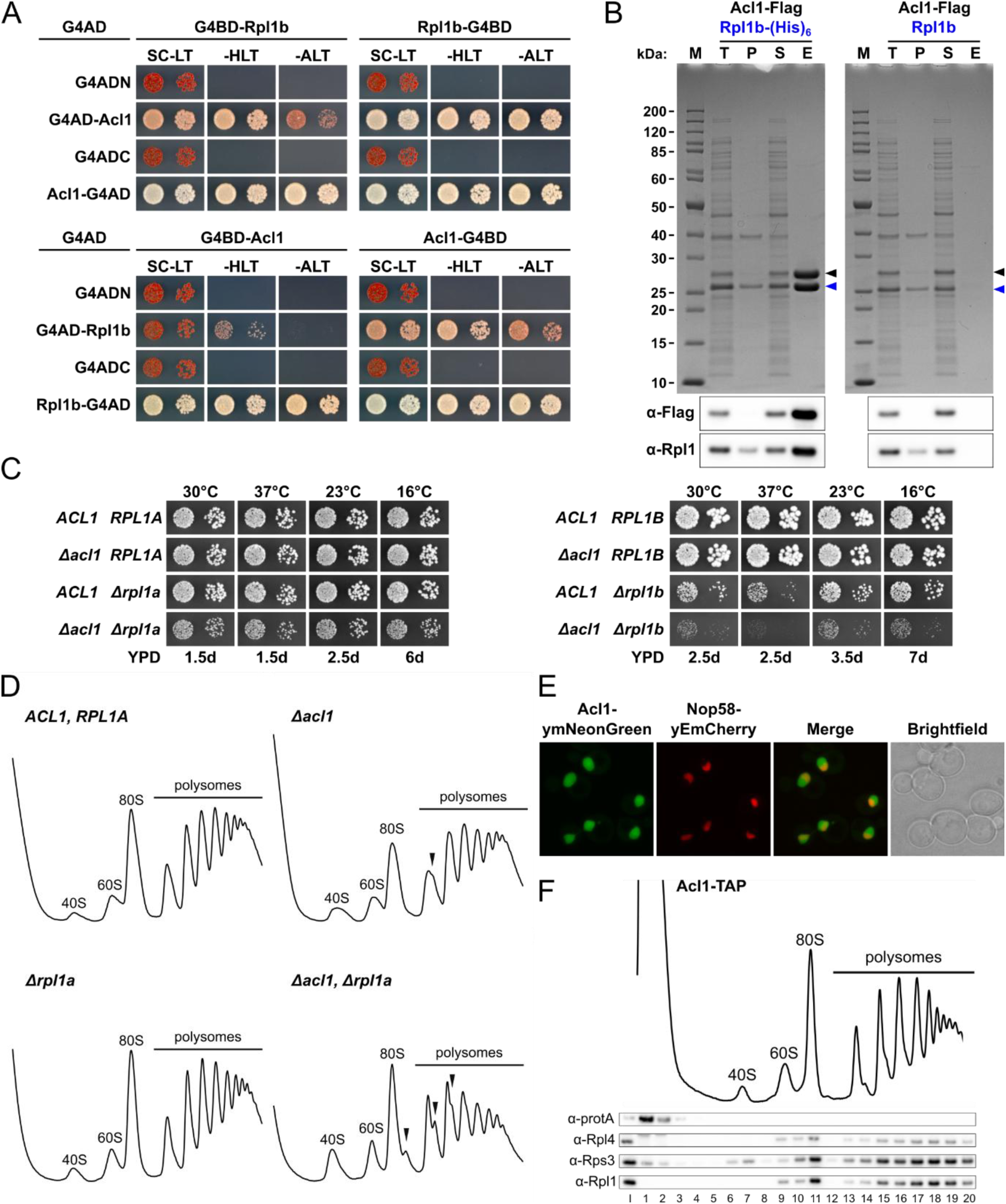
Acl1 is a direct interaction partner of Rpl1 and is involved in LSU biogenesis. **(A)** Y2H interaction assays between Rpl1 and Acl1. **(B)** In vitro binding assay between Rpl1 and Acl1. Untagged Rpl1b (Rpl1b) or C-terminally (His)_6_-tagged Rpl1b (Rpl1b-(His)_6_) and C-terminally Flag-tagged Acl1 (Acl1-Flag) were co-expressed in *E. coli* and purified by Ni-NTA affinity purification. Proteins were revealed by SDS-PAGE and Coomassie staining (top) or by Western blotting using anti-Flag and anti-Rpl1 antibodies (bottom). Bands corresponding to C-terminally (His)_6_-tagged and untagged Rpl1b or to Acl1-Flag are indicated by blue or black arrowheads. M: molecular weight standard, T: total extract, P: pellet fraction (insoluble proteins), S: soluble extract, E: imidazole eluate. **(C)** Negative genetic interactions between Δ*acl1* and Δ*rpl1a* (left panel) or Δ*rpl1b* (right panel). Strains with the indicated genotypes, derived from tetratype tetrads, were spotted in 10-fold serial dilution steps onto YPD plates, which were incubated for the indicated times at 30, 37, 23, and 16°C. **(D)** Polysome profiles of wild-type (*ACL1*, *RPL1A*), Δ*acl1*, Δ*rpl1a*, and Δ*acl1*/Δ*rpl1a* cells grown at 30°C in YPD medium. Whole cell lysates were prepared under polysome-preserving conditions in the presence of cycloheximide. Eight A_260_ units of clarified lysate were separated in 10-50% sucrose gradients by ultracentrifugation, and the A_254_ absorption profiles were recorded by continuous monitoring. Sedimentation is from left to right. Free 40S and 60S subunits, free 80S couples/monosomes (80S), and polysomes are labelled. Half-mer peaks are indicated by arrowheads. **(E)** The subcellular localization of genomically expressed Acl1-ymNeonGreen was assessed by fluorescence microscopy in cells grown at 30°C in SC-Ade medium. The subcellular position of the nucleolus was revealed by the nucleolar marker protein Nop58-yEmCherry, which was expressed from plasmid under the control of its cognate promoter. **(F)** An *ACL1*-TAP strain, expressing C-terminally TAP-tagged Acl1 from its genomic locus under the transcriptional control of its cognate promoter, was grown at 30°C in YPD medium, and the whole cell lysate was prepared under polysome-preserving conditions and analysed by sucrose gradient centrifugation and fractionation. Five A_260_ units of clarified lysate were separated in a 10-50% sucrose gradient by ultracentrifugation and the absorption profile was recorded by continuous monitoring at A_254_ (upper panel). 20 gradient fractions were collected from the top to the bottom at equidistant time intervals, and proteins were concentrated by TCA precipitation. An input (I) control (0.05 A_260_ units of the clarified lysate) and the gradient fractions (1-20) were analysed by Western blotting using anti-protA, anti-Rpl4, anti-Rps3, and anti-Rpl1 antibodies (lower panel).

**Figure 5.**
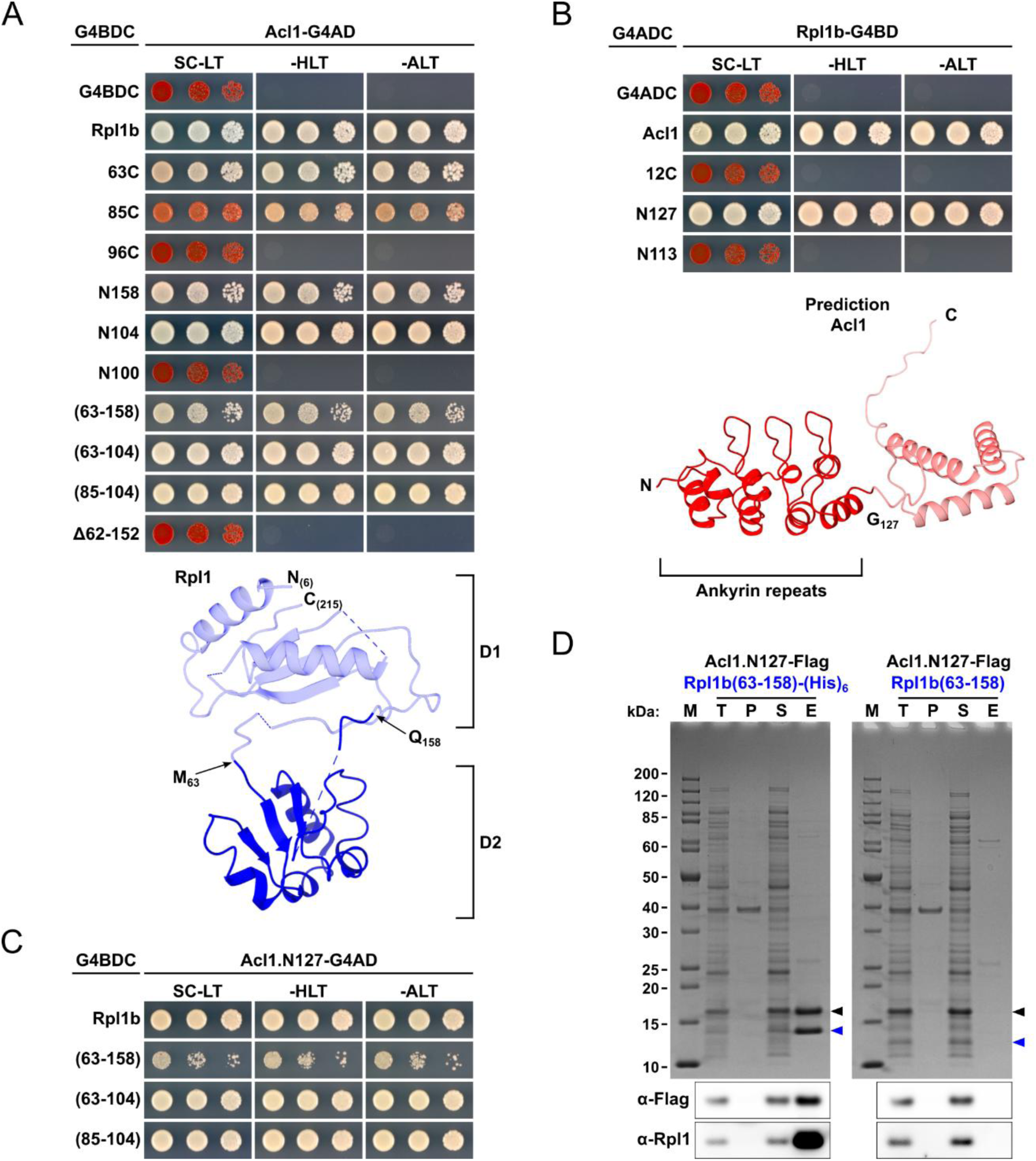
Acl1 interacts via its ankyrin repeat domain with the second domain of Rpl1. (**A, B**) Mapping of the respective interaction-mediating regions on Rpl1 **(A, upper panel)** and Acl1 **(B, upper panel)** by Y2H assays. **(A, B, lower panels)** Cartoon representation of the structure of Rpl1 (as observed in the cryo-EM structure of the Rix1-Rea1 pre-60S particle; PDB: 6YLH [99]) and the AlphaFold model of Acl1 (AF-P25631-F1-v4). The two structural domains of Rpl1 (D1 and D2) and the ankyrin repeat domain of Acl1 are indicated. **(C)** Y2H interaction between minimal internal Rpl1 fragments and the ankyrin repeat domain of Acl1 (Acl1.N127). **(D)** In vitro binding assay between Rpl1(63-158) and Acl1.N127. Untagged Rpl1b(63-158) or Rpl1b(63-158)-(His)_6_ and Acl1.N127-Flag were co-expressed in *E. coli* and purified by Ni-NTA affinity purification. Proteins were revealed by SDS-PAGE and Coomassie staining (top) or by Western blotting using anti-Flag and anti-Rpl1 antibodies (bottom). Bands corresponding to C-terminally (His)_6_-tagged and untagged Rpl1b(63-158) or to Acl1.N127-Flag are indicated by blue or black arrowheads. M: molecular weight standard, T: total extract, P: pellet fraction, S: soluble extract, E: imidazole eluate.

**Figure 6.**
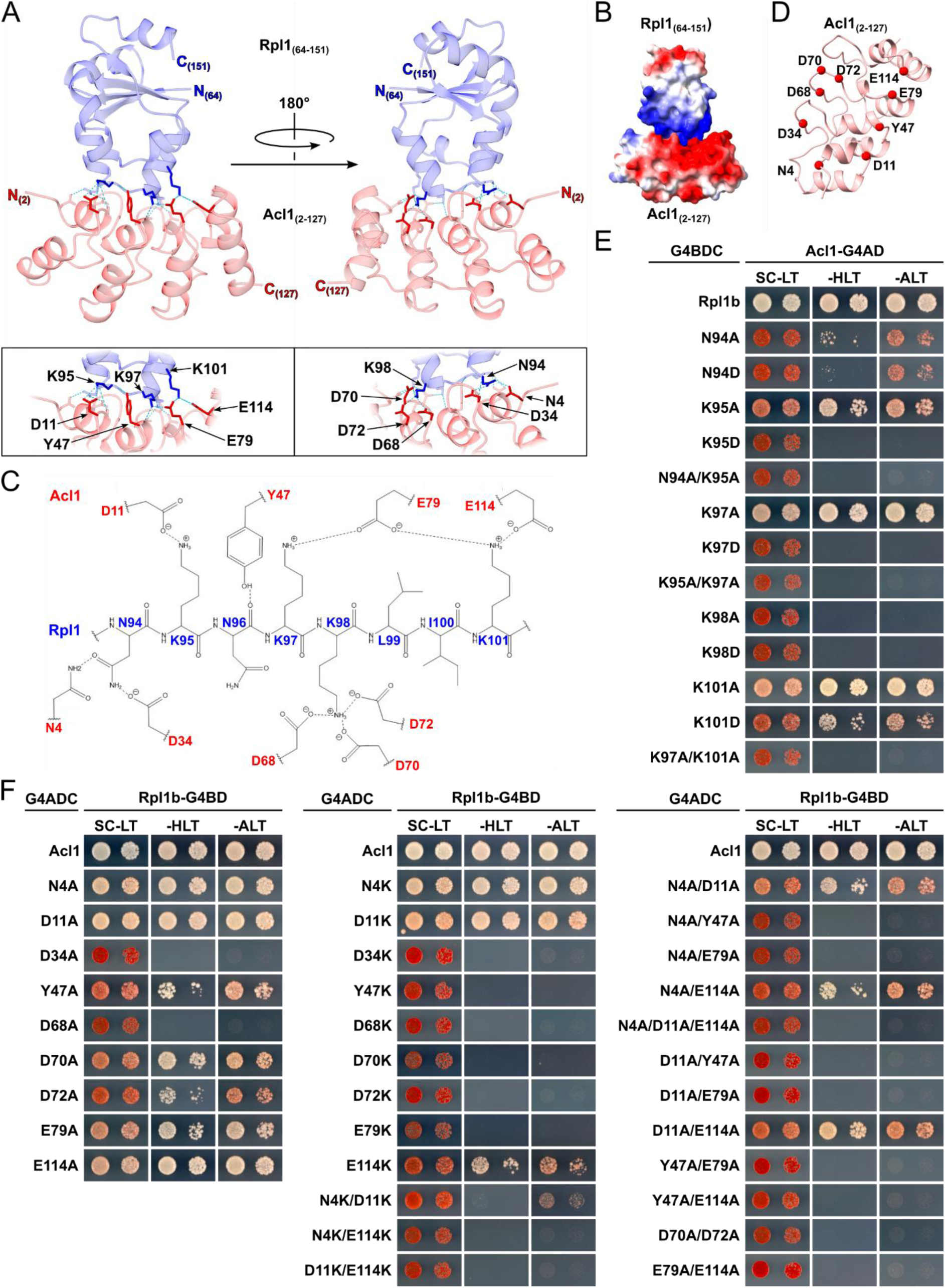
Crystal structure of Acl1’s ankyrin domain in complex with the second Rpl1 domain. (**A)** Cartoon representation of the X-ray co-structure of the second structural domain of Rpl1 (blue; resolved residues 64-151) and the ankyrin repeat domain Acl1 (red; resolved residues 2-127). In the lower part, the interacting residues are highlighted and labelled. The observed H-bonds are depicted as dotted lines. **(B)** Representation of the electrostatic surface potential of the Acl1-Rpl1 complex; red indicates negative potential and blue positive potential. **(C)** Schematic representation of the interaction mode. The backbone and side chains of Rpl1 residues 94-101 are shown as an elongated polypeptide and the side chains of the interacting Acl1 residues are indicated. H-bonds are depicted as dotted lines. **(D)** Distribution of the interaction-mediating Acl1 residues (highlighted by red dots) on the ankyrin repeat domain. **(E, F)** Mutational analysis of the Acl1-Rpl1 interaction interface. Y2H assays between the indicated Rpl1 mutant variants and Acl1 **(E)** and between the indicated Acl1 mutant variants and Rpl1 **(F)**. Single-letter abbreviations for the amino acid residues are as follows: A, Ala; D, Asp; E, Glu; K, Lys; N, Asn; Y, Tyr.

**Figure 7.**
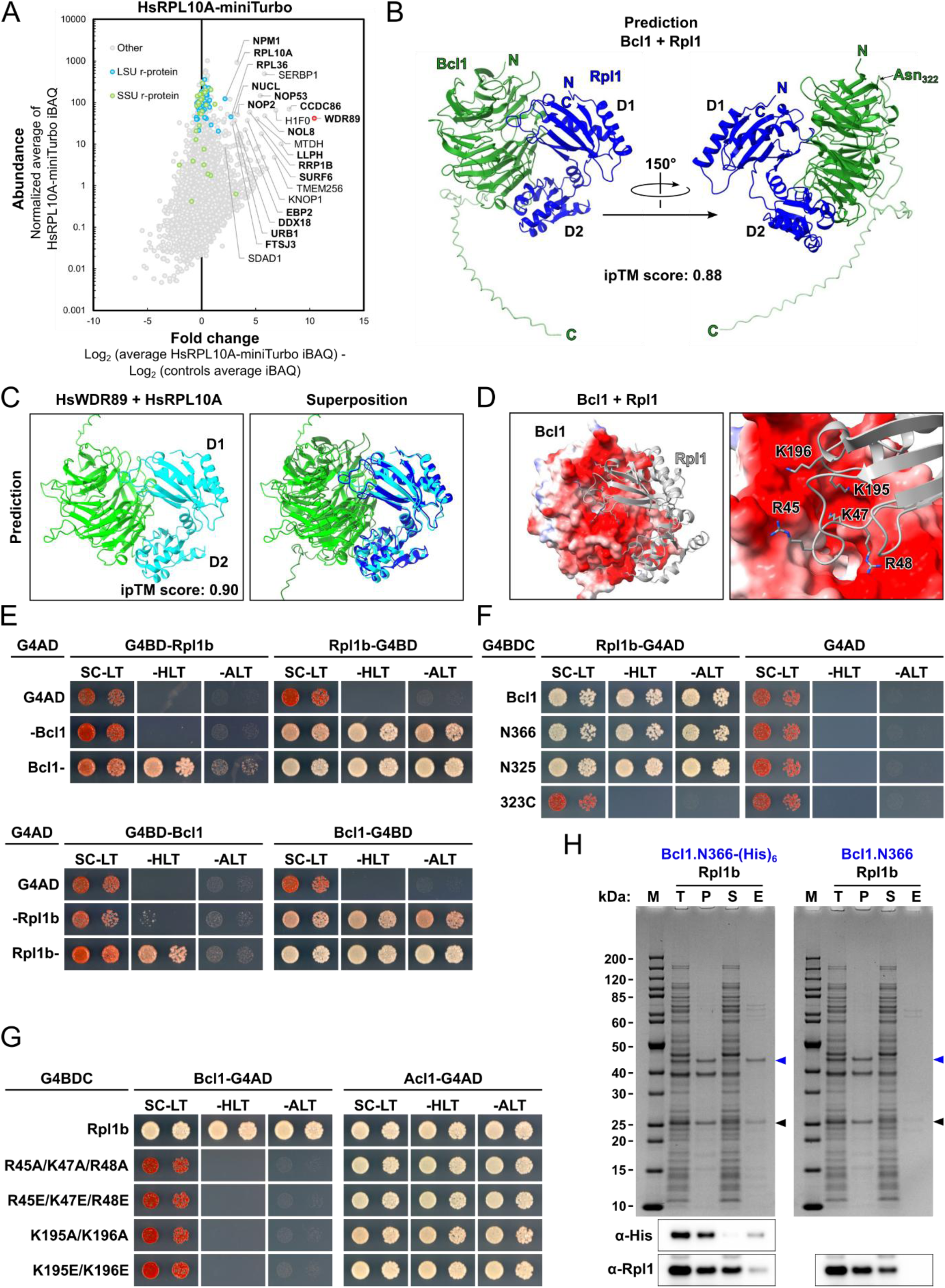
The conserved Bcl1 interacts via its WD40 β-propeller with Rpl1. **(A)** Proximity labelling assay with C-terminally miniTurbo-tagged RPL10A (HsRPL10A-miniTurbo), the orthologous human protein of yeast Rpl1, in HeLa cells. The assay was carried out in three experimental replicates, with miniTurbo-EGFP and EGFP-miniTurbo-NLS serving as cytoplasmic and nuclear background controls. The RPL10A bait r-protein and selected enriched proteins are written in bold, the red dot highlights the highly enriched WDR89. **(B)** AlphaFold3 model of the Bcl1-Rpl1 complex in two different orientations. The seven-bladed β-propeller domain of Bcl1 is coloured in green and the C-terminal extension in light green; the position of residue Asn322 (N322) is indicated to better visualize from where the C-terminal extension emanates. **(C)** AlphaFold3 model of the WDR89-RPL10A complex (left) and its structural superposition with the AlphaFold3 model of the Bcl1-Rpl1 complex (right). **(D)** Predicted electrostatic surface potential of Bcl1 (left) and close-up view of two of the three Rpl1 sites, indicating residues predicted to form H-bonds with Bcl1, that are in contact with the negatively charged top surface of the β-propeller (right). **(E-G)** Y2H interaction assays between the full-length Rpl1 and Bcl1 proteins **(E)**, between full-length Rpl1 and the C-terminally truncated Bcl1.N366 and Bcl1.N325 variants or the C-terminal extension of Bcl1 (323C) **(F)**, and between the indicated Rpl1 mutant variants and either Bcl1 or Acl1 **(G)**. Single-letter abbreviations for the amino acid residues are as follows: A, Ala; E, Glu; K, Lys; R, Arg. **(H)** In vitro binding assay between Bcl1.N366 and Rpl1. Bcl1.N366-(His)_6_ or Bcl1.N366 and Rpl1b were co-expressed in *E. coli* and purified by Ni-NTA affinity purification. Proteins were revealed by SDS-PAGE and Coomassie staining (top) or by Western blotting using anti-His and anti-Rpl1 antibodies (bottom). Bands corresponding to Bcl1.N366-(His)_6_ and Bcl1.N366 or to Rpl1b are indicated by blue or black arrowheads. M: molecular weight standard, T: total extract, P: pellet fraction, S: soluble extract, E: imidazole eluate.

### Acl1 is a direct interaction partner of Rpl1 and is involved in LSU biogenesis

As Bcl1 was only detected and, moreover, only moderately enriched in the proximity labelling assays with N-terminally TurboID-tagged Rpl1b, we first focused our efforts on obtaining further insight into the physical and functional interaction between Acl1 and Rpl1. To obtain evidence for a direct physical interaction, we first conducted Y2H assays between all possible combinations of N– and C-terminally G4BD– and G4AD-tagged Rpl1b and Acl1. For all but one combination (G4BD-Acl1 with G4AD-Rpl1b), we observed very strong or relatively strong Y2H interactions as indicated by the extent of growth, enabled by activation of the *HIS3* and *ADE2* reporter genes, on plates lacking histidine or adenine as well as by the white or only light red colony colour (Fig. 4A). To corroborate the Y2H data and to obtain definite evidence for their direct interaction, we co-expressed C-terminally (His)_6_-tagged Rpl1b and C-terminally Flag-tagged Acl1 in *E. coli* and subsequently purified the Rpl1b-(His)_6_ bait by Ni-NTA affinity purification. Analysis of the binding assay by SDS-PAGE and Coomassie staining as well as by Western blotting revealed an efficient and specific co-purification of Acl1-Flag with Rpl1b-(His)_6_ (Fig. 4B). When co-expressed with non-tagged Rpl1b, the recovery of Acl1-Flag in the imidazole eluate was, however, strongly reduced, showing that the observed substantial enrichment of Acl1-Flag with the Rpl1b-(His)_6_ bait was not due to its non-specific binding to the Ni-NTA agarose beads. We conclude that Acl1 is a direct interaction partner of Rpl1.

Next, we wished to gain insight into the functional relevance of Acl1 as an interaction partner of Rpl1. As haploid Δ*acl1* null mutant strains, generated *de novo* by deletion disruption of one *ACL1* copy in the diploid W303 background and subsequent tetrad dissection, did not display any clearly discernible growth defect on YPD plates at different incubation temperatures (Fig. 4C), we assessed whether the absence of Acl1 would have an effect on growth when Rpl1 is present in lower amounts. The essential and universally conserved r-protein Rpl1 is encoded by two paralogous genes, *RPL1A* and *RPL1B*, as proteins (Rpl1a and Rpl1b) with identical amino acid sequence [72]. In agreement with the higher *RPL1B* mRNA levels and the higher cellular abundance of Rpl1b [72,102,184], Δ*rpl1b* null mutant cells exhibited a strong slow-growth phenotype, which was especially pronounced at 37°C, while Δ*rpl1a* null mutant cells did not show any obvious growth defect (Fig. 4C), confirming the previously observed growth phenotypes of these null mutant strains in the same (W303) or a different genetic background [72,101,184]. Analysis of the growth characteristics of spore clones derived from tetratype tetrads, obtained by crossing Δ*rpl1a* with Δ*acl1* null mutant strains and subsequent sporulation of the heterozygous diploid strains, revealed that Δ*acl1*/Δ*rpl1a* double mutants exhibited a synthetic growth defect at all tested temperatures (Fig. 4C); thus, validating the negative genetic interaction previously indicated by a large-scale genetic interaction study [180]. Moreover, absence of Acl1 aggravated the growth defect of the Δ*rpl1b* null mutant to an extent that Δ*acl1*/Δ*rpl1b* double mutant cells were only barely growing (Fig. 4C). These genetic findings indicate that Acl1 becomes functionally relevant when Rpl1 is present in lower amounts, presumably owing to its contribution as a direct interaction partner to the provisioning of assembly-competent Rpl1. To evaluate the impact of the absence of Acl1 on ribosome biogenesis, both in the presence and absence of Rpl1a, we analysed the polysome profiles of wild-type, Δ*acl1*, Δ*rpl1a*, and Δ*acl1*/Δ*rpl1a* cells originating from the same tetratype tetrad (Fig. 4D). While the polysome profiles of wild-type and Δ*rpl1a* cells were almost indistinguishable, absence of Acl1 resulted in a slight increase in free 40S subunits and a faintly perceptible appearance of half-mers (Fig. 4D, see also Fig. 10B), which consist of mRNAs that are associated with one or multiple translating 80S ribosomes and a 43S pre-initiation complex awaiting subunit joining [185]. Notably, Δ*acl1*/Δ*rpl1a* double mutant cells showed a clearly noticeable accumulation of free 40S subunits and of half-mers that was accompanied by an overall decrease in polysome content, indicating, as the size of the 60S peak was not reduced, a defect in pre-60S export and/or subunit joining (Fig. 4D). In line with this, previous studies have shown that pre-60S subunits lacking Rpl1 are inefficiently exported from the nucleus and that the exported 60S subunits are less competent for subunit joining [177,184]. Additionally, we also examined the subcellular localization and (pre-)ribosome association of genomically expressed Acl1 that was fused at its C-terminal end to a yeast codon-optimized mNeonGreen (ymNeonGreen) or, respectively, the TAP tag. Fluorescence microscopy revealed that Acl1-ymNeonGreen localized almost exclusively to the nucleus, where it was predominantly located, as also judged by the co-localization with the nucleolar marker protein Nop58-yEmCherry, in the nucleoplasm (Fig. 4E). In line with the Acl1 TurboID assays not highlighting close proximities with known AFs or other r-proteins besides Rpl1 (Fig. 3B), sucrose gradient fractionation and subsequent Western blot analysis revealed that Acl1-TAP was exclusively present in the soluble fractions (Fig. 4F), altogether indicating that Acl1 is, if at all, only very transiently associated with pre-60S particles. We conclude that Acl1 contributes to the biogenesis of 60S subunits as a transient interaction partner of free Rpl1. Considering their robust interaction, it seems probable that Acl1 could already bind to Rpl1 in the cytoplasm and get transported together with Rpl1 to the nucleus, where it would dissociate from assembly-competent Rpl1 prior to or during the incorporation of Rpl1 into nucleolar pre-60S particles.

### Acl1 interacts via its ankyrin repeat domain with the second domain of Rpl1

To gain insight into their mode of interaction, we decided to first map the respective interaction-mediating regions by Y2H assays, expecting that this information should also help to obtain minimal Rpl1-Acl1 complexes that are amenable to structure determination by X-ray crystallography. As we started this endeavour before the creation of the AlphaFold Protein Structure Database or the availability of AlphaFold-Multimer to predict the structures of protein complexes [80,81,171], we based the design of our constructs on available Rpl1 structures (mainly the one of Rpl1 in the cryo-EM structure of the Rix1-Rea1 pre-60S particle and the one of human RPL10A in the cryo-EM structure of the cytoplasmic state C pre-60S particle [99,186]) and, in the case of Acl1, on multiple sequence alignments (T-Coffee [85]) and secondary structure predictions (PSIPRED [187]). Notably, Rpl1 mainly consists of two separate structural domains (Fig. 5A, lower panel). The first domain (D1) is formed both by N-(approximately from residue 6 to 37) and C-terminal (approximately from residue 165 to 217) primary sequence segments, whereas the second domain (D2) is exclusively formed by an internal primary sequence segment (approximately from residue 64 to 153). To map the Acl1-binding surface on Rpl1, we therefore performed Y2H assays with N-terminal, C-terminal, and internal deletion variants as well as with internal fragments of Rpl1, all C-terminally fused to the G4BD (for the complete Y2H mapping data set, see Supplementary Fig. S12A). These mapping experiments revealed that the second domain of Rpl1 is both sufficient and required for the interaction with C-terminally G4AD-tagged full-length Acl1 and that a short segment within D2 (residues 85-104) contains the relevant interaction-mediating residues (Fig. 5A, upper panel). The Y2H assays with N– and C-terminal deletion variants of Acl1 showed that the N-terminal 127 residues, essentially containing the four predicted ankyrin repeats (Fig. 5B, lower panel), correspond to the minimal fragment mediating full interaction with full-length Rpl1 (Fig. 5B, upper panel; for the complete Y2H mapping data set, see Supplementary Fig. S12B). Further Y2H assays revealed that the minimal Rpl1 fragments (residues 63-158, 63-104, and 85-104) exhibiting a strong interaction with full-length Acl1 still interacted equally well with the C-terminally truncated Acl1.N127 variant (Fig. 5C). In agreement with these Y2H results, the D2-containing Rpl1 fragment (residues 63-158) and C-terminally Flag-tagged Acl1.N127 formed, when co-expressed in *E. coli* and purified via the C-terminally (His)_6_-tagged Rpl1b bait by Ni-NTA affinity purification, a seemingly stoichiometric complex in vitro, as indicated by the similar intensity of the two Coomassie-stained bands (Fig. 5D).

Next, we aimed to exploit the efficient in vitro production of this minimal complex to illuminate the molecular details of the Rpl1-Acl1 interaction by X-ray crystallography. To this end, we co-expressed Rpl1b(63-158)-(His)_6_ and untagged Acl1.N127 in *E. coli* and purified the complex by Ni-affinity chromatography followed by size exclusion chromatography and subjected the purified complex to crystallisation screens. Gratifyingly, diffracting crystals were obtained and the co-structure of Rpl1 (residues 64-151) and Acl1 (residues 2-127) could be solved at 2.6 Å by molecular replacement (Fig. 6A; see Materials and Methods and Supplementary Table S1). Analysis of the electrostatic surface properties revealed that the negatively charged ankyrin repeat surface accommodates the complementary, positively charged surface of Rpl1 (Fig. 6B and Supplementary Fig. S13A). Accordingly, the interaction interface is primarily established by salt bridges and hydrogen bonding networks (Fig. 6C). In good agreement with the Y2H mapping data (Fig. 5A), the five Rpl1 residues involved in the interaction, i.e., Asn94, Lys95, Lys97, Lys98, and Lys101, cluster within a short segment (residues 94-101) of the D2 domain (Fig. 6A and C); moreover, these residues are conserved in Rpl1 orthologues from metazoans, plants, and other fungi (Supplementary Fig. S13D). Notably, an Rpl1 segment (residues 98-105) containing the interaction-mediating residues Lys98 and Lys101 is in direct contact with the tip of the L1 stalk (rRNA residues 2481-2486 that are part of the bulge connecting 25S rRNA helices H78 and H77) both on state E pre-60S particles and mature 60S subunits [98,188] (Supplementary Fig. S13B and C), indicating that Acl1 and the L1 stalk interact in a mutually exclusive manner with the D2 domain of Rpl1. On Acl1, the residues involved in the interaction are located on the one hand at the end of the first α-helix (Asp11 and Tyr47) or within the short loop connecting the first and second α-helix (Glu79 and Glu114) of each ankyrin repeat and, on the other hand, within the β-turn at the end of the long repeat-connecting loops (Asp34, Asp68, Asp70, and Asp72), in both cases aligned in-line to jointly form the Rpl1-binding interface along the concave inner, groove-like surface of the ankyrin repeat domain (Fig. 6A and D). Moreover, residue Asn4, which is situated before the first α-helix, contributes by hydrogen bonding to the interaction with Rpl1 residue Asn94 (Fig. 6C). According to BLAST searches and multiple sequence alignments, most of the nine interaction-mediating residues are identical or highly similar in predicted Acl1-like proteins of the Fungi kingdom, especially in those belonging to the Dikarya subkingdom (Supplementary Fig. S11), suggesting that all fungal Acl1 homologues likely fulfil a conserved function as Rpl1-binding proteins. To get an idea of the importance of the above residues for the mutual interaction, we first individually substituted the five Rpl1 residues by alanine or aspartate and the nine Acl1 residues by alanine or lysine and assessed the effects of these amino acid exchanges on the interaction by Y2H assays. Of the individual alanine substitutions in Rpl1, only the N94A and the K98A exchange had a major impact and, respectively, either strongly reduced or completely abolished the Y2H interaction with Acl1 (Fig. 6E). Moreover, the K95A exchange resulted in a visibly weaker Y2H interaction, while the K97A and the K101A exchange only slightly, if at all, affected the interaction capacity of Rpl1. As expected, the charge reversal substitutions had a more drastic effect, leading to a complete loss (K95D and K97D) or a substantial weakening (K101D) of the Y2H interaction. However, replacing the carboxamide of Asn94 by a carboxyl group (N94D exchange) did not much more weaken the Y2H interaction than the N94A exchange. Finally, simultaneous alanine substitution of two neighbouring Rpl1 residues (i.e., N94A/K95A, K95A/K97A, and K97A/K101A), whose individual alanine substitution still permitted a reduced or almost full interaction, respectively, abolished the Y2H interaction with Acl1, indicating that these Rpl1 residues synergistically contribute to the interaction with Acl1. In line with the importance of Rpl1 residues Asn94 and Lys98 for the interaction, alanine substitution of the contacting Acl1 residues Asp34 and Asp68, respectively, eliminated the Y2H interaction (Fig. 6F, left panel). Moreover, while the N4A, D11A, and E114A exchanges had almost no effect, the Y47A, D70A, D72A, and E79A exchanges affected, albeit to slightly different extents, the interaction capacity of Acl1. Consistent with this, introducing the positively charged lysine at these positions (Y47K, D70K, D72K, and E79K exchanges) abolished the Y2H interaction (Fig. 6F, middle panel). However, this radical charge insertion or reversal did not observably (N4K and D11K) or completely (E114K) reduce the interaction capacity of these Acl1 variants, indicating that Glu114 and, especially, Asn4 and Asp11 are less important interaction determinants. Their contribution to Rpl1 binding became, however, apparent when assessing the effects of simultaneous amino acid substitutions. While the N4A/D11A, N4A/E114A, and D11A/E114A exchanges already resulted in a visibly weaker Y2H interaction, Acl1 variants bearing the analogous lysine substitutions or their triple alanine substitution only barely (N4K/D11K) or no longer (N4K/E114K, D11K/E114K, and N4A/D11A/E114A) interacted with Rpl1 (Fig. 6F, middle and right panels). Moreover, combining the N4A, D11A, and E114A substitutions with the Y47A or the E79A substitution, which on their own still permitted Acl1 to engage in a productive but reduced Y2H interaction with Rpl1, completely abolished the interaction capacity of Acl1 (Fig. 6F, right panel). Likewise, introducing the double Y47A/E79A exchange or reducing the negative charge within the second β-turn by the simultaneous D70A/D72A exchange also eliminated the Y2H interaction between Acl1 and Rpl1. To sum up, all Rpl1 and Acl1 residues that are prominently involved in the mutual interaction according to the crystal structure also contribute, albeit to different degrees, to the interaction in the Y2H assays, thus providing experimental support for the validity of the structurally visualized mode of interaction. Remarkably, our experimentally derived co-structure is practically congruent with the predicted structure model, indicating that AlphaFold3 can apparently predict co-structures of ankyrin repeat domains and their protein binding partners with good accuracy (Supplementary Fig. S13E).

### The conserved Bcl1 interacts via its WD40 β-propeller with Rpl1

As our homology and database searches, combined with sequence analyses and AlphaFold-Multimer predictions to evaluate the conservation of the above-determined interaction-mediating residues and the mode of mutual interaction, failed to identify clear eukaryotic Acl1 orthologues beyond the Fungi kingdom, Acl1 is likely a fungi-specific Rpl1-binding protein. Intrigued by this possibility as the majority of the known DCs are conserved throughout eukaryotes [42–44,189], we performed proximity labelling in HeLa cells with the human Rpl1 orthologue RPL10A (see Materials and Methods), fused at its C-terminus to the promiscuous miniTurbo biotin ligase variant [87], to explore the proxiOME of RPL10A in an unbiased manner and, if possible, identify its elusive, evolutionarily conserved binding partner. Enriched and abundantly detected proteins included nucleolin (NUCL), a resident of the dense fibrillar component of the nucleolus, nucleophosmin (NPM1), the scaffold protein of the granular component of the nucleolus, the early-acting AFs NOL8 (yeast Nop8) and URB1 (Npa1/Urb1), and several components of nucleolar and nucleoplasmic pre-60S particles, such as SURF6 (Rrp14), RRP1B (Rrp1), DDX18 (Has1), EBP2 (Ebp2), FTSJ3 (Spb1), LLPH (Ybl028c), CCDC86 (Cgr1), and NOP53 (Nop53) [190–196] (Fig. 7A). Notably, and as observed in the TurboID assay with C-terminally TurboID-tagged Rpl1b (Fig. 3A), NOP2, which is like yeast Nop2 in direct contact with the L1 stalk and can be visualized in close proximity of RPL10A on early to late nucleolar human pre-60S particles [98,197] (for the Rpl1-proximal location of Nop2 on the late nucleolar state E2 yeast pre-60S particle, see Supplementary Fig. S10B), was also enriched. Contrary to the proximity labelling with the Rpl1b-TurboID bait where Rpl13, but not the similarly proximal Rpl36 (eL36) (see PDB 8AGX; [188]), was found to be strongly enriched (Fig. 3A), only RPL36, which is like RPL13 in immediate proximity of the C-terminal end of RPL10A in the human 80S ribosome (see PDB 6QZP; [198]), but not RPL13 was enriched in the RPL10A miniTurbo assay (Fig. 7A). Moreover, a strong enrichment of the ribosome preservation factor SERBP1 (yeast Stm1), which is required for the activation of dormant 80S ribosomes [199], could be observed. Interestingly, while a human Acl1-like protein could not be revealed by this approach, the most enriched protein in the miniTurbo assay with the RPL10A bait was WDR89 (Fig. 7A), the predicted human orthologue of Bcl1. In line with our proximity labelling data, WDR89 had previously been identified in affinity purifications of C-terminally HA-tagged RPL10A from two different human cell lines in large-scale interactome studies (see BioPlex Explorer website; [200]), and AlphaFold3 predicted with high confidence the formation of a WDR89-RPL10A complex (Fig. 7C). Notably, Bcl1 and Rpl1 are predicted to form a binary complex that shares remarkable similarity with the WDR89-RPL10A complex (Fig. 7B and C), strongly suggesting that Bcl1 is the sought-after conserved binding partner of Rpl1. According to the structure models, the negatively charged top and side surface of the seven-bladed WD-repeat β-propeller (formed by residues 4-322 of Bcl1) accommodate conserved positively charged residues of the D1 domain, the D1-D2 linker segments, and, respectively, the D2 domain (Fig. 7B and D). In line with a direct interaction, Y2H assays, performed with all possible combinations of N– and C-terminally G4BD– and G4AD-tagged Rpl1b and Bcl1, revealed a robust interaction for half of the combinations, especially when both proteins were tagged at their C-terminal end (Fig. 7E). Moreover, and as predicted, the WD-repeat β-propeller domain (see Bcl1.N325 deletion variant) was sufficient to mediate an equally strong Y2H interaction as full-length Bcl1, whereas the largely unstructured C-terminal extension (residues 323-389) did not have the capacity to interact with Rpl1 (Fig. 7F). Focusing on predicted interactions that occur at the top surface of the WD-repeat β-propeller, we simultaneously mutated neighbouring positively charged Rpl1 residues of the D1 domain (K195 and K196) or the D1-D2 linker segment (R45, K47, and R48), representing two of the three main Rpl1 sites that are in contact with the top surface, either to alanine or glutamate and assessed their effects on the Y2H interaction. Notably, the introduction of alanines was in both cases already sufficient to eliminate the interaction with Bcl1, while the interaction with Acl1 remained unaffected even by the charge reversal exchanges (Fig. 7G), highlighting that these positively charged Rpl1 residues are relevant constituents of the Rpl1-Bcl1 interaction interface. In further support for their direct interaction, Rpl1 could be co-purified by Ni-NTA affinity purification with the C-terminally (His)_6_-tagged Bcl1.N366 variant, which lacks the highly basic last third of the C-terminal extension (theoretical pI of 11.26 of the last 23 residues) and exhibits greater solubility than full-length Bcl1 (data not shown), upon co-expression of the two proteins in *E. coli* (Fig. 7H). We conclude that Bcl1 is an evolutionarily conserved binding partner of Rpl1 that, in contrast to the relatively simple binding mode underlying the recognition of Rpl1 by the fungi-specific Acl1, appears to contact multiple positively charged sites within different structural regions (D1, D2, and interdomain linker segments) of Rpl1 in a rather sophisticated manner via two negatively charged surfaces of its WD-repeat β-propeller domain.

### Acl1 and Bcl1 functionally cooperate and can form a trimeric complex with Rpl1

Next, we examined the functional relevance of the direct interaction between Bcl1 and Rpl1. As haploid Δ*bcl1* null mutant strains, generated *de novo* by deletion disruption of one *BCL1* copy in diploid W303 cells, did not exhibit a discernible growth phenotype (Fig. 8A), we assessed whether absence of Bcl1 would, like the one of Acl1 (Fig. 4C), have an impact on growth when Rpl1 is present in reduced amounts. While the simultaneous absence of Bcl1 and Rpl1a (Δ*bcl1*/Δ*rpl1a* double mutant) did not affect growth (Fig. 8A), a weak, but clearly discernible enhancement of the slow-growth phenotype of Δ*rpl1b* mutant cells could be observed in the absence of Bcl1 (Δ*bcl1*/Δ*rpl1b* double mutant) at 30°C and, especially, at 37°C (Fig. 8B). Accordingly, the polysome profile of Δ*bcl1* cells was almost indistinguishable from the one of wild-type cells, and only a faintly perceptible formation of half-mers could be observed in Δ*bcl1*/Δ*rpl1a* double mutant cells (Supplementary Fig. S14A). Polysome profile analysis with Δ*bcl1*/Δ*rpl1b* double mutant cells revealed an accumulation of free 40S subunits without a concomitant reduction in free 60S subunits, the appearance of half-mers, and an overall decrease in polysome content, these alterations were, however, not perceptibly more pronounced than in Δ*rpl1b* single mutant cells (Supplementary Fig. S14B). We conclude that Bcl1 has some functional relevance when Rpl1 is present in lower amounts, especially at elevated growth temperatures, but its absence has clearly a lesser impact than the one of Acl1.

**Figure 8.**
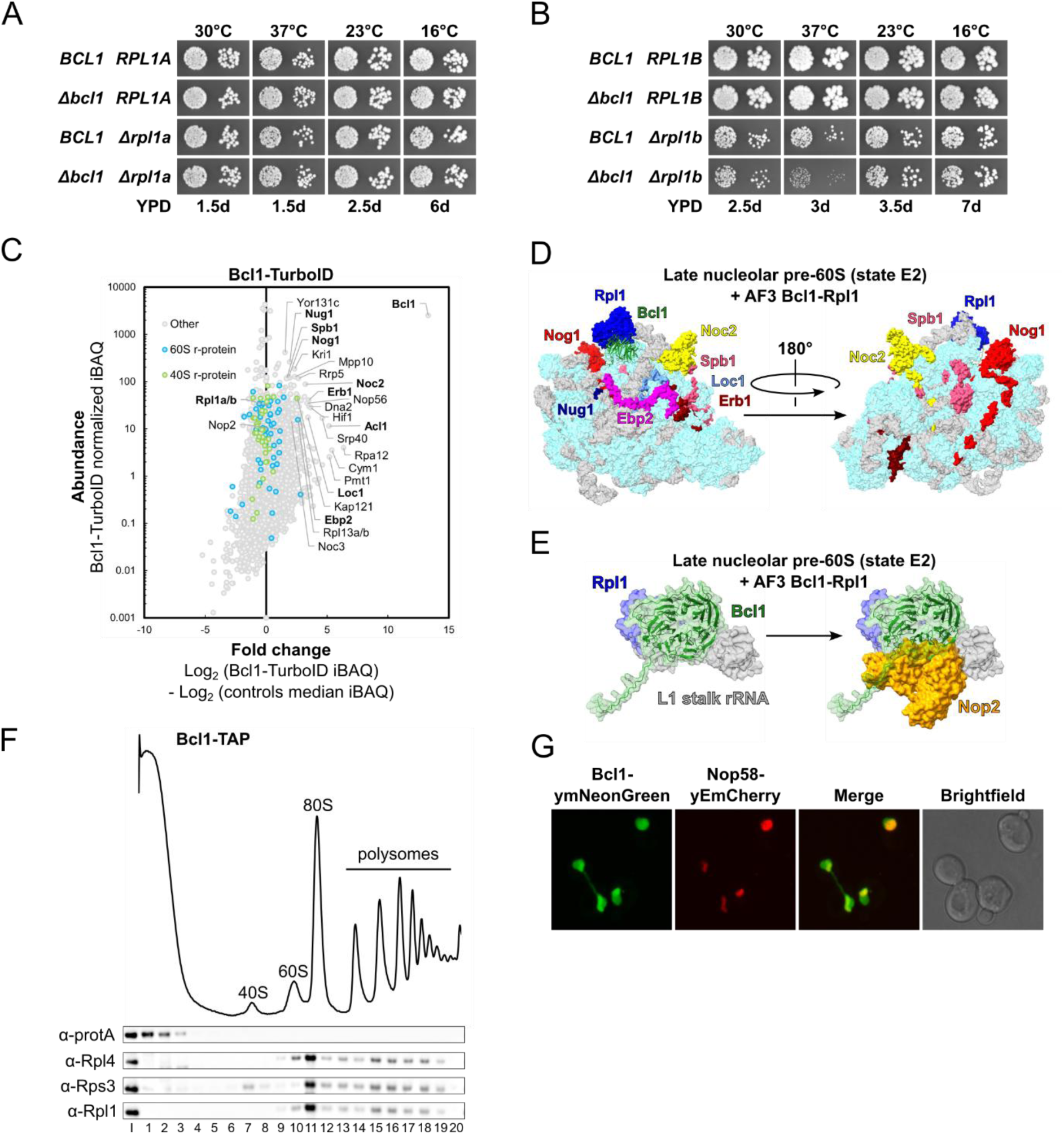
Bcl1 may transiently interact with nucleolar pre-60S particles. (**A, B)** Genetic interactions between Δ*bcl1* and Δ*rpl1a* **(A)** or Δ*rpl1b* **(B)**. Strains with the indicated genotypes, derived from tetratype tetrads, were spotted in 10-fold serial dilution steps onto YPD plates, which were incubated for the indicated times at 30, 37, 23, and 16°C. **(C)** TurboID results obtained with Bcl1-TurboID. The bait protein and selected enriched proteins are written in bold. **(D, E)** Superposition of the AlphaFold3 model of the Bcl1-Rpl1 complex onto Rpl1 within the late nucleolar state E2 pre-60S particle (PDB: 7NAC [98]). Bcl1-proximal proteins in the neighbourhood of the superimposed Bcl1-Rpl1 complex are indicated and highlighted in different colours, other r-proteins and AFs are coloured in cyan and (pre-)rRNAs in light grey **(D)**. Close-up view of the L1 stalk rRNA and the superimposed Bcl1-Rpl1 complex without (left) or with (right) showing Nop2 **(E)**. **(F)** A *BCL1*-TAP strain, expressing C-terminally TAP-tagged Bcl1 from its genomic locus, was grown at 30°C in YPD medium. The whole cell lysate was prepared under polysome-preserving conditions and analysed by sucrose gradient centrifugation and fractionation as described in the legend of Fig. 4F. **(G)** The subcellular localization of Bcl1-ymNeonGreen, expressed from plasmid under the control of its cognate promotor in a Δ*bcl1* null mutant strain, was assessed by fluorescence microscopy in cells grown at 30°C in SC-Ade-Leu medium. The subcellular position of the nucleolus was revealed by the nucleolar marker protein Nop58-yEmCherry, which was expressed from a plasmid under the control of its cognate promoter.

To gain insight into the functional environment of Bcl1, we next performed proximity labelling with C-terminally TurboID-tagged Bcl1. In agreement with the substantial enrichment of Bcl1 in the Acl1 TurboID assays (Fig. 3B), Acl1 was prominently enriched in the reciprocal proximity labelling with the Bcl1-TurboID bait (Fig. 8C). Despite being a direct interaction partner of Bcl1 (Fig. 7E and H), and in contrast to the Acl1 TurboID assays, Rpl1 was, however, not enriched in the Bcl1 TurboID assay (Fig. 8C). The reason for this is not clear, but the long length (67 residues) and/or a preferential orientation of Bcl1’s C-terminal extension, combined with the predicted shielding of multiple surface-exposed lysine residues when Rpl1 is bound by Bcl1 and, even more so, when it could be simultaneously bound by both Bcl1 and Acl1 (Fig. 7D and Supplementary Fig. S15), might preclude an efficient biotinylation of Rpl1. Interestingly, the proximity labelling assay with C-terminally TurboID-tagged Bcl1 revealed that several pre-60S AFs, particularly Ebp2, Erb1, Loc1, Noc2, Nog1, Nug1, and Spb1, were among the abundantly present and discernibly enriched proteins (Fig. 8C). Notably, when superposing the predicted Bcl1-Rpl1 complex onto Rpl1 in the structure of the late nucleolar state E2 pre-60S intermediate [98], the visualized parts of all the above-mentioned AFs cluster around Bcl1 (Fig. 8D and Supplementary Fig. S14D). Moreover, and consistent with all Rpl1 residues, except maybe Lys130, predicted to interact with Bcl1 not being in direct contact with the L1 stalk (Supplementary Fig. S13B and C), an interaction of Bcl1 with Rpl1 does not appear to be precluded, at least at this late nucleolar pre-60S maturation stage, by the association of Rpl1 with its rRNA-binding sites (Fig. 8E). However, the superposition indicates that β-propeller blades 1 to 3 of Bcl1 would clash with Nop2 (Fig. 8E), suggesting that Bcl1 must dissociate from Rpl1 prior to the incorporation of Nop2 into nucleolar pre-60S particles. In line with the rather moderate enrichment of the above-mentioned proximal pre-60S AFs and, thus, a potentially transient association of Bcl1 with nucleolar pre-60S particles, sucrose gradient fractionation of total extracts from cells genomically expressing C-terminally TAP-tagged Bcl1 revealed that Bcl1 was strongly enriched in the soluble fractions (Fig. 8F). Finally, fluorescence microscopy revealed that C-terminally ymNeonGreen-tagged Bcl1, expressed from a centromeric plasmid under the control of its cognate promoter in Δ*bcl1* cells, localized to the nucleus, where it was enriched, as indicated by its prevalent co-localization with the nucleolar marker protein Nop58-yEmCherry, in the nucleolus (Fig. 8G). Thus, the steady-state localization of Bcl1 is fully consistent with the observed enrichment of nucleolar pre-60S AFs in the proximity labelling assay, altogether strongly suggesting a functional role of Bcl1 in the nucleolus. Moreover, our data indicate that Bcl1, like most of the known DCs (Supplementary Fig. S8), very likely transiently interacts with pre-ribosomal particles and that, besides directly interacting with Rpl1, it could also interact with Acl1 or form a trimeric complex with Rpl1 and Acl1.

To explore these possibilities, we next investigated whether these dimeric or trimeric complexes can be reconstituted in vitro. In line with AlphaFold not predicting the formation of a binary Bcl1-Acl1 complex (data not shown, see also below), C-terminally Flag-tagged Acl1 could, unlike Rpl1 (Fig. 7H), not be co-purified by Ni-NTA affinity purification with C-terminally (His)_6_-tagged Bcl1.N366 when the two proteins were co-expressed in *E. coli* (Fig. 9A). However, upon inclusion of Rpl1 by co-expressing all three proteins together in *E. coli* (Rpl1b and Acl1-Flag from the same plasmid and Bcl1.N366-(His)_6_ from a separate plasmid), the trimeric Bcl1.N366-Rpl1-Acl1 complex could be purified by consecutive Ni-NTA and anti-Flag affinity purification (Fig. 9B). In support of this biochemical evidence, AlphaFold3 also predicted the formation of a trimeric complex (Fig. 9C), with the two dimeric modules (Bcl1-Rpl1 and Rpl1-Acl1) exhibiting a mostly identical overall structure and interaction mode when compared to their individually predicted co-structures (Supplementary Fig. S15A). Moreover, no direct contacts between Bcl1 and Acl1 could be observed in the predicted structure of the trimeric complex. Notably, the AlphaFold3 predictions of the trimeric complex as well as of the dimeric Acl1-Rpl1 complex also revealed a potential second Acl1-Rpl1 interaction interface. In addition to the N-terminal ankyrin repeat domain, which according to our Y2H, biochemical, and structural data is a fully sufficient and essential Rpl1-binding determinant (see Figs. 5 and 6), residues of the three-helix fold (starting at residue 145 and ending approximately at residue 205) within Acl1’s C-terminal half (residues 127-222) would also engage in predicted interactions with Rpl1 (Supplementary Figs. S15A and S16B). In line with this possibility, Y2H assays revealed a weak interaction between Acl1’s predicted three-helix fold and full-length Rpl1 (Supplementary Fig. S16C). Interestingly, while the ankyrin repeat domain recognizes a short Rpl1 segment that partially overlaps with the equally short rRNA-binding segment of the D2 domain (Fig. 6A and C and Supplementary Fig. S13B and C), the predicted interaction-mediating surface of the three-helix fold would get into contact with the rRNA-binding surface of the D1 domain, which almost exclusively forms interactions with helix H77 of the L1 stalk [98,188] (Supplementary Fig. S13B and C). Thus, by interacting with Rpl1 in this manner, Acl1 would effectively occlude both rRNA-binding surfaces of Rpl1 (Supplementary Fig. S16A). Additionally, by embedding Rpl1 in the trimeric complex, the majority of its positively charged surfaces would be efficiently shielded by Acl1 and Bcl1 (Fig. 9D and Supplementary Fig. S15B and C). Taken together, the so far presented data, including the above biochemical experiments and AlphaFold predictions, provide strong evidence that Rpl1, by employing distinct surfaces to independently interact with its two binding partners, can form a trimeric complex with Acl1 and Bcl1. Accordingly, the reciprocal enrichment of Bcl1 and Acl1 in the proximity labelling assays with the Acl1 and Bcl1 baits can only be interpreted in such a way that the trimeric complex must also occur in vivo (Figs. 3B and 8C). To obtain additional evidence for the in vivo formation of the trimeric complex, we performed Y2H assays between Bcl1 and Acl1, as we hypothesised that endogenously expressed Rpl1 might be able to bridge an interaction of the two proteins at the promoters of the reporter genes. Consistent with this assumption, a Y2H interaction between Bcl1 and Acl1 could be observed (Fig. 9E), albeit for only two of the eight combinations, i.e., when both proteins were tagged at their C-terminal end, and somewhat weaker than in the case of their respective interaction with Rpl1 (Figs. 4A and 7E). In support of their Y2H interaction indeed being bridged by Rpl1, Bcl1 no longer interacted with Acl1 variants exhibiting a visibly reduced (Y47A, D70A, and D72A) or no (D34A and D68A) Rpl1-binding capacity, and, moreover, showed a strongly reduced interaction with Acl1 variants being only mildly (E79A) or barely (N4A) affected in their ability to interact with Rpl1 (Fig. 9F and Supplementary Fig. S16D). Moreover, Bcl1 still interacted to a largely unchanged extent with Rpl1 variants that showed no or a much weaker Y2H interaction with Acl1 (Fig. 9G and Supplementary Fig. S16E), indicating that the in vivo interaction between Bcl1 and Rpl1 is not necessarily dependent on Acl1 and, thus, could in principle also occur outside the trimeric complex in the normal cellular context. We conclude that Rpl1, Acl1, and Bcl1 can form, both upon their co-overexpression in bacteria and according to AlphaFold predictions, a trimeric complex that, based on several lines of evidence, also occurs in yeast cells, at least for a certain duration along Rpl1’s journey to its assembly site on nucleolar pre-60S particles. Moreover, the Y2H interaction between Bcl1 and Rpl1 variants deficient in Acl1 binding raises the possibility that not only Acl1, as strongly suggested by its prominent enrichment in the Rpl1 TurboIDs and its Y2H interaction with Rpl1 variants deficient in Bcl1 binding (Figs. 3A and 7G), but also Bcl1 could possibly form a binary complex with Rpl1 in vivo.

**Figure 9.**
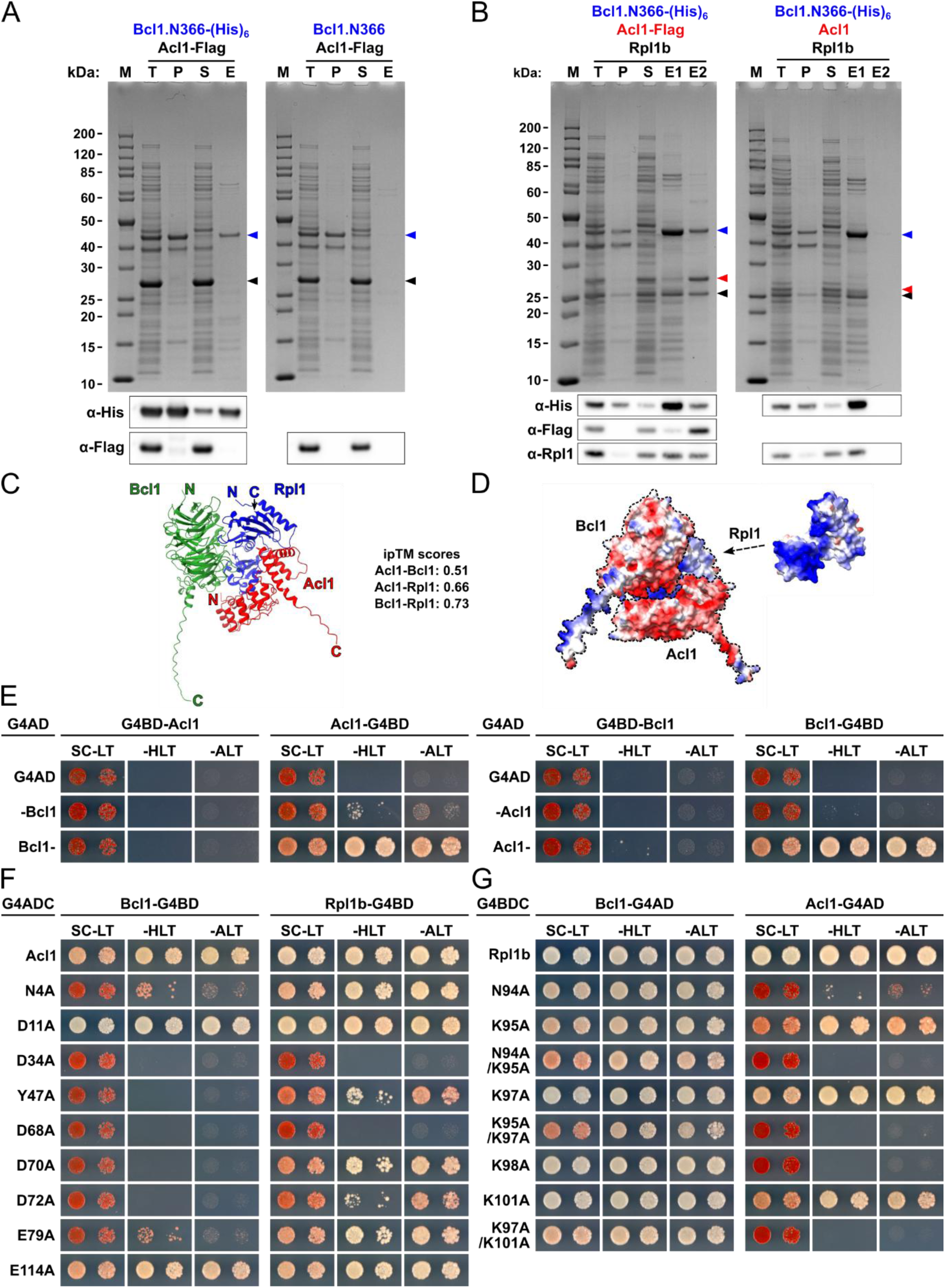
Acl1, Bcl1, and Rpl1 can form a trimeric complex. **(A)** In vitro binding assay between Bcl1.N366 and Acl1. Bcl1.N366-(His)_6_ or Bcl1.N366 and Acl1-Flag were co-expressed in *E. coli* and purified by Ni-NTA affinity purification. Proteins were revealed by SDS-PAGE and Coomassie staining (top) or by Western blotting using anti-His and anti-Flag antibodies (bottom). Bands corresponding to Bcl1.N366-(His)_6_ and Bcl1.N366 or to Acl1-Flag are indicated by blue or black arrowheads. M: molecular weight standard, T: total extract, P: pellet fraction, S: soluble extract, E: imidazole eluate. **(B)** Purification of the trimeric Acl1-Rpl1-Bcl1 complex. Bcl1.N366-(His)_6_, Rpl1b, and either Acl1-Flag (left) or untagged Acl1 (right) were co-expressed in *E. coli* and purified by consecutive Ni-NTA and anti-Flag affinity purification. Proteins were revealed by SDS-PAGE and Coomassie staining (top) or by Western blotting using anti-His, anti-Flag, and anti-Rpl1 antibodies (bottom). Bands corresponding to Bcl1.N366-(His)_6_, Rpl1b, and Acl1-Flag or Acl1 are indicated by blue, black, or red arrowheads. M: molecular weight standard, T: total extract, P: pellet fraction, S: soluble extract, E1: imidazole eluate, E2: Flag eluate. **(C, D)** Cartoon representation **(C)** and predicted electrostatic surface potential with the protected positively charged surface of Rpl1 being highlighted **(D)** of the AlphaFold3 model of the trimeric Acl1-Rpl1-Bcl1 complex. The ipTM scores of the respective binary interactions are indicated. **(E-G)** Y2H interaction assays between the full-length Acl1 and Bcl1 proteins **(E)**, between the indicated Acl1 mutant variants and either Bcl1 or Rpl1 **(F)**, and between the indicated Rpl1 mutant variants and either Bcl1 or Acl1 **(G)**. Single-letter abbreviations for the amino acid residues are as follows: A, Ala; D, Asp; E, Glu; K, Lys; N, Asn; Y, Tyr.

**Figure 10.**
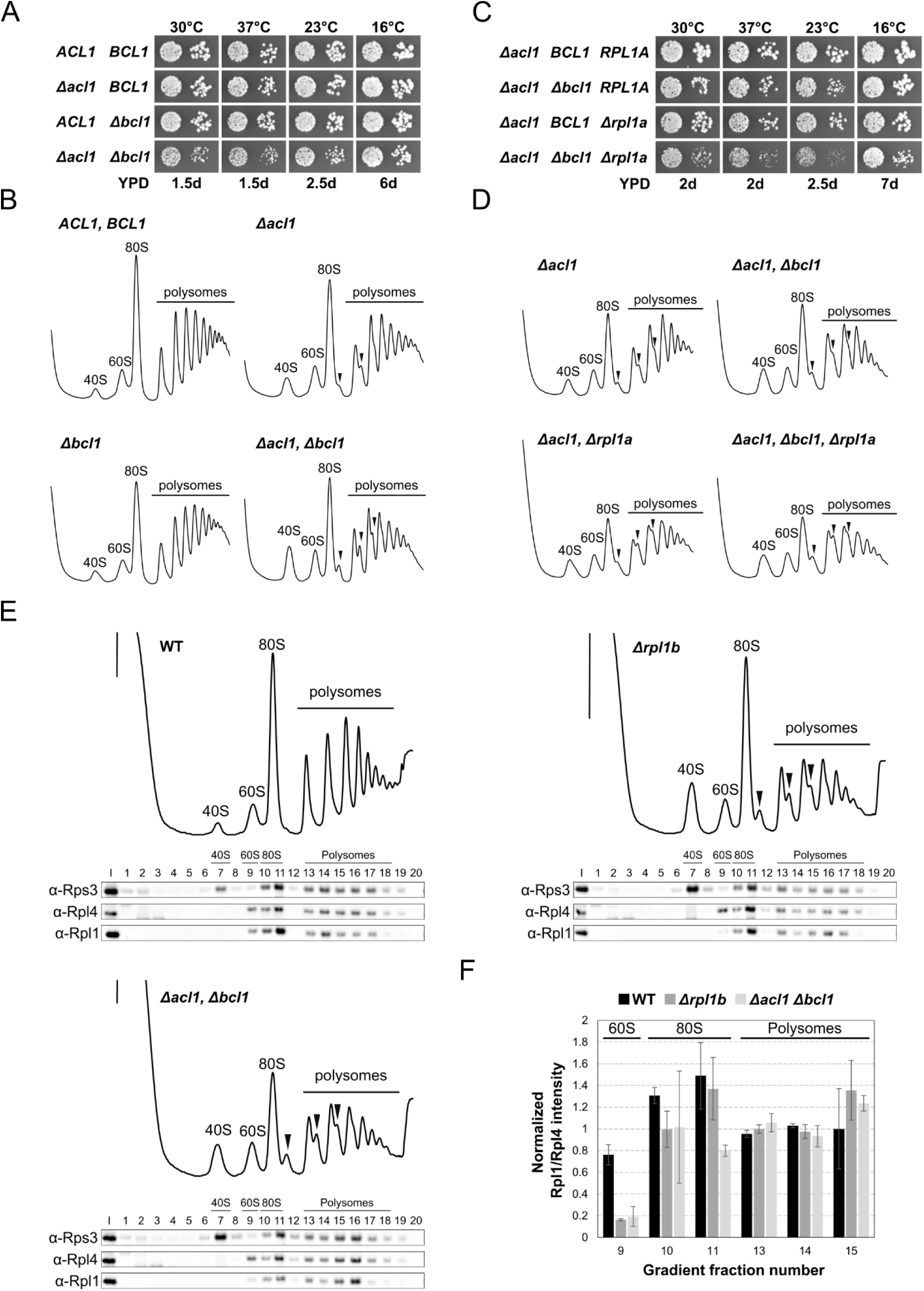
Acl1 and Bcl1 functionally cooperate to ensure efficient incorporation of Rpl1. (**A, C)** Negative genetic interactions between Δ*acl1* and Δ*bcl1* **(A)** or between Δ*acl1*, Δ*bcl1*, and Δ*rpl1a* **(C)**. Strains with the indicated genotypes, derived from tetratype tetrads, were spotted in 10-fold serial dilution steps onto YPD plates, which were incubated for the indicated times at 30, 37, 23, and 16°C. **(B, D)** Polysome profiles of wild-type (*ACL1*, *BCL1*), Δ*acl1*, Δ*bcl1*, and Δ*acl1*/Δ*bcl1* cells **(B)**, and of Δ*acl1*, Δ*acl1*/Δ*bcl1*, Δ*acl1*/Δ*rpl1a*, and Δ*acl1*/Δ*bcl1*/Δ*rpl1a* cells **(D)** grown at 30°C in YPD medium. **(E)** Whole cell lysates of wild-type (WT), Δ*rpl1b*, and Δ*acl1*/Δ*bcl1* strains, grown at 30°C in YPD medium, were prepared under polysome-preserving conditions and analysed by sucrose gradient centrifugation and fractionation as described in the legend of Fig. 4F. An input (I) control (0.05 A_260_ units of the clarified lysate) and the gradient fractions (1-20) were analysed by Western blotting using anti-Rps3, anti-Rpl4, and anti-Rpl1 antibodies. **(F)** Bar graph showing the average and standard deviation (n=2) of the normalized Rpl1/Rpl4 ratios in the 60S (9), 80S (10, 11), and polysomal (13-15) fractions of the indicated strains.

Among the r-proteins associated with DCs, Rpl1 is a special case in that it transiently interacts, most likely for a certain duration simultaneously, with two distinct binding partners. To explore the functional relevance of this particular setting, we next assessed the effects of the simultaneous absence of Acl1 and Bcl1 on growth, synthesis of 60S subunits, and the assembly of Rpl1. Consistent with a functional synergism, Δ*acl1*/Δ*bcl1* double mutant cells showed a clearly discernible growth defect at all tested temperatures compared to the virtually normal growth of Δ*acl1* and Δ*bcl1* single mutant cells (Fig. 10A). In line with the observed growth defect, polysome profile analysis revealed a clear decrease in polysome content in Δ*acl1*/Δ*bcl1* double mutant cells; moreover, the accumulation of free 40S subunits and half-mer polysomes was more pronounced than in Δ*acl1* single mutant cells (Fig. 10B). However, as in the case of Δ*acl1*/Δ*rpl1a* double mutant cells (Fig. 4D), the size of the 60S peak was not reduced in Δ*acl1*/Δ*bcl1* double mutant cells, indicating that the formation and stability of pre-60S particles is not substantially reduced and, hence, that a fraction of the produced 60S subunits must rather be altered in a manner that renders pre-60S export or subunit joining less efficient. Considering that the absence of Rpl1a already negatively affected the growth of Δ*acl1* single mutant cells (Fig. 4C), we next asked whether this slight reduction in Rpl1 levels would also have an additional impact on the growth of Δ*acl1*/Δ*bcl1* double mutant cells. As expected, Δ*acl1*/Δ*bcl1*/Δ*rpl1a* triple mutant cells grew substantially slower than Δ*acl1*/Δ*bcl1* and Δ*acl1*/Δ*rpl1a* double mutant cells (Fig. 10C). Moreover, this further decline in growth rate was accompanied by an additional decrease in polysome content (Fig. 10D). As a previous study had shown that 60S subunits lacking Rpl1 can be observed in Δ*rpl1b* mutant cells [184], i.e., when Rpl1 production is substantially reduced [72,102], we next assessed whether simultaneous absence of Acl1 and Bcl1 would also result in the formation of 60S subunits that do not contain Rpl1. To this end, we fractionated total extracts, prepared under polysome-preserving conditions by inhibiting translation elongation with cycloheximide, from wild-type, Δ*rpl1b*, and Δ*acl1*/Δ*bcl1* cells by sucrose gradient centrifugation and assessed the gradient localization and abundance of Rpl1 and the LSU and SSU control r-proteins Rpl4 and Rps3 by Western blotting (Fig. 10E and Supplementary Fig. S17). Upon quantification of the signal intensities of Rpl1 and Rpl4 in each fraction, we first separately calculated the relative abundance of Rpl1 and Rpl4 in the individual fractions by dividing their signal intensity in these fractions by their average signal intensity in two polysomal fractions (fraction 13 and 14), assuming that 60S subunits within translating 80S ribosomes not only obligatorily contain Rpl4, whose incorporation is essential for the formation of 60S subunits [34,201], but should also be equipped with Rpl1 in all three strains, as previously indicated by the similar relative levels of Rpl1 in the polysome fractions of wild-type and Δ*rpl1b* mutant cells [184]. Then, we formed the ratio between the above-determined normalized abundance values of Rpl1 and Rpl4 in the individual fractions to assess the proportion of 60S subunits lacking Rpl1 in the different fractions of our two experimental replicates. Surprisingly, by applying this quantification strategy, we observed that only around 75% of the 60S subunits contained Rpl1 in the wild-type strain (Fig. 10F). In the Δ*rpl1b* mutant strain, the proportion of 60S subunits containing Rpl1 decreased to slightly below 20%, which, after normalization against the above wild-type value, agrees well with the previously reported reduction in Rpl1 presence on 60S subunits to around 30% of wild-type levels [184]. Notably, cells simultaneously lacking Acl1 and Bcl1 (Δ*acl1*/Δ*bcl1* double mutant) exhibited a similar decrease in the proportion of 60S subunits containing Rpl1 as Δ*rpl1b* mutant cells, indicating that the two binding partners of Rpl1 cooperatively ensure that Rpl1 is provided in sufficient amounts and/or gets efficiently incorporated into pre-60S particles. Taken together, we conclude that the joint function of Acl1 and Bcl1, which together, by embedding Rpl1 within a trimeric complex, are predicted to efficiently shield almost all positively charged surfaces of Rpl1, is related to the provisioning of assembly-competent Rpl1.

### Acl1 and Bcl1 contain functional NLSs and mediate nuclear import of Rpl1

The nuclear import of highly basic proteins, such as histones and r-proteins, has been shown to rely, despite their small size, on active, importin-mediated transport processes [9,28,202,203]. However, as none of the different importins was particularly enriched in the Rpl1b TurboID assays and as a cytoplasmically formed trimeric complex, owing to its size (calculated molecular mass of 93 kDa) exceeding the limit for efficient passive diffusion across the selective phase of the NPC [204,205], would probably be precluded from efficiently entering the nucleus, we wondered whether Acl1 and/or Bcl1 might be involved in mediating the nuclear import of Rpl1. Consistent with this possibility, inspection of the primary sequence of Acl1 and Bcl1, which are both predominantly located in the nucleus in exponentially growing yeast cells (Figs. 4E and 8G), revealed that both proteins harbour a potential NLS. Acl1 contains at its C-terminal extremity four consecutive basic residues (KRRK) (Fig. 11A), which, as also predicted by PSORT II [206], correspond to a classical monopartite NLS bearing the loose K-K/R-x-K/R consensus sequence that is expected to be recognized by the importin-α Kap60 [207]. In line with this, AlphaFold3 predictions of the putative pentameric Kap95-Kap60-Acl1-Rpl1-Bcl1 import complex indicated that these positively charged residues would interact in a canonical manner with the major NLS-binding site of Kap60 [133] (Supplementary Fig. S18A), suggesting that the transport adaptor Kap60, in conjunction with the importin-β Kap95, could be responsible for the nuclear import of Acl1 as well as of the dimeric Acl1-Rpl1 and the trimeric Acl1-Rpl1-Bcl1 complex. Bcl1 contains, like the validated Kap104 import cargo Hrp1 [208–210], a basic PY-NLS (bPY-NLS) at its C-terminal extremity (Fig. 11B). Consistent with matching well the consensus sequence (basic-enriched stretch that is followed by a C-terminal R/K/H-x_(2-5)_-P-Y/ϕsignature, with ϕdenoting a hydrophobic residue) [152,164], AlphaFold3 predicted with good confidence that the bPY-NLS of Bcl1 (residues 371-389) would be accommodated by the importin-β Kap104 in a similar manner as bPY-NLSs in co-structures with the human Kap104 equivalent Kapβ2/TNPO1 [211,212] (Supplementary Fig. S18C). Moreover, AlphaFold3 also predicted the formation of the putative tetrameric Kap104-Bcl1-Rpl1-Acl1 import complex (Supplementary Fig. S18B), in which the two modules, i.e., the dimeric Kap104-Bcl1 complex and the trimeric Acl1-Rpl1-Bcl1 complex, retained their overall structure and interaction mode when compared to their individually predicted co-structures. In further support of Kap104 being a strong candidate importin for mediating the nuclear import of Bcl1 and, thus, possibly also of the dimeric Bcl1-Rpl1 and the trimeric Acl1-Rpl1-Bcl1 complex, Bcl1 was, like the known bPY-NLS-containing Kap104 cargos Hrp1, Nab2, Syo1, and Tfg2 [46,209,210,213,214], previously detected by LC-MS/MS in a one-step affinity purification of Flag-tagged Kap104 [215].

**Figure 11.**
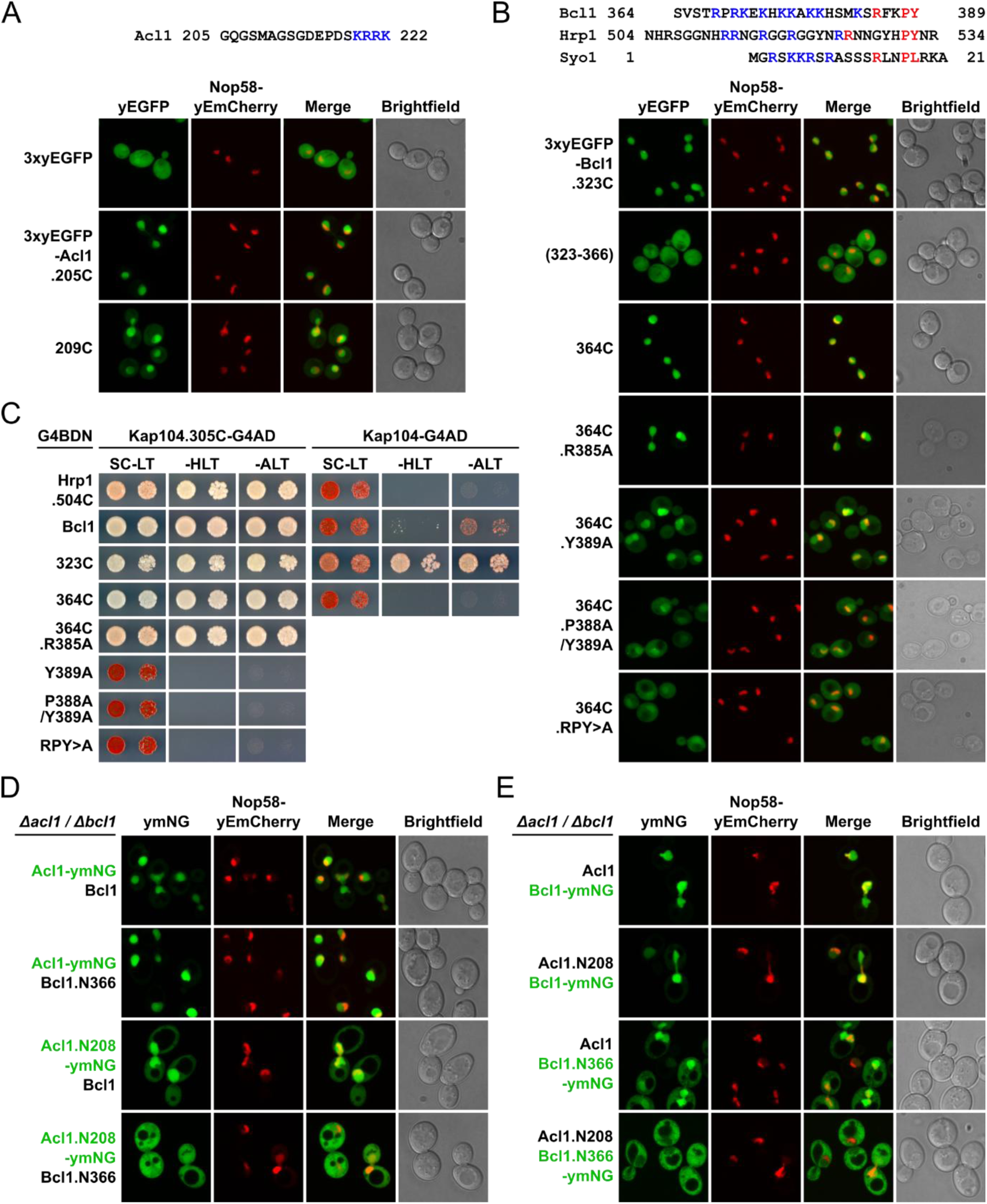
Acl1 and Bcl1 contain functional NLSs and can be co-imported as a trimeric Acl1-Rpl1-Bcl1 complex. (**A, B)** The C-terminal extremity of Acl1 contains a predicted classical NLS **(A, upper panel)**, and the C-terminal extremity of Bcl1 resembles a bPY-NLS, as illustrated by its comparison with the known bPY-NLSs of Hrp1 and Syo1 **(B, upper panel)**. Basic residues are coloured in blue and residues of the characteristic C-terminal signature of PY-NLSs in red. **(A, B, lower panels)** The nuclear targeting activity, as inferred from the capacity to confer nuclear localization to an N-terminally fused 3xyEGFP reporter, of C-terminal Acl1 segments (205C, residues 205-222; 209C, residues 209-222) **(A, lower panels)**, and of the entire C-terminal extension of Bcl1 (323C, residues 323-389), the extension lacking the terminal bPY-NLS region (residues 323-366), and the bPY-NLS-containing C-terminal extremity of Bcl1 (364C, residues 364-389) as well as the indicated mutant variants thereof **(B, lower panels)** was assessed by fluorescence microscopy in cells grown at 30°C in SC-Leu medium. The 3xyEGFP control protein and the N-terminally 3xyEGFP-tagged fusion proteins were expressed from plasmid under the control of the *ADH1* promoter in cells expressing the nucleolar marker protein Nop58-yEmCherry from the genomic locus. **(C)** Y2H interaction assays between the N-terminally G4BD-tagged (G4BDN) bPY-NLS-containing C-terminal extremity of Hrp1 (Hrp1.504C), full-length Bcl1, Bcl1.323C, and Bcl1.364C, as well as the indicated mutant variants thereof, and C-terminally G4AD-tagged full-length Kap104 (right) or the N-terminally truncated Kap104.305C variant (left). Single-letter abbreviations for the amino acid residues are as follows: A, Ala; P, Pro; R, Arg; Y, Tyr. **(D, E)** The subcellular localization of C-terminally ymNeonGreen-tagged Acl1 (Acl1-ymNG) or its C-terminally truncated, NLS-lacking variant (Acl1.N208-ymNG) **(D)** and of C-terminally ymNeonGreen-tagged Bcl1 (Bcl1-ymNG) or its C-terminally truncated, NLS-lacking variant (Bcl1.N366-ymNG) **(E)** was assessed by fluorescence microscopy in Δ*acl1*/Δ*bcl1* cells, grown at 30°C in SC-Ade-Leu-Trp medium, that simultaneously expressed Bcl1 or Bcl1.N366 **(D)** and, respectively, Acl1 or Acl1.N208 **(E)**. The ymNeonGreen-tagged and untagged proteins were expressed from plasmid under the control of the cognate *ACL1* or *BCL1* promoter. The subcellular position of the nucleolus was revealed by the nucleolar marker protein Nop58-yEmCherry, which was expressed from plasmid under the control of its cognate promoter.

To experimentally address whether the C-terminal segments of Acl1 and Bcl1 carrying the supposed NLS sequences would indeed correspond to functional NLSs in vivo, we assessed their capacity to target an N-terminally fused triple yEGFP reporter (3xyEGFP) to the nucleus by fluorescence microscopy. In line with the four basic residues at the C-terminal extremity of Acl1 being able to mediate nuclear import, the 3xyEGFP reporter fused to residues 205-222 (Acl1.205C) or 209-222 (Acl1.209C) was exclusively or, respectively, predominantly present in the nucleus (Fig. 11A). Similarly, both the entire C-terminal extension (residues 323-389, Bcl1.323C) of Bcl1 and a shorter, 26-residue-long segment (residues 364-389, Bcl1.364C), essentially consisting of the predicted bPY-NLS, were sufficient to confer exclusive nuclear localization to the 3xyEGFP reporter, whereas the C-terminal extension lacking the bPY-NLS did not exhibit nuclear targeting activity (Fig. 11B).

To obtain experimental evidence for a direct interaction between the predicted bPY-NLS of Bcl1 and Kap104, we intended to employ, as already before for the validation of interactions between importin-βs and r-proteins (see Fig. 2), the Y2H method. To this end, we generated the Kap104.305C variant, which, akin to the previously successfully used Kap121.302C variant (Fig. 2A), lacks HEAT repeats 1 to 7 and, thus, the majority of the RanGTP-interacting residues of Kapβ2/TNPO1 [142,154], but retains all Kap104 residues predicted to interact with Bcl1 (Supplementary Fig. S18C). As hoped, Y2H assays between the N-terminally G4BD-tagged bPY-NLS-containing segment of Hrp1 (residues 504-534, Hrp1.504C) and the full-length Kap104 and Kap104.305C prey proteins, fused at their C-terminal end to the G4AD, permitted to recapitulate the previously reported RanGTP-sensitive in vitro interaction between Kap104 and a C-terminal Hrp1 segment starting at residue 494 [210] (Fig. 11C). Consistent with Bcl1 being an import cargo of Kap104, the short bPY-NLS-containing segment of Bcl1 (Bcl1.364C) exhibited a strong Y2H interaction with the Kap104.305C variant but did not interact with full-length Kap104 (Fig. 11C). Similarly, also full-length Bcl1 interacted strongly with the Kap104.305C variant, while still showing a weak residual interaction with full-length Kap104. Intriguingly, however, the entire C-terminal extension of Bcl1 (Bcl1.323C), while, as expected, robustly interacting with the Kap104.305C variant, still displayed a moderately strong Y2H interaction with full-length Kap104. The reason for this is unclear, but we speculate that the segment preceding the bPY-NLS might somehow reduce the efficiency by which the Y2H interaction between Kap104 and the complete C-terminal extension of Bcl1 is disrupted by endogenous GTP-loaded Gsp1. To examine the relevance of the characteristic R-x_(2)_-P-Y signature for the interaction, we evaluated the effects of the individual or combined alanine substitution of these residues on the Y2H interaction. While the R385A substitution in the context of the complete C-terminal extension or the short bPY-NLS-containing segment had no or, respectively, only a marginal effect on the Y2H interaction with the Kap104.305C variant, it abolished the interaction between the complete C-terminal extension and full-length Kap104 (Fig. 11C and Supplementary Fig. S18D). Strikingly, the single Y389A exchange, disrupting the predicted hydrogen bond between the tyrosyl of the C-terminal Tyr389 residue and the carboxyl group of residue Asp408 of Kap104 (Supplementary Fig. S18C), was already sufficient to abrogate, both in the context of the entire and short C-terminal region, the Y2H interaction with the Kap104.305C variant (Fig. 11C and Supplementary Fig. S18D); accordingly, also the simultaneous P388A/Y389A double and R385A/P388A/Y389A triple substitutions abolished the interaction. These Y2H results are reminiscent of previously reported binding affinity measurements between mutant bPY-NLS variants of Hrp1 and full-length Kap104, which showed that the introduction of the Y532A and P531A/Y532A substitutions resulted in no detectable in vitro binding, whereas the R525A exchange did not affect the binding capacity of Hrp1’s bPY-NLS [210]; thus, further substantiating the relevance of the tyrosine residue within the PY motif for the interaction with Kap104 in the case of C-terminally located bPY-NLSs.

Next, we determined whether these amino acid substitutions would also affect the nuclear targeting activity of the complete C-terminal extension or the short bPY-NLS-containing segment. In line with the Y2H interaction data, the R385A substitution did not alter the exclusive nuclear accumulation of the N-terminally fused 3xyEGFP reporter (Fig. 11B and Supplementary Fig. S18E). Surprisingly, however, the Y389A, P388A/Y389A, and R385A/P388A/Y389A exchanges, both in the Bcl1.323C and Bcl1.364C context, only resulted in a similar, partial relocalization of the 3xyEGFP reporter to the cytoplasm, showing that the nuclear targeting activity of the bPY-NLS-containing segment is not abolished by mutations that abrogate its Y2H interaction with the Kap104.305C variant. This observation either implies that this 26-residue-long segment, which is rich in positively charged amino acids (eight lysines and three arginines), can also be recognized and get imported by another importin than Kap104 or could indicate that the introduced amino acid substitutions do not completely prevent the formation of a productive import complex with full-length Kap104. A promising alternative importin was Kap121, as it was the only importin that was noteworthily enriched in the proximity labelling assay with C-terminally TurboID-tagged Bcl1 (Supplementary Fig. S18F). Consistent with their suggested physical proximity, the complete C-terminal extension of Bcl1, when fused at its N-terminal end to the G4BD, interacted quite well with the C-terminally G4AD-tagged Kap121.302C variant, but, and in line with Bcl1 being a potential import cargo of Kap121, it did not show a Y2H interaction with full-length Kap121 (Supplementary Fig. S18G). To our surprise, the shorter, 26-residue-long Bcl1.364C segment did, however, not interact with the Kap121.302C variant, indicating that Kap121 might not be the second importin contributing to its nuclear targeting when N-terminally fused to the 3xyEGFP reporter and, thus, that at least one further importin could potentially be involved in the import of Bcl1. However, it cannot be excluded that a steric constraint might selectively prevent the formation of the Y2H complex between Bcl1.364C and the N-terminally truncated Kap121.302C variant at the promoters of the reporter genes, while formation of the import complex with full-length Kap121 might still be possible. We conclude that Bcl1 harbours an NLS region at its C-terminal extremity that not only contains a bPY-NLS for Kap104-mediated import, but also very likely features at least one or possibly even two additional, yet-to-be-defined determinants that enable nuclear import, with one of them potentially being recognized by Kap121.

As Acl1 and Bcl1 contain functional NLSs and can form a trimeric complex with Rpl1 in vitro and in vivo, we next examined whether their nuclear import would show some interdependence. To this end, we expressed combinations of Acl1, Bcl1, and their C-terminally truncated variants lacking the NLS region (i.e., Acl1.N208 and Bcl1.N366), with one of the two being C-terminally tagged with ymNeonGreen, under the control of their cognate promoters from separate plasmids in Δ*acl1*/Δ*bcl1* cells. As expected, Acl1-ymNeonGreen and Bcl1-ymNeonGreen showed almost exclusive nuclear accumulation when co-expressed with either the full-length or NLS-lacking variant of Bcl1 and Acl1 (Fig. 11D and E). Notably, also ymNeonGreen-tagged Acl1.N208 and Bcl1.N366 exhibited, when co-expressed with full-length Bcl1 or Acl1, a predominant nuclear localization, suggesting that these NLS-lacking variants get, albeit slightly less efficiently, imported as part of the trimeric Acl1-Rpl1-Bcl1 complex. In line with this conjecture, no nuclear accumulation of ymNeonGreen-tagged Acl1.N208 and Bcl1.N366 could be observed when they were co-expressed together with Bcl1.N366 or Acl1.N208. These results strongly suggest that Acl1 and Bcl1 can form a trimeric Acl1-Rpl1-Bcl1 complex in the cytoplasm, whose nuclear import can be mediated by either one of the two C-terminal NLS regions.

To assess if Acl1 and Bcl1 are relevant for the nuclear import of Rpl1, we examined whether nuclear accumulation of Rpl1 can be observed in cells lacking both Acl1 and Bcl1 when pre-60S export is blocked. To this end, we determined the localization of C-terminally yEGFP-tagged Rpl1b, expressed from plasmid under the control of its cognate promoter, in wild-type and Δ*acl1*/Δ*bcl1* cells upon galactose-induced overexpression of a dominant-negative Nmd3 variant lacking the C-terminal nuclear export signal (NES) region (Nmd3.N418, residues 1-418; corresponding to the previously described Nmd3Δ100 variant [216,217]) (Fig. 12), which gets incorporated into pre-60S subunits and inhibits their nuclear export by disabling the interaction with the NES-recognizing exportin Crm1 [217]. As expected, overexpression of Nmd3.N418, induced for 8 h by the addition of galactose to the culture medium containing raffinose as carbon source, in wild-type cells resulted in a strong nuclear accumulation of C-terminally yEGFP-tagged Rpl25 (uL23), which is standardly used as a reporter for monitoring pre-60S export [217,218]. Importantly, Rpl1b-yEGFP showed a similarly strong nuclear accumulation under these conditions, revealing that it gets efficiently imported into the nucleus when Acl1 and Bcl1 are present. However, upon overexpression of Nmd3.N418 in cells lacking Acl1 and Bcl1, while Rpl25-yEGFP still displayed nuclear accumulation, Rpl1b-yEGFP no longer accumulated in the nucleus and even showed nuclear exclusion, strongly suggesting that Rpl1’s transfer into the nuclear compartment largely depends on Acl1 and Bcl1. Moreover, no nuclear accumulation of Rpl25-yEGFP could be observed in Δ*acl1*/Δ*bcl1* cells containing the empty vector or the plasmid expressing wild-type Nmd3, indicating that their absence, and thus the reduced incorporation of Rpl1 into pre-60S particles in the nucleus, does not confer a pre-60S export defect, at least when cells are grown in synthetic media with raffinose or galactose as carbon source.

**Figure 12.**
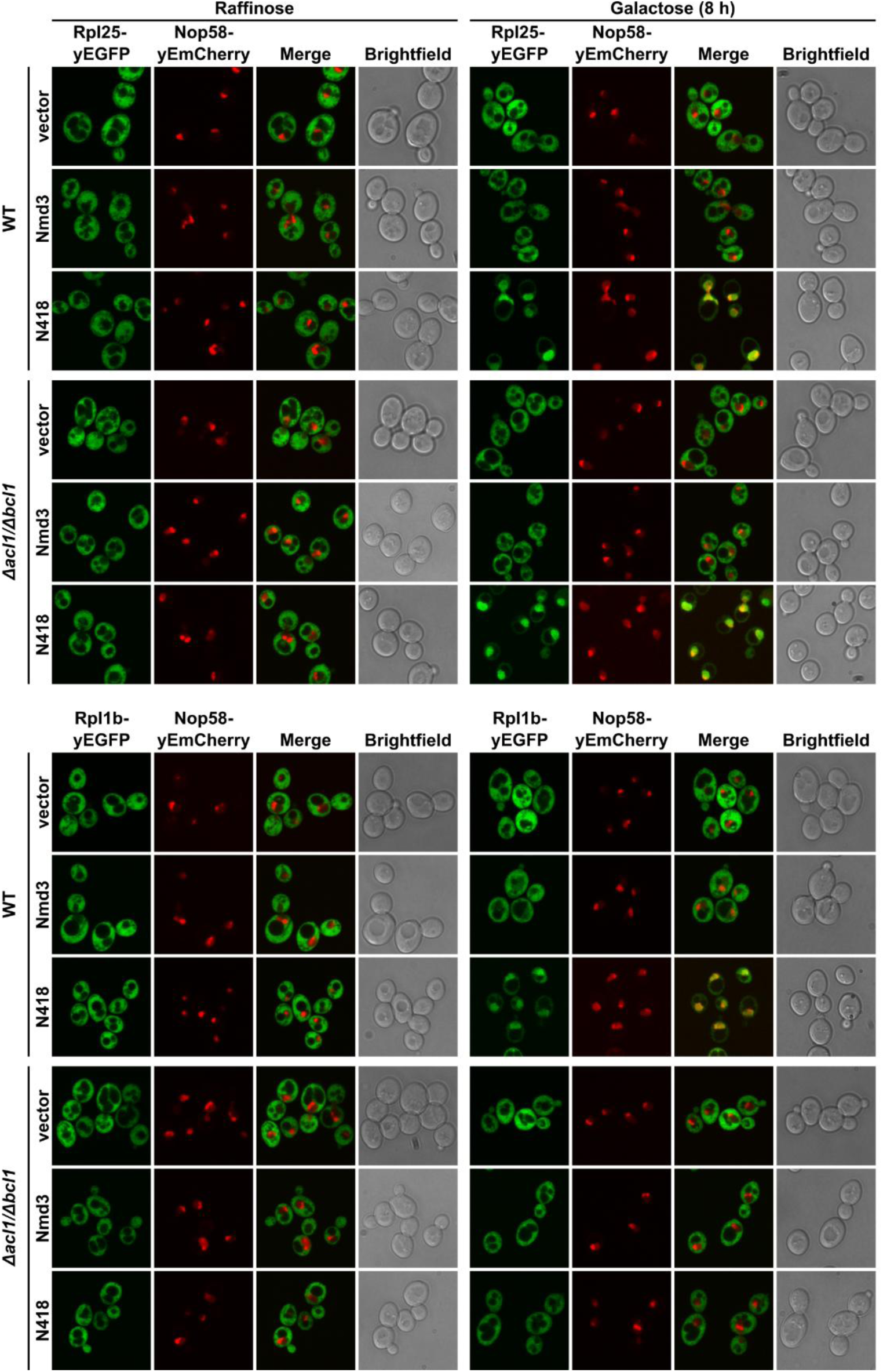
Acl1 and Bcl1 are required for the nuclear accumulation of Rpl1 when pre-60S export is impaired. Wild-type (WT) and Δ*acl1*/Δ*bcl1* cells were co-transformed with plasmids expressing C-terminally yEGFP-tagged Rpl25 or Rpl1b from their cognate promoters, Nmd3 or an Nmd3 variant lacking the C-terminal NES region (N418, residues 1-418) from the inducible *GAL1-10* promoter, and, to indicate the subcellular position of the nucleolus, Nop58-yEmCherry from its cognate promoter. Cells were grown in SRaf-Ade-Leu-Trp medium (Raffinose; left panels) and expression of Nmd3 or Nmd3.N418 was induced for 8 h with 2% galactose (Galactose; right panels). The subcellular localization of Rpl25-yEGFP (upper panels) and Rpl1b-yEGFP (lower panels) was assessed by fluorescence microscopy in cells that were maintained in exponential growth phase.

Finally, as a cytoplasmic recognition of Rpl1 by Acl1 and/or Bcl1 could already occur during its synthesis, we assessed the capacity of C-terminally TAP-tagged Acl1 and Bcl1 to co-translationally capture nascent Rpl1 by IgG-Sepharose pull-down and real-time qRT-PCR (see Materials and Methods). While purification of TAP-tagged Rrb1 and Acl4, as previously shown [34,41], specifically enriched the *RPL3* or *RPL4* mRNA, an enrichment of the mRNA encoding Rpl1 could neither be observed in the Acl1 nor the Bcl1 purification (Supplementary Fig. S19). Likewise, the experiment with both N-and C-terminally TAP-tagged Bcp1 did not reveal an enrichment of the *RPL23* mRNA. These results indicate that both Acl1 and Bcl1 most likely only interact with newly synthesized Rpl1 once it has been released from the ribosome.

Taken together, we conclude that Acl1 and Bcl1 are functionally cooperating DCs that both directly interact with Rpl1, can form a trimeric Acl1-Rpl1-Bcl1 complex, mediate the nuclear import of newly synthesized Rpl1, and enable a shielded and coordinated transfer of Rpl1 to its assembly site on nucleolar pre-60S particles.

## DISCUSSION

In this study, with the aim of expanding the inventory of DCs of r-proteins, we have performed a TurboID-based proximity labelling screen with all LSU r-proteins in the yeast *S. cerevisiae* (Fig. 1A and Supplementary Fig. S1). Despite certain potential limitations that are inherent to this proximity-dependent biotinylation approach, such as the availability of surface-exposed proximal lysine residues as biotinylation acceptor sites on the overall acidic DCs, the distance between the attached TurboID biotin ligase and the bound DC, and the productive complex formation between the tagged r-protein and its DC being hindered or prevented by the fused TurboID moiety, the majority (six out of eight) of the known DCs as well as one novel candidate DC could be reliably identified as prominently and selectively enriched proteins in the proximity labelling assays of their respective N– and/or C-terminally TurboID-tagged r-protein client (Supplementary Fig. S6; see also below). Importantly, analysis of multiple experimental replicates of the TurboID assays performed with the Rpl3-TurboID and TurboID-Rpl1b baits revealed high reproducibility (Fig. 1C and Supplementary Fig. S10A), indicating that the single replicate screen generated a robust and reliable data set. In further support of the suitability of the chosen approach and its experimental design (induced expression of the plasmid-encoded TurboID-tagged r-protein for 1 h with simultaneous induction of biotinylation by addition of exogenous biotin) for the identification of transient interaction partners of unassembled r-proteins, the TurboID-based proximity labelling screen also unveiled potential interactions between LSU r-proteins and importins, which we confirmed in the case of four r-proteins by Y2H assays (Fig. 2). Remarkably, even though designed to preferentially capture transient interactions of r-proteins prior to their assembly into pre-ribosomal particles, our approach also permitted to illuminate r-protein neighbourhoods on successive pre-ribosomal particles at spatiotemporal resolution, as exemplarily illustrated for the AFs and r-proteins that were reproducibly enriched in the proximity labelling assays with the Rpl3-TurboID and TurboID-Rpl1b baits (Fig. 1D-G and Supplementary Fig. S10B-H). Finally, the reciprocal TurboID assays with the already known DCs not only prominently enriched in each case the respective r-protein client but also revealed in most cases an enrichment of distinct AFs (Supplementary Fig. S8), which are notably either in proximity of their r-protein client on pre-ribosomal particles or have previously been functionally connected to their r-protein client, indicating that almost all DCs appear to be transiently associated with pre-ribosomal particles in order to promote the coordinated incorporation of their r-protein clients.

Notably, our proximity labelling screen enabled the identification of the fungi-specific ankyrin repeat-containing Acl1 as a transient interaction partner of Rpl1, which is the universally conserved r-protein associated with the flexible L1 stalk of the 60S subunit. Strikingly, and unlike the hitherto known DC-bound r-proteins, Rpl1 is also transiently associated with a second interaction partner, Bcl1, a predicted WD-repeat β-propeller protein that is conserved throughout eukaryotes. While Acl1 and Bcl1 are individually dispensable for optimal growth (Fig. 10A), several lines of evidence strongly suggest that they functionally cooperate to ensure a sufficient supply of assembly-competent Rpl1. First, their simultaneous absence results in a clearly discernible growth defect, which is further amplified in cells concurrently lacking Rpl1a, and the synthesis of 60S subunits that can be devoid of Rpl1 (Fig. 10). Second, Acl1 and Bcl1, although not directly interacting with each other, can form a trimeric complex with Rpl1 in vitro (Fig. 9A and B), which, based on the prominent enrichment of Acl1 and Bcl1 in the reciprocal TurboID assays (Figs. 3B and 8C), must also occur in vivo, at least for some duration during Rpl1’s journey from the cytoplasm to its assembly site on nucleolar pre-60S particles. Third, the experimentally solved co-structure of Acl1’s ankyrin repeat domain in complex with the second Rpl1 domain as well as AlphaFold3 predictions of the trimeric complex revealed that the suggested mode of Rpl1 accommodation would efficiently shield almost all of its positively charged surfaces (Fig. 9D), which could prevent the aggregation of Rpl1 and/or facilitate its transport through the selective phase of the NPC.

Based on the here presented data and the current state of knowledge, we propose the following model for how Rpl1, at least in the yeast *S. cerevisiae*, might be chaperoned by Acl1 and Bcl1 on its voyage to its rRNA-binding site on the L1 stalk within nucleolar pre-60S particles (Fig. 13). Considering the prominent reciprocal enrichment of Acl1 and Rpl1 in their TurboID assays, the efficient formation of a soluble dimeric Acl1-Rpl1 complex upon co-expression of the two proteins in *E. coli*, and their simple mode of interaction (Figs. 3, 4B, and 6A), we believe that Acl1 binds very early on to newly synthesized Rpl1; however, most likely not in a co-translational manner (Supplementary Fig. S19). As nuclear accumulation of Acl1 and Bcl1 is only abrogated when both proteins simultaneously lack their C-terminal NLS region (Fig. 11D and E), one can infer that Acl1 and Bcl1 can get co-imported and, thus, that the trimeric Acl1-Rpl1-Bcl1 complex can already form in the cytoplasm. Notably, as indicated by the lack of nuclear accumulation of Rpl1b-yEGFP in Δ*acl1*/Δ*bcl1* cells when nuclear export of pre-60S subunits is impaired (Fig. 12), nuclear import of Rpl1 appears to fully depend on Acl1 and Bcl1. We have shown that the C-terminal extremities of both Acl1 and Bcl1 exhibit nuclear targeting activity (Fig. 11A and B). While the very C-terminal KRRK sequence of Acl1 is predicted to be recognized in a canonical manner by the major NLS-binding site of Kap60 and, thus, most likely enables classical import via the Kap60-Kap95 importin-α/β heterodimer [133,207] (Supplementary Fig. S18A), the NLS region of Bcl1 appears to be more complex as it not only contains a C-terminally located bPY-NLS that interacts with the importin-β Kap104 in Y2H assays [152,164] (Fig. 11B and C and Supplementary Fig. S18B and C), but likely also features a binding site for the importin-β Kap121 (Supplementary Fig. S18G), which is the only discernibly enriched importin in the proximity labelling assay with the Bcl1-TurboID bait (Fig. 8C and Supplementary Fig. S18F), and presumably also for at least one additional, yet-to-be-assigned importin. These observations suggest that nuclear import of the trimeric Acl1-Rpl1-Bcl1 complex could be independently mediated by at least three different importins; hence, further experiments are required to first corroborate an actual contribution of the above-mentioned importins and, second, to reveal the favoured import pathway of the trimeric complex. However, considering the minor phenotypic consequences of the individual absence of Acl1 or Bcl1 (Fig. 10A and B), their capacity to independently interact with Rpl1 in vitro (Figs. 4B and 7H), and the finding that Rpl1 variants deficient in Acl1 or Bcl1 binding can still engage in a Y2H interaction with Bcl1 or, respectively, Acl1 (Figs. 7G and 9G), it can be assumed that import of Rpl1 does not obligatorily require prior formation of the trimeric complex but could also occur in the context of the dimeric Acl1-Rpl1 and, presumably to a lesser degree, the dimeric Bcl1-Rpl1 complex. Moreover, as yeast cells lacking both Acl1 and Bcl1 are viable, whereas Rpl1 is an essential LSU r-protein that could be visualized in the cryo-EM structures both of early and late nucleolar pre-60S intermediates [98,116], one would expect that Rpl1 should additionally also be able to enter the nucleus on its own, either by passive diffusion or by its direct association with an importin. However, our cell biological finding that Rpl1b-yEGFP exhibits an exclusive cytoplasmic localization upon inhibition of nuclear pre-60S export in cells lacking Acl1 and Bcl1 suggests that nuclear import of Rpl1 is largely dependent on Acl1 and Bcl1 (Fig. 12), and, thus, rather raises the possibility that Rpl1, at least when Acl1 and Bcl1 are absent, can also get incorporated into cytoplasmic pre-60S intermediates. Under normal conditions, however, we assume that newly synthesized Rpl1 is efficiently captured by Acl1 and presumably also by Bcl1, but likely not in a co-translation manner (Supplementary Fig. S19), and, thus, would get rapidly imported into the nucleus, primarily in the context of the trimeric Acl1-Rpl1-Bcl1 and, to a lesser extent, the dimeric Acl1-Rpl1 complex, and preferentially incorporated into nucleolar pre-60S particles. Once in the nucleus, these Rpl1-containing cargo complexes are expected to be released from the respective transport-mediating importin by the binding of RanGTP, enabling the export and recycling of the importin. In case that the dimeric Acl1-Rpl1 complex was imported, the trimeric complex could potentially still be formed in the nucleus upon recruitment of Bcl1, which exhibits both a nucleolar and nucleoplasmic steady-state localization (Fig. 8G), from its free nuclear pool.

**Figure 13.**
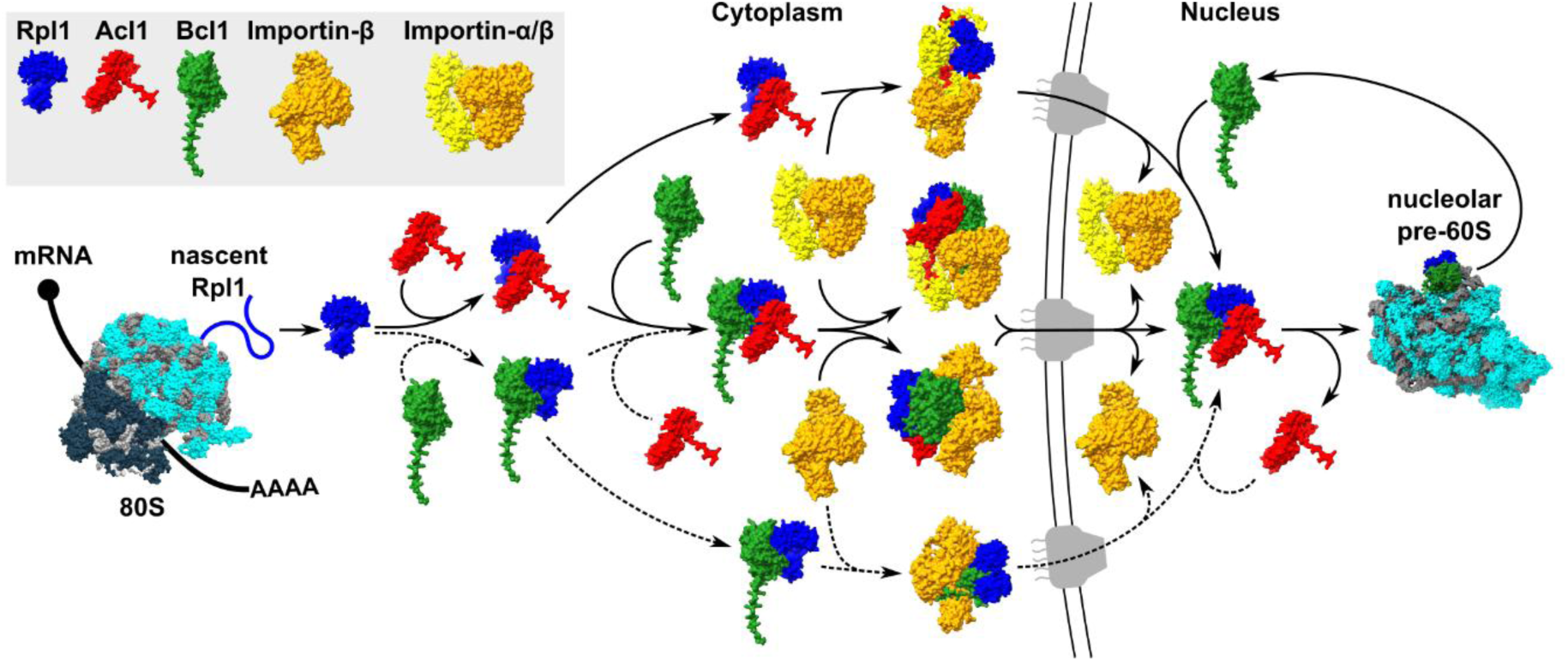
Simplified model showing how Acl1 and Bcl1 functionally cooperate to mediate the safe transfer of newly synthesized Rpl1 to its rRNA-binding site on the L1 stalk within nucleolar pre-60S particles. Upon its release from translating 80S ribosomes, newly synthesized Rpl1 is captured by Acl1 and/or Bcl1 and imported either as a dimeric Acl1-Rpl1 and trimeric Acl1-Rpl1-Bcl1 complex (main pathways) or a dimeric Bcl1-Rpl1 complex (minor pathway). Nuclear import of these complexes is mediated by importins, most likely the importin-α/β heterodimer Kap60-Kap95 (Acl1-Rpl1 and Acl1-Rpl1-Bcl1) or the importin-β Kap104 (Bcl1-Rpl1 and Acl1-Rpl1-Bcl1), upon recognition of the respective NLSs that are located at the C-terminal extremity of both Acl1 and Bcl1. Once in the nucleus, these cargo complexes are dissociated from the importins by the binding of RanGTP. As Acl1’s ankyrin repeat domain protects one of the two rRNA-binding surfaces of Rpl1, Acl1 will get efficiently dissociated by the full integration of Rpl1 into nucleolar pre-60S particles, whereas Bcl1, as suggested by the identification of Rpl1-proximal AFs in its proxiOME, may remain transiently associated until a certain maturation stage with nucleolar pre-60S particles. For simplicity, the anticipated return of Acl1 and Bcl1 to the cytoplasm (for another round of Rpl1 binding, import, and delivery), the RanGTP-mediated cargo release in the nucleus, the export of exportin-bound importin-α and RanGTP-bound importin-βs, and the cytoplasmic recycling of importins (dissociation of RanGTP upon its conversion to RanGDP) for another round of cargo binding are not depicted. For more details, see Discussion.

Based on the observed and predicted interactions of Rpl1 with the L1 stalk rRNA and within the trimeric complex as well as the revealed in vivo neighbourhoods of Acl1 and Bcl1, we envisage the following scenario for the transfer of Rpl1 from the trimeric complex onto its pre-ribosomal binding site. As the robust interaction between the ankyrin repeat domain of Acl1 and the short segment within the second structural domain of Rpl1 occludes the rRNA-binding surface of the D2 domain [98] (Figs. 5D and 6A and Supplementary Figs. S13B and S16A), the initial interaction of Rpl1 with the L1 stalk rRNA is most likely mediated by the composite rRNA-binding surface of the D1 domain (Supplementary Fig. S13B). This surface might, however, also be partially masked by predicted contacts with the predicted three-helix fold within Acl1’s C-terminal part (Supplementary Fig. S16A and B). But, as the three-helix fold is only very weakly interacting with Rpl1 in Y2H assays (Supplementary Fig. S16C), it can be assumed that the D1 domain will preferentially bind to its cognate rRNA-binding site. Once docked via the D1 domain on helix H77 of the L1 stalk, the presumed higher affinity of the second rRNA-binding site, consisting of rRNA residues that are part of the bulge connecting 25S rRNA helices H78 and H77 [98] (Supplementary Fig. S13B), is expected to enable the rapid release of Acl1 from its binding site on the D2 domain and the full binding of Rpl1. In line with a highly transient association of Acl1 with pre-60S particles during the process of Rpl1 incorporation, no Rpl1-surrounding pre-60S AFs were enriched in the Acl1 TurboID assays (Fig. 3B). In contrast, Bcl1 appears to remain associated with Rpl1 upon Acl1 release, as indicated by the moderate enrichment of several nucleolar pre-60S AFs, which notably all cluster around Bcl1 when the predicted Bcl1-Rpl1 complex is superposed onto Rpl1 in the cryo-EM structure of the late nucleolar state E2 pre-60S intermediate [98] (Fig. 8D), in the proximity labelling assay with the Bcl1-TurboID bait (Fig. 8C). According to this superposition, an interaction of Bcl1 with rRNA-bound Rpl1 is possible but would most likely be precluded by a clash of β-propeller blades 1 to 3 of Bcl1 with Nop2 (Fig. 8E), which is associated with the concave surface of the L1 stalk rRNA. This raises the possibility that Bcl1 could get displaced from Rpl1 by Nop2 and would imply that incorporation of Rpl1, when transferred from the trimeric complex, very likely occurs prior to the full integration of Nop2 into nucleolar pre-60S particles. How Acl1 and Bcl1 get back to the cytoplasm for another round of recognition and import of newly synthesized Rpl1 remains to be determined, but their relatively small sizes (calculated molecular mass of 24.67 kDa and 43.83 kDa, respectively) would be compatible with a passive diffusion mechanism [205]. Clearly, future experiments will be required to fully appreciate the exact functional roles of Acl1 and Bcl1 as binding partners of Rpl1. Moreover, it will be of interest to learn how the protection, transport, and incorporation of uL1 is achieved in more advanced eukaryotic organisms, which notably appear to lack an Acl1-like binding partner of uL1. The identification of WDR89, the putative human orthologue of Bcl1, in the proximity labelling assay with human uL1 in HeLa cells (Fig. 7A), as well as the remarkable similarity of the predicted structure models of the human WDR89-RPL10A and the yeast Bcl1-Rpl1 complex (Fig. 7B and C), indicate that Bcl1-like proteins may play a prominent, potentially conserved role in this process in all eukaryotic organisms.

Surprisingly, Acl1 was the only novel selective r-protein binding partner that could be readily uncovered by our TurboID-based proximity labelling screen, raising the possibility that there might be no further DCs of LSU r-proteins in the yeast *S. cerevisiae* than the already discovered ones. Considering, however, that only six of the eight known DCs, as well as Acl1, were prominently enriched in the proximity labelling assays of their respective N– and/or C-terminally TurboID-tagged r-protein client (Fig. 3A and Supplementary Fig. S6), while the reciprocal TurboID assays with the DC baits, except in the case of Bcl1, effortlessly identified each respective r-protein client (Fig. 3B and Supplementary Fig. S8), it is possible that our proximity labelling screen may not have revealed all previously undiscovered DCs of LSU r-proteins. Moreover, we deem it likely that some selective r-protein interactors, as observed for Bcl1 in the assays with the TurboID-Rpl1b bait (Fig. 3A), could still be hidden among the numerous, less prominently enriched proteins, indicating that the comprehensive identification of further candidate DCs would require a more detailed analysis of the data set. Alternatively, more general binding partners of free basic proteins such as the histone chaperone Nap1, which has previously been shown to not only interact with the SSU r-protein Rps6 (eS6) but also, at least in Y2H assays, with the LSU r-proteins Rpl39 (eL39) and Rpl42 [44], could take over a protective function for several r-proteins. In line with this conjecture, Nap1 was discernibly, albeit to different extents, enriched in the TurboID assays of Rpl39 and Rpl42 as well as in the ones of additional LSU r-proteins, including Rpl8, Rpl15 (eL15), Rpl24, and Rpl37 (eL37) (Supplementary Fig. S1). Not unexpectedly, importins emerged as another class of transient interaction partners of LSU r-proteins in our TurboID-based proximity labelling screen. Here, we provide first evidence, by employing Y2H assays, for a direct interaction between four importin/r-protein pairs that were highlighted by the TurboID assays (Fig. 2). While Rpl6, Rpl30, and Rpl38 likely correspond to bona fide import cargo of Kap114, Kap119, and, respectively, Kap121, the finding that Rpl40, which gets incorporated during the cytoplasmic stage of pre-60S maturation [100,105,118], only interacts with full-length Kap114 but not with variants lacking partially or completely the N-terminal RanGTP-binding site in the Y2H assays suggests that Kap114 could mediate the export of Rpl40. A bidirectional transport activity has been previously assigned to Kap122 [154,163]; thus, Kap114, if indeed exporting Rpl40, would be the second *S. cerevisiae* biportin [152]. Notably, these four r-proteins are predicted to be almost entirely accommodated by the respective importin in the AlphaFold3 models of the binary complexes (Fig. 2). Compared to the prevailing recognition of short linear NLS sequences for nuclear targeting, the predicted entrapment of these r-proteins by the inner concave surface of the importins would efficiently shield their positively charged surfaces. Thus, our study reinforces the notion that importins, as already postulated by the Görlich laboratory more than 20 years ago [9], combine the protection of basic domains with their nucleocytoplasmic transport.

Taken together, this study has highlighted the suitability of the TurboID-based proximity labelling approach for the identification of transient interaction partners of r-proteins. We believe that this data set will not only fuel our exciting journey towards a comprehensive overview of all transient interactors of unassembled r-proteins but may also serve as an inspiring resource for illuminating interactions on pre-ribosomal particles that have escaped structural analyses and for exploring the unknOME side of the LSU r-protein proxiOMEs.

## ACKNOWLEDGEMENTS

We gratefully acknowledge the Bioimage Core Facility of the University of Fribourg as well as Michael Stumpe of the Proteomics Unit of the Metabolomics and Proteomics Platform (MAPP) of the Department of Biology of the University of Fribourg for their support and assistance. We thank the European Synchrotron Radiation Facility (ESRF) for excellent support during data collection and Jesús de la Cruz for providing the anti-Rpl1 antibody.

## AUTHOR CONTRIBUTIONS

S.F.: Conceptualization, Data curation, Formal analysis, Investigation, Methodology, Project administration, Software, Supervision, Validation, Visualization, Writing – original draft, Writing – review and editing; B.P.: Conceptualization, Formal analysis, Investigation, Methodology, Software, Visualization, Writing – review and editing; F.B.: Data curation, Formal analysis, Investigation; D.S.S.: Data curation, Formal analysis, Investigation; A.M.-G.: Investigation; S.K.: Data curation, Formal analysis; J.D.: Data curation, Funding acquisition, Resources, Supervision, Writing – review and editing; G.B.: Data curation, Funding acquisition, Resources, Supervision; D.K.: Conceptualization, Data Curation, Formal analysis, Funding acquisition, Investigation, Methodology, Project administration, Resources, Supervision, Validation, Visualization, Writing – original draft, Writing – review and editing.

## DATA AVAILABILITY

The mass spectrometry proteomics data have been deposited to the ProteomeXchange Consortium via the PRIDE [219] partner repository (https://www.ebi.ac.uk/pride/) with the dataset identifiers PXD070550 and PXD070586.

Atomic coordinates and structure factors for the crystal structure of the Rpl1(63-158)-Acl1.N127 complex have been deposited to the Protein Data Bank under accession code 9T3L.

