## Supplementary data for "Exploration of the proxiOME of large subunit ribosomal proteins reveals Acl1 and Bcl1 as cooperating dedicated chaperones of Rpl1"

### SUPPLEMENTARY FIGURES

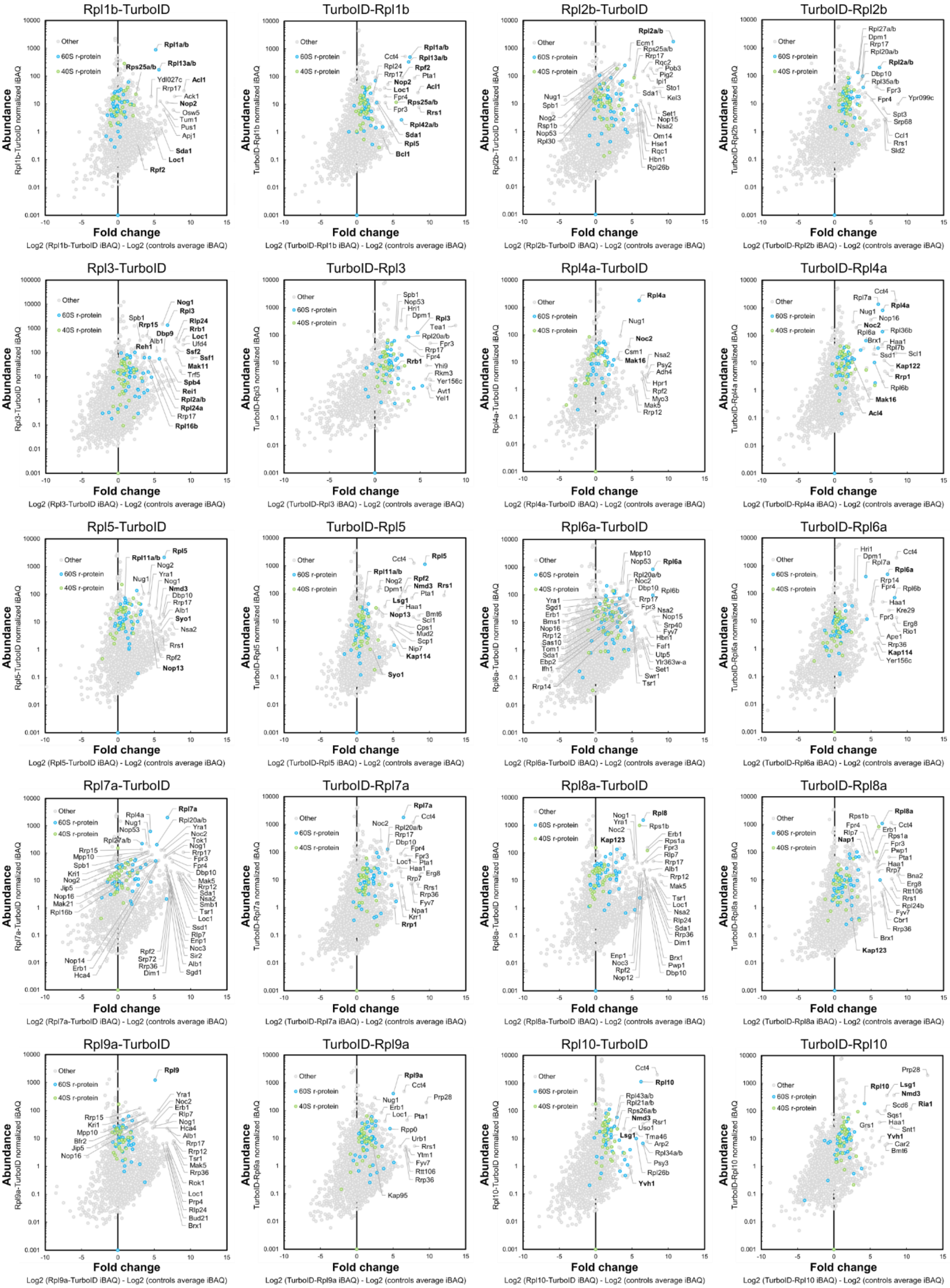

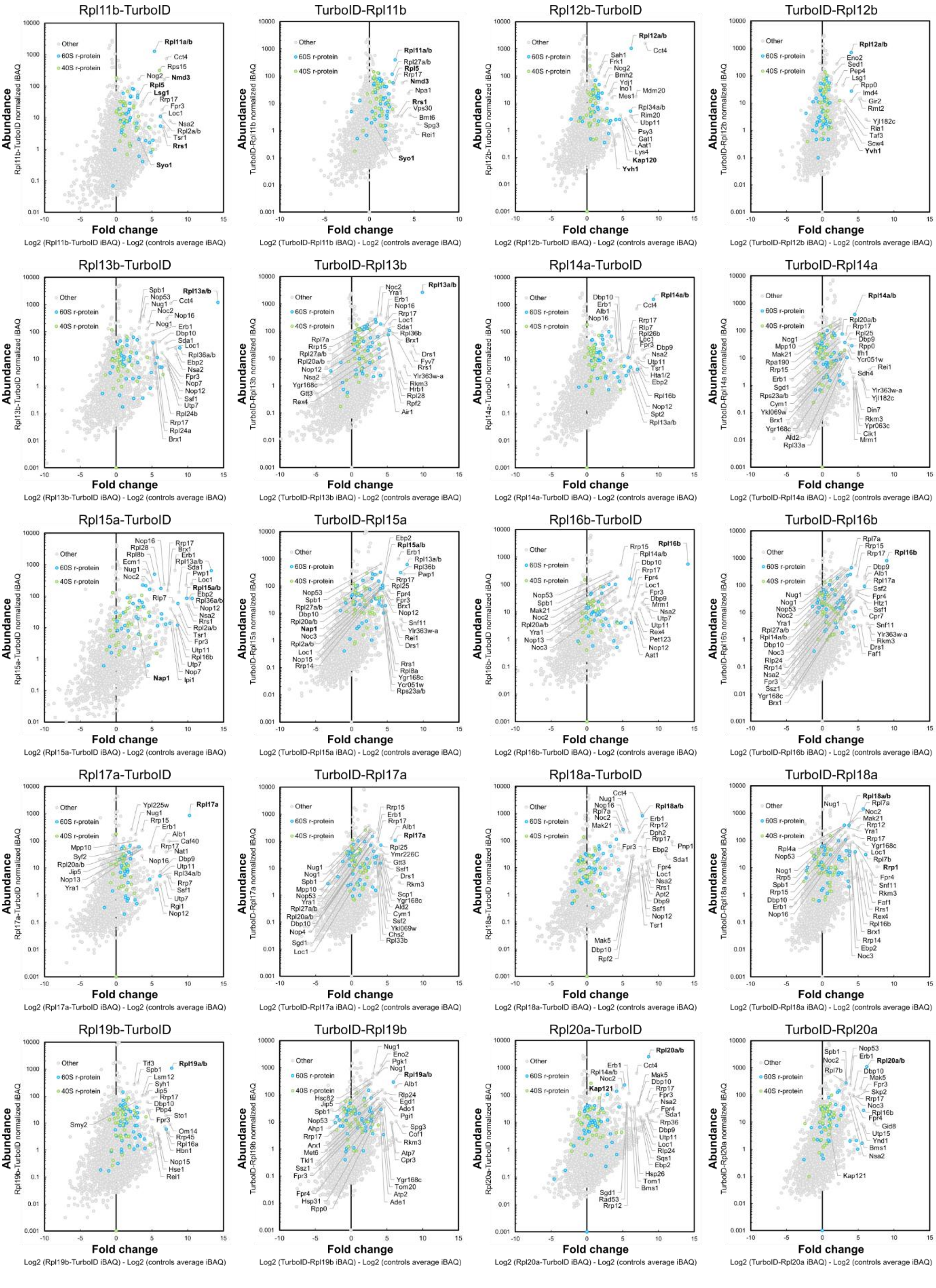
Rpl11b-TurboID
TurboID-Rpl11b
Rpl12b-TurboID
TurboID-Rpl12b
Rpl13b-TurboID
TurboID-Rpl13b
Rpl14a-TurboID
TurboID-Rpl14a
Rpl15a-TurboID
TurboID-Rpl15a
Rpl16b-TurboID
TurboID-Rpl16b
Rpl17a-TurboID
TurboID-Rpl17a
Rpl18a-TurboID
TurboID-Rpl18a
Rpl19b-TurboID
TurboID-Rpl19b
Rpl20a-TurboID
TurboID-Rpl20a
Other
60S r-protein
40S r-protein
Abundance
Fold change

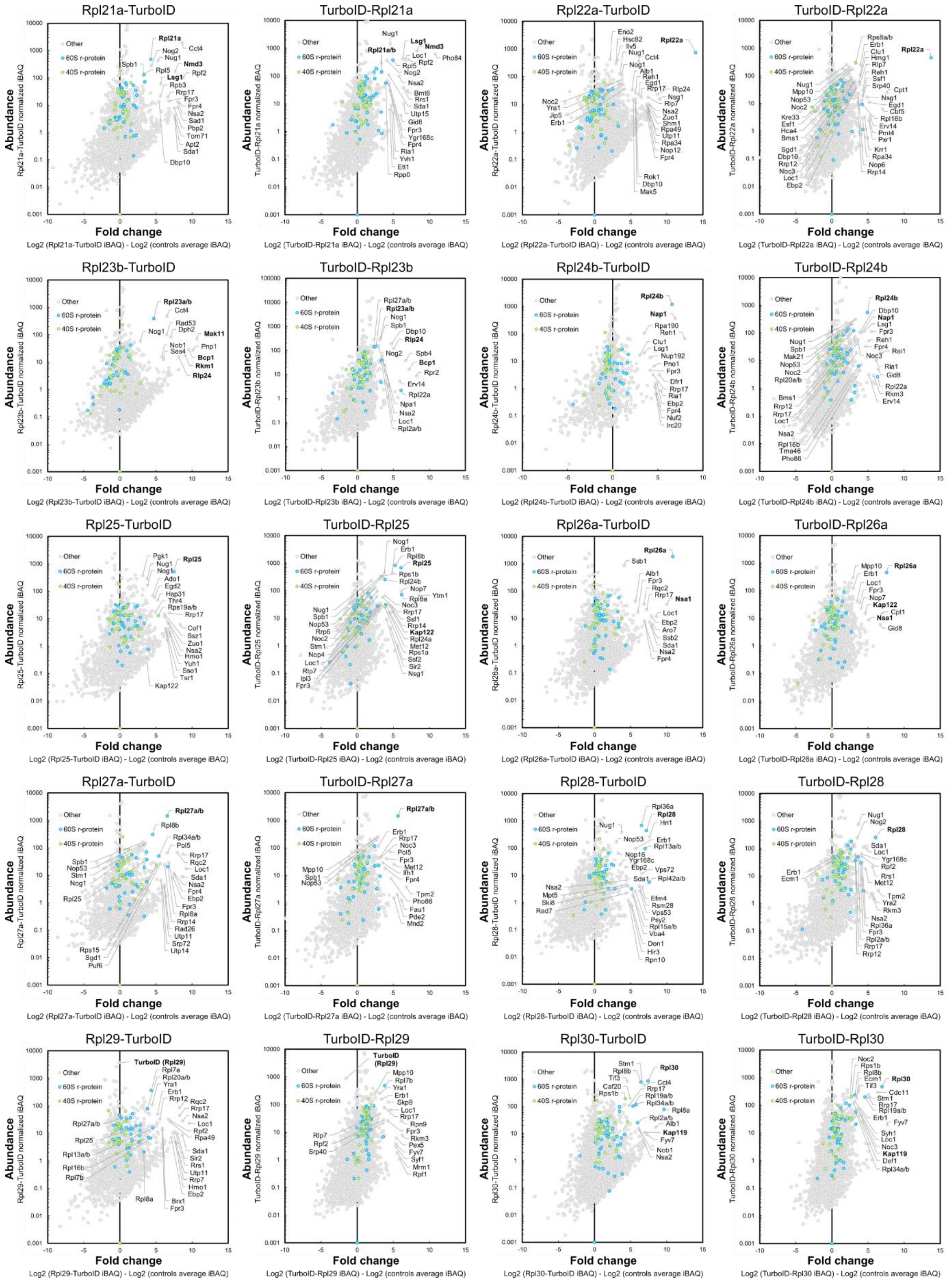

Rpl21a-TurboID
TurboID-Rpl21a
Rpl22a-TurboID
TurboID-Rpl22a
Rpl23b-TurboID
TurboID-Rpl23b
Rpl24b-TurboID
TurboID-Rpl24b
Rpl25-TurboID
TurboID-Rpl25
Rpl26a-TurboID
TurboID-Rpl26a
Rpl27a-TurboID
TurboID-Rpl27a
Rpl28-TurboID
TurboID-Rpl28
Rpl29-TurboID
TurboID-Rpl29
Rpl30-TurboID
TurboID-Rpl30
Other
60S r-protein
40S r-protein
Abundance
Fold change
Log2 (Rpl21a-TurboID iBAQ) - Log2 (controls average iBAQ)
Log2 (TurboID-Rpl21a iBAQ) - Log2 (controls average iBAQ)
Log2 (Rpl22a-TurboID iBAQ) - Log2 (controls average iBAQ)
Log2 (TurboID-Rpl22a iBAQ) - Log2 (controls average iBAQ)
Log2 (Rpl23b-TurboID iBAQ) - Log2 (controls average iBAQ)
Log2 (TurboID-Rpl23b iBAQ) - Log2 (controls average iBAQ)
Log2 (Rpl24b-TurboID iBAQ) - Log2 (controls average iBAQ)
Log2 (TurboID-Rpl24b iBAQ) - Log2 (controls average iBAQ)
Log2 (Rpl25-TurboID iBAQ) - Log2 (controls average iBAQ)
Log2 (TurboID-Rpl25 iBAQ) - Log2 (controls average iBAQ)
Log2 (Rpl26a-TurboID iBAQ) - Log2 (controls average iBAQ)
Log2 (TurboID-Rpl26a iBAQ) - Log2 (controls average iBAQ)
Log2 (Rpl27a-TurboID iBAQ) - Log2 (controls average iBAQ)
Log2 (TurboID-Rpl27a iBAQ) - Log2 (controls average iBAQ)
Log2 (Rpl28-TurboID iBAQ) - Log2 (controls average iBAQ)
Log2 (TurboID-Rpl28 iBAQ) - Log2 (controls average iBAQ)
Log2 (Rpl29-TurboID iBAQ) - Log2 (controls average iBAQ)
Log2 (TurboID-Rpl29 iBAQ) - Log2 (controls average iBAQ)
Log2 (Rpl30-TurboID iBAQ) - Log2 (controls average iBAQ)
Log2 (TurboID-Rpl30 iBAQ) - Log2 (controls average iBAQ)

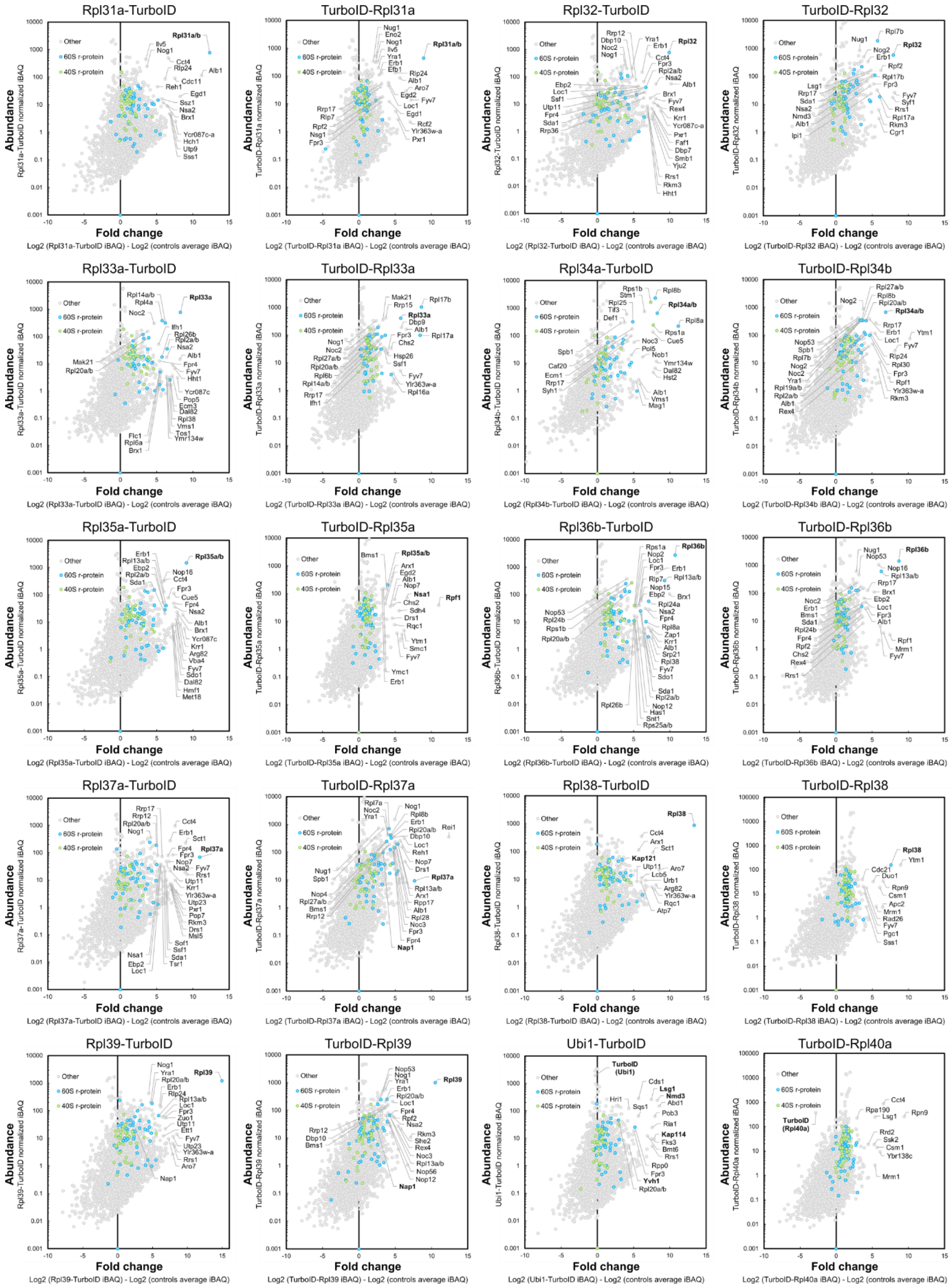

Rpl31a-TurboID
TurboID-Rpl31a
Rpl32-TurboID
TurboID-Rpl32
Rpl33a-TurboID
TurboID-Rpl33a
Rpl34a-TurboID
TurboID-Rpl34b
Rpl35a-TurboID
TurboID-Rpl35a
Rpl36b-TurboID
TurboID-Rpl36b
Rpl37a-TurboID
TurboID-Rpl37a
Rpl38-TurboID
TurboID-Rpl38
Rpl39-TurboID
TurboID-Rpl39
Ubi1-TurboID
TurboID-Rpl40a
Other
60S r-protein
40S r-protein
Abundance
Fold change
Log2 (Rpl31a-TurboID iBAQ) - Log2 (controls average iBAQ)
Log2 (TurboID-Rpl31a iBAQ) - Log2 (controls average iBAQ)
Log2 (Rpl32-TurboID iBAQ) - Log2 (controls average iBAQ)
Log2 (TurboID-Rpl32 iBAQ) - Log2 (controls average iBAQ)
Log2 (Rpl33a-TurboID iBAQ) - Log2 (controls average iBAQ)
Log2 (TurboID-Rpl33a iBAQ) - Log2 (controls average iBAQ)
Log2 (Rpl34b-TurboID iBAQ) - Log2 (controls average iBAQ)
Log2 (TurboID-Rpl34b iBAQ) - Log2 (controls average iBAQ)
Log2 (Rpl35a-TurboID iBAQ) - Log2 (controls average iBAQ)
Log2 (TurboID-Rpl35a iBAQ) - Log2 (controls average iBAQ)
Log2 (Rpl36b-TurboID iBAQ) - Log2 (controls average iBAQ)
Log2 (TurboID-Rpl36b iBAQ) - Log2 (controls average iBAQ)
Log2 (Rpl37a-TurboID iBAQ) - Log2 (controls average iBAQ)
Log2 (TurboID-Rpl37a iBAQ) - Log2 (controls average iBAQ)
Log2 (Rpl38-TurboID iBAQ) - Log2 (controls average iBAQ)
Log2 (TurboID-Rpl38 iBAQ) - Log2 (controls average iBAQ)
Log2 (Rpl39-TurboID iBAQ) - Log2 (controls average iBAQ)
Log2 (TurboID-Rpl39 iBAQ) - Log2 (controls average iBAQ)
Log2 (Ubi1-TurboID iBAQ) - Log2 (controls average iBAQ)
Log2 (TurboID-Rpl40a iBAQ) - Log2 (controls average iBAQ)

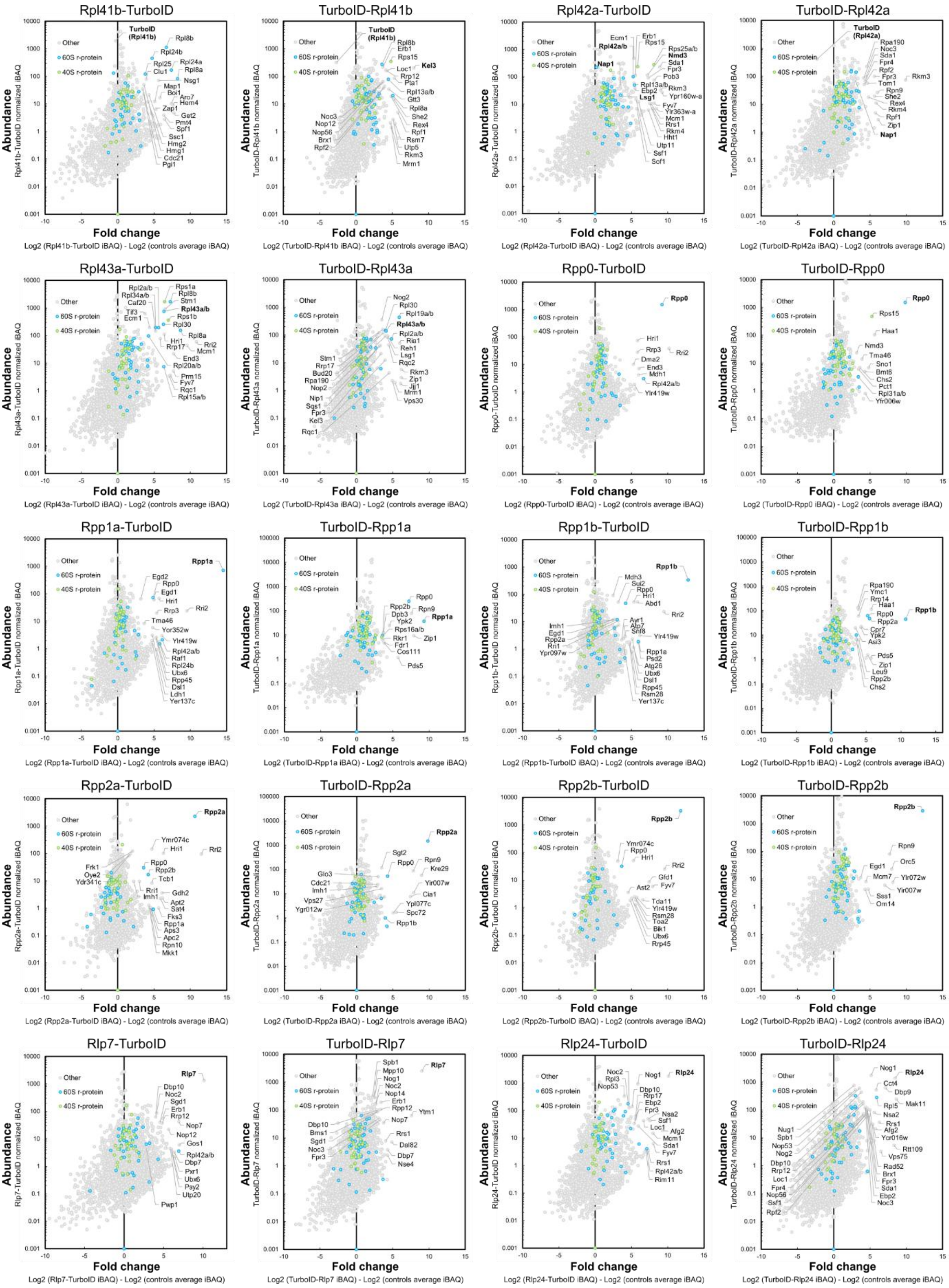
Rpl41b-TurboID
TurboID-Rpl41b
Rpl42a-TurboID
TurboID-Rpl42a
Rpl43a-TurboID
TurboID-Rpl43a
Rpp0-TurboID
TurboID-Rpp0
Rpp1a-TurboID
TurboID-Rpp1a
Rpp1b-TurboID
TurboID-Rpp1b
Rpp2a-TurboID
TurboID-Rpp2a
Rpp2b-TurboID
TurboID-Rpp2b
Rlp7-TurboID
TurboID-Rlp7
Rlp24-TurboID
TurboID-Rlp24
Other
60S r-protein
40S r-protein
Abundance
Fold change

**Supplementary Figure S1. The proxiOME of LSU r-proteins.** Graphical representation of the TurboID results obtained with all N- and C-terminally TurboID-tagged LSU r-proteins as well as with the ribosomal-like AFs Rlp7 and Rlp24. The employed bait protein is indicated above each graph. The bait proteins, enriched DCs and importins, and selected enriched r-proteins and AFs are written in bold. Note that in the case of the N- and C-terminally TurboID-tagged Rpl29, Rpl40a/Ubi1, and Rpl41b bait proteins and of TurboID-Rpl42a only the TurboID moiety but not the fused r-protein could be detected in the analysis of non-biotinylated tryptic peptides, which were used for the identification and quantification of proteins. In these cases, the abundance and relative enrichment of the fused TurboID moiety is indicated in the graphs.

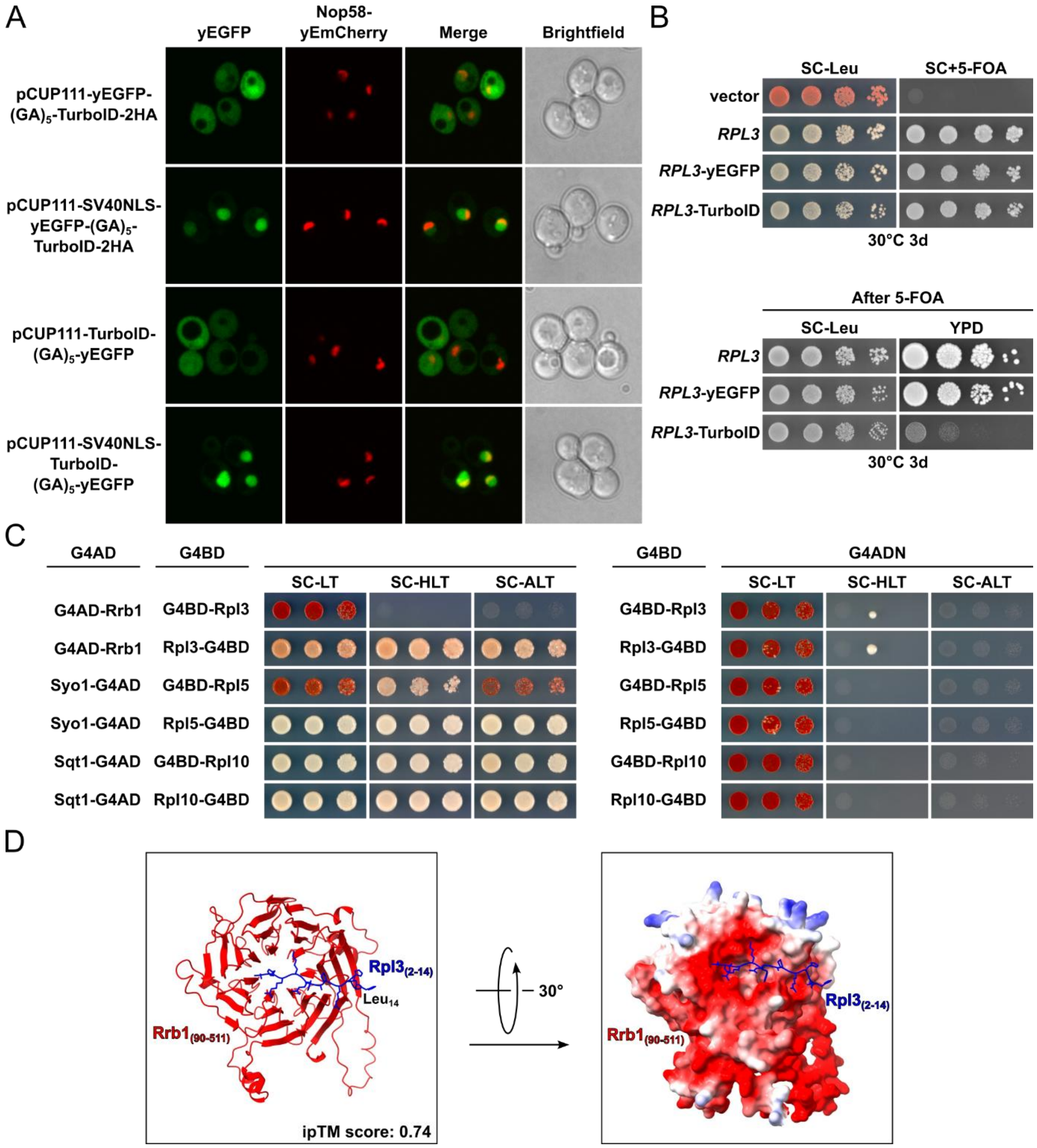


**Supplementary Figure S2.** **(A)** Localization of the four TurboID-tagged GFP control proteins, expressed for 1 h in the presence of 0.1 mM $CuSO_4$ under the control of the *CUP1* promoter from plasmid, was assessed by fluorescence microscopy in cells expressing the nucleolar marker protein Nop58-yEmCherry from the genomic locus that were grown in SC-Leu medium at 30ºC. **(B)** Evaluation of the functionality of C-terminally TurboID-tagged Rpl3. Empty vector (YCplac111) and plasmids harbouring *RPL3*, *RPL3*-yEGFP, or *RPL3*-TurboID, expressed under the transcriptional control of the *RPL3* promoter, were transformed into an *RPL3* shuffle strain. Transformants were restreaked on SC-Leu plates and cells were then spotted in 10-fold serial dilution steps onto SC-Leu plates and SC plates containing 5-fluoroorotic acid (5-FOA), which were incubated for 3 days at 30°C. After plasmid shuffling, cells were restreaked on SC-Leu plates and then spotted in 10-fold serial dilution steps onto SC-Leu and YPD plates. **(C)** Y2H interaction assays between the r-proteins Rpl3, Rpl5, and Rpl10 (tagged at their N- or C-termini with the G4BD) and their respective G4AD-tagged DC Rrb1, Syo1, and Sqt1. **(D)** AlphaFold3 model of the Rrb1-Rpl3 complex. The prediction was done with full-length Rrb1 and Rpl3 lacking its N-terminal methionine as input, but, for clarity, only residues 90-511 of Rrb1 (red) and residues 2-14 of Rpl3 (blue) are shown in the cartoon representation (left) and the representation depicting the electrostatic surface potential of Rrb1 (right).

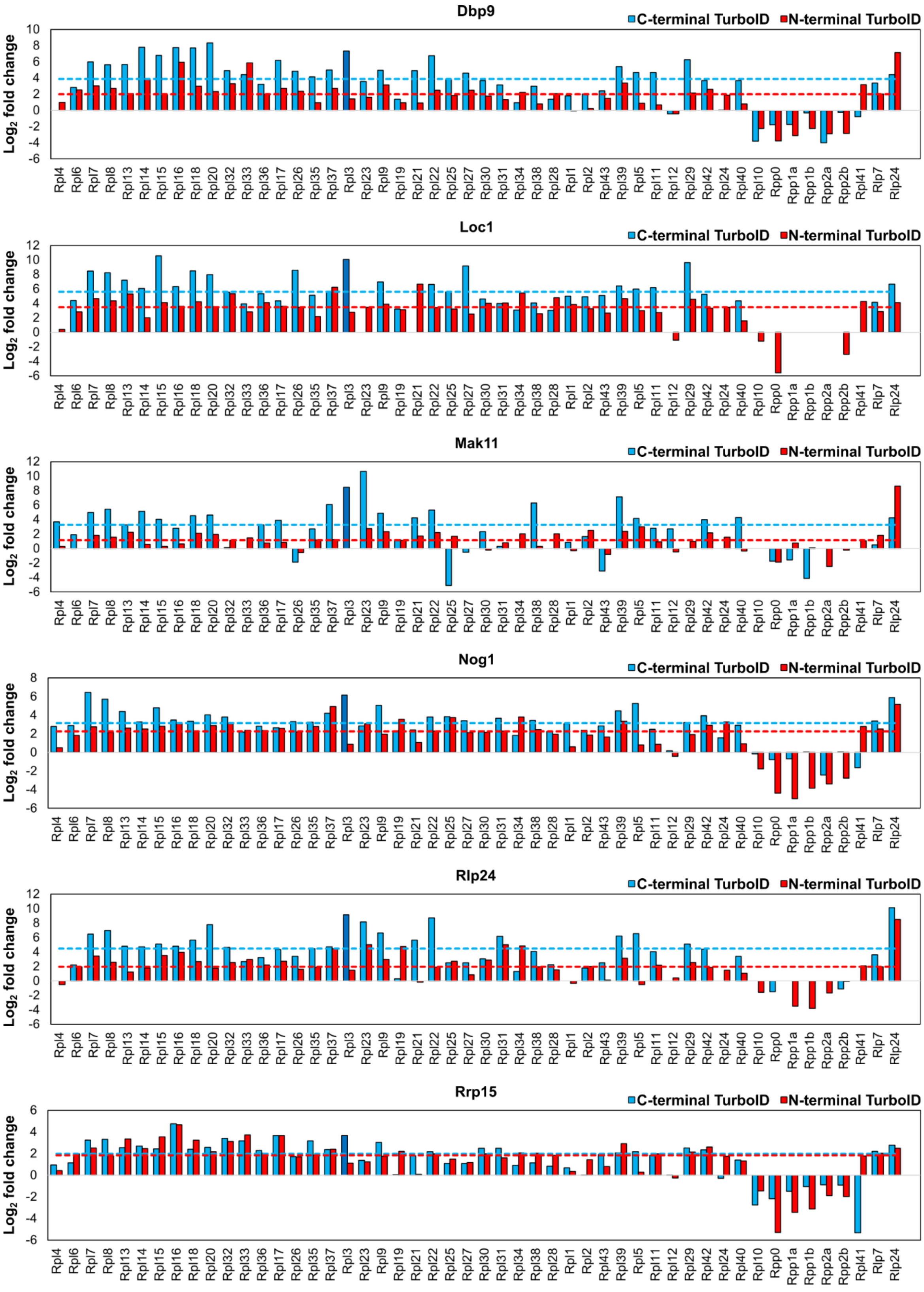
Dbp9
C-terminal TurboID
N-terminal TurboID
Log2 fold change
Rpl4 Rpl6 Rpl7 Rpl8 Rpl13 Rpl14 Rpl15 Rpl16 Rpl18 Rpl20 Rpl32 Rpl33 Rpl36 Rpl17 Rpl26 Rpl35 Rpl37 Rpl3 Rpl23 Rpl9 Rpl19 Rpl21 Rpl22 Rpl25 Rpl27 Rpl30 Rpl31 Rpl34 Rpl38 Rpl28 Rpl1 Rpl2 Rpl43 Rpl39 Rpl5 Rpl11 Rpl12 Rpl29 Rpl42 Rpl24 Rpl40 Rpl10 Rpp0 Rpp1a Rpp1b Rpp2a Rpp2b Rpl41 Rlp7 Rlp24
Loc1
Mak11
Nog1
Rlp24
Rrp15

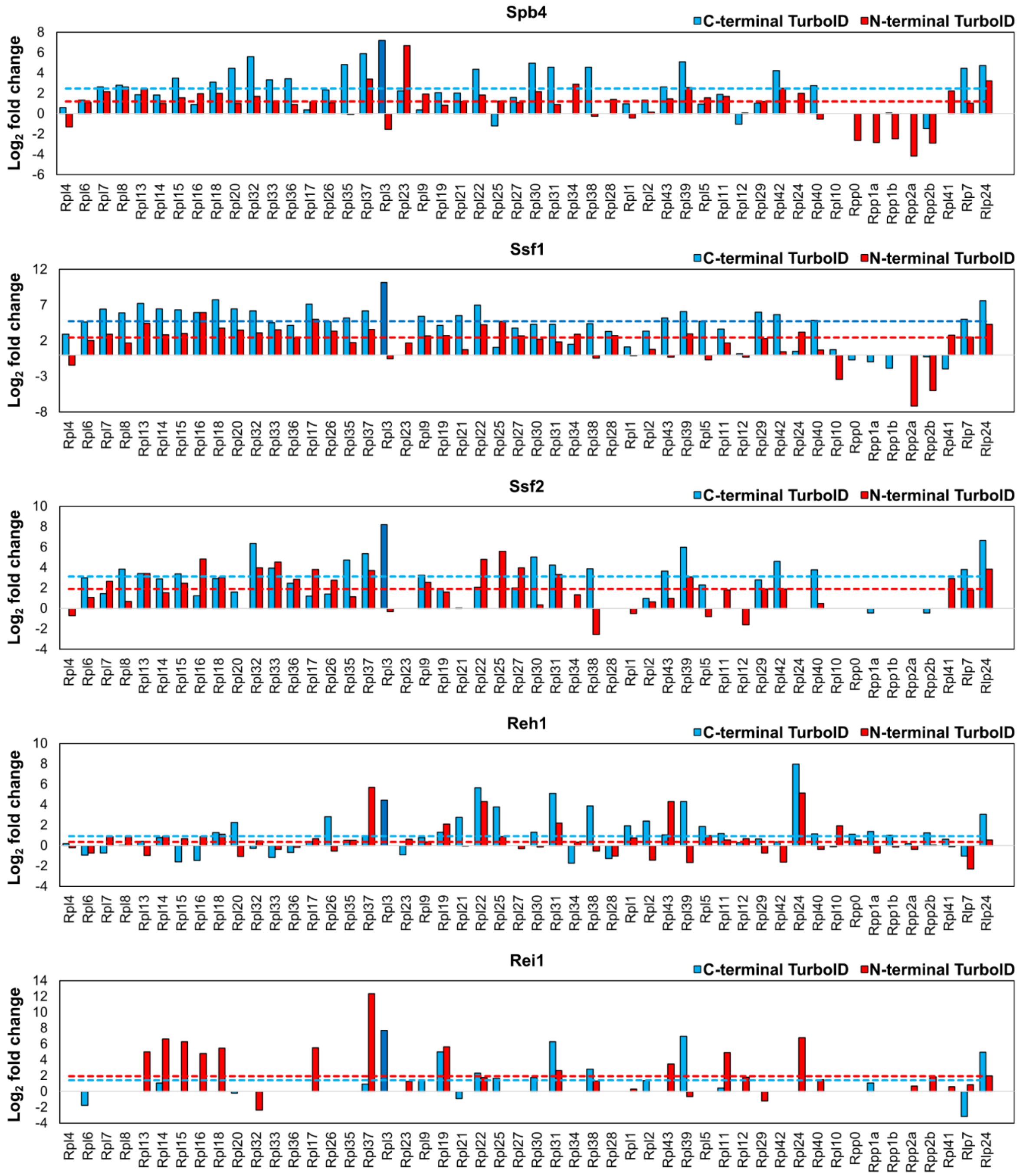


**Supplementary Figure S3.** Bar graphs showing the log2 fold enrichment of AFs, which were enriched in the TurboID assays with the Rpl3-TurboID bait (see Fig. 1B), in the TurboID assays of all N- (red bars) and C-terminally (blue bars) TurboID-tagged LSU r-proteins as well as the ones of Rlp7 and Rlp24. In the case of the Rpl3-TurboID bait (dark blue bars) and the TurboID-Rpl1 bait, their log2 fold enrichment corresponds to the average of the eight or, respectively, six experimental replicates. Dashed lines indicate the median log2 fold enrichment of an AF across all shown TurboID assays. The order of the LSU r-proteins (from left to right) is according to their first visualization in structures of early nucleolar to cytoplasmic pre-60S particles.

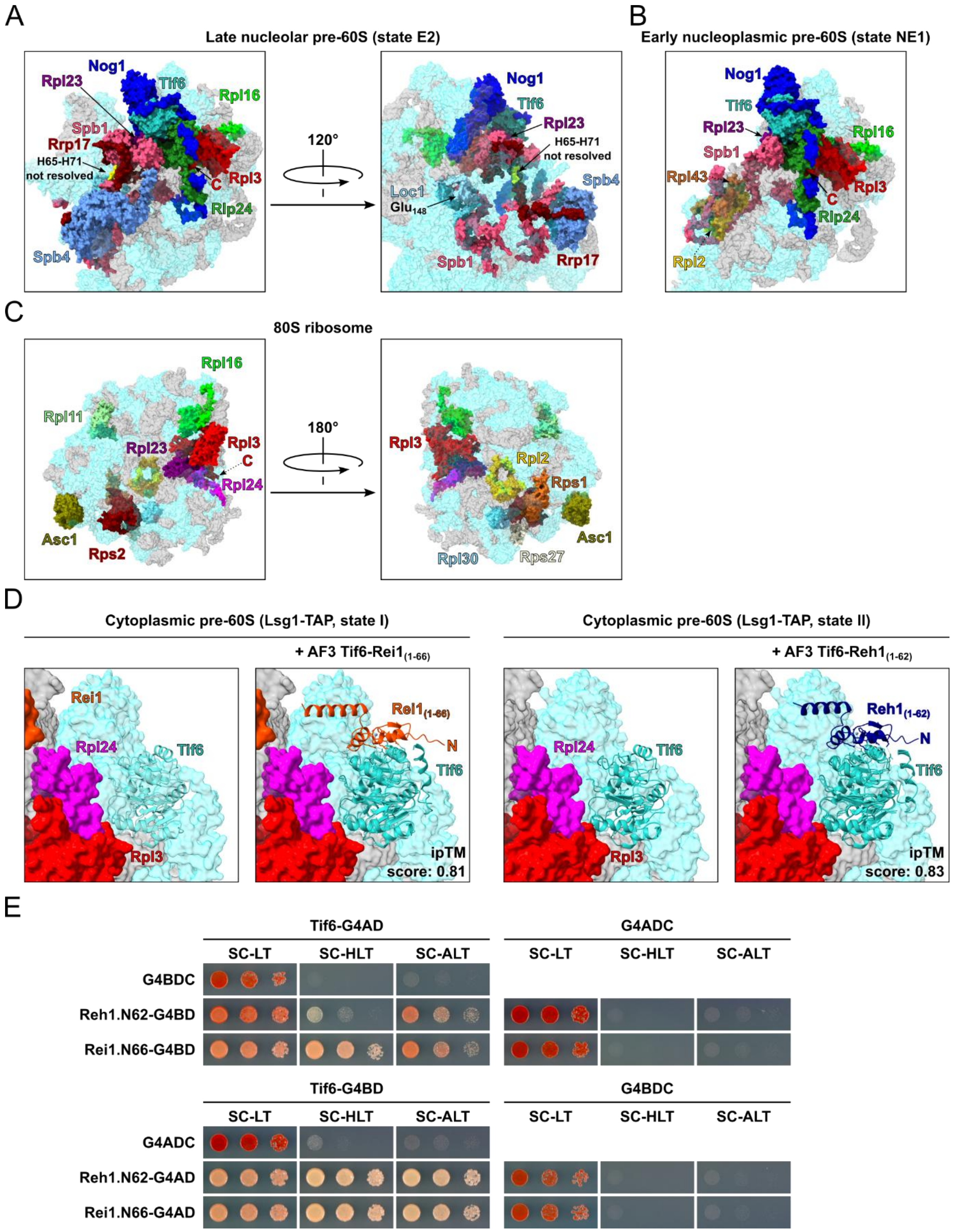


**Supplementary Figure S4. (A-C)** Location of Rpl3 and its neighbouring proteins on: **(A)** the late nucleolar state E2 pre-60S particle (PDB: 7NAC [1]), **(B)** the early nucleoplasmic state NE1 pre-60S particle (PDB: 7U0H [1]), and **(C)** the 80S ribosome (PDB 4V88 [2]). The indicated proteins have been highlighted in different colours; other r-proteins and AFs are coloured in light cyan and (pre-)rRNAs in light grey. **(D)** Superposition of the AlphaFold3 model of the Tif6-Rei1.N66 (left) and Tif6-Reh1.N62 (right) complex onto Tif6 (shown in cartoon representation) within the cytoplasmic Lsg1-TAP state I (left; PDB: 6RZZ) or Lsg1-TAP state II (right; PDB: 6S05) pre-60S particles [3], in which either parts of Rei1 or Reh1 can be visualized. Rei1 is coloured in orange and Reh1 in dark blue. **(E)** Y2H interaction assays between Tif6 and Reh1.N62 (residues 1-62) or Rei1.N66 (residues 1-66). Note that multicopy plasmids were used for these Y2H assays.

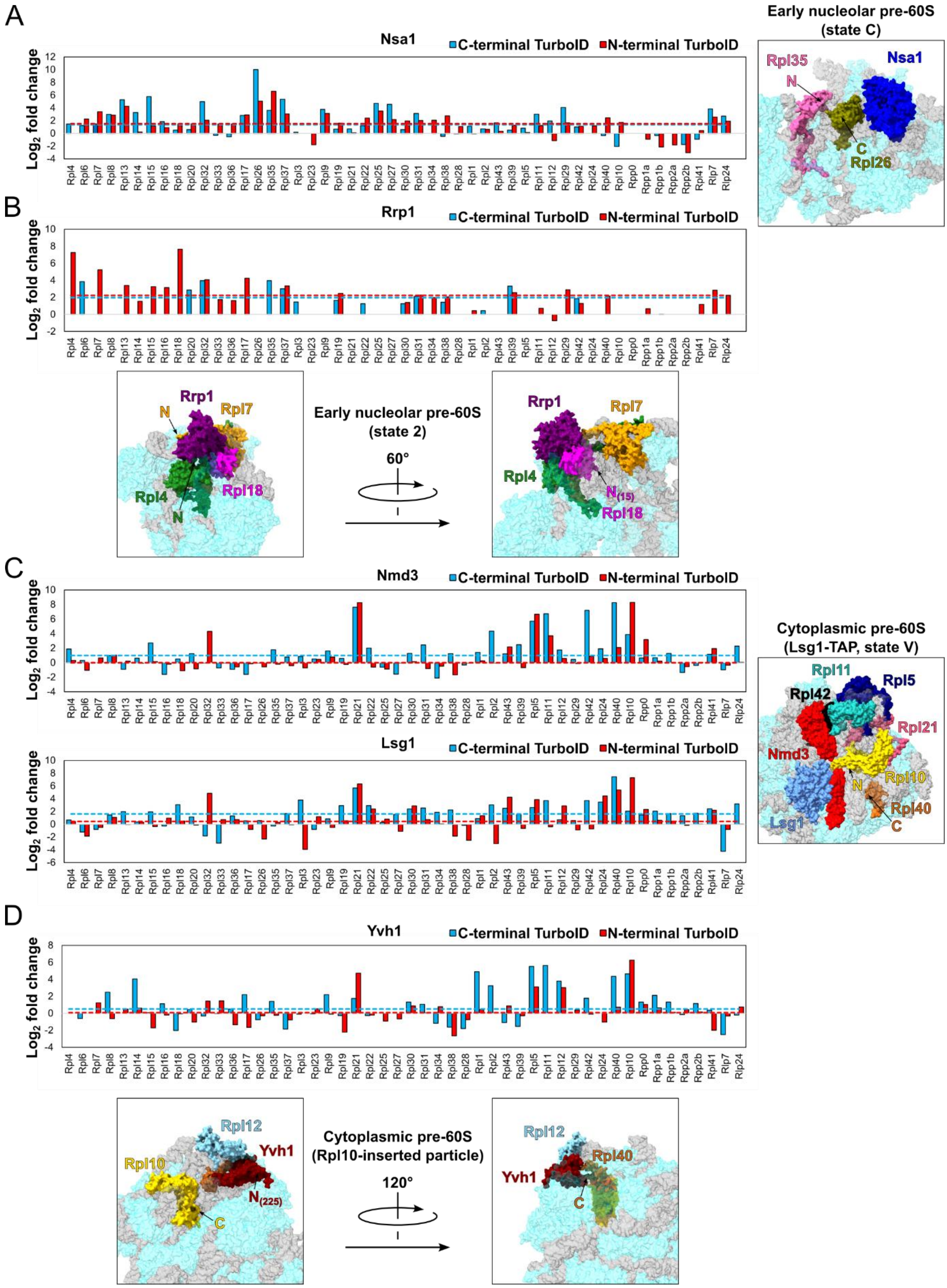


**Supplementary Figure S5. (A-D)** Bar graphs showing the log2 fold enrichment of Nsa1 **(A)**, Rrp1 **(B)**, Nmd3 and Lsg1 **(C)**, and Yvh1 **(D)** in the TurboID assays of all N- (red bars) and C-terminally (blue bars) TurboID-tagged LSU r-proteins as well as the ones of Rlp7 and Rlp24. In the case of the Rpl3-TurboID bait (dark blue bars) and the TurboID-Rpl1 bait, their log2 fold enrichment corresponds to the average of the eight or, respectively, six experimental replicates. Dashed lines indicate the median log2 fold enrichment of an AF across all shown TurboID assays. The order of the LSU r-proteins (from left to right) is according to their first visualization in structures of early nucleolar to cytoplasmic pre-60S particles. The right or lower panels show the location and neighbouring proteins of: **(A)** Nsa1 on the early nucleolar state C pre-60S particle (PDB: 6EM1 [4]), **(B)** Rrp1 on the early nucleolar state 2 pre-60S particle (PDB: 6C0F [5]), **(C)** Nmd3 and Lsg1 on a cytoplasmic pre-60S intermediate (Lsg1-TAP, state V; PDB: 6RI5 [3]), **(D)** Yvh1 on the cytoplasmic Rpl10-inserted pre-60S particle (PDB: 6N8O [6]). The indicated proteins have been highlighted in different colours; other r-proteins and AFs are coloured in light cyan and (pre-)rRNAs in light grey.

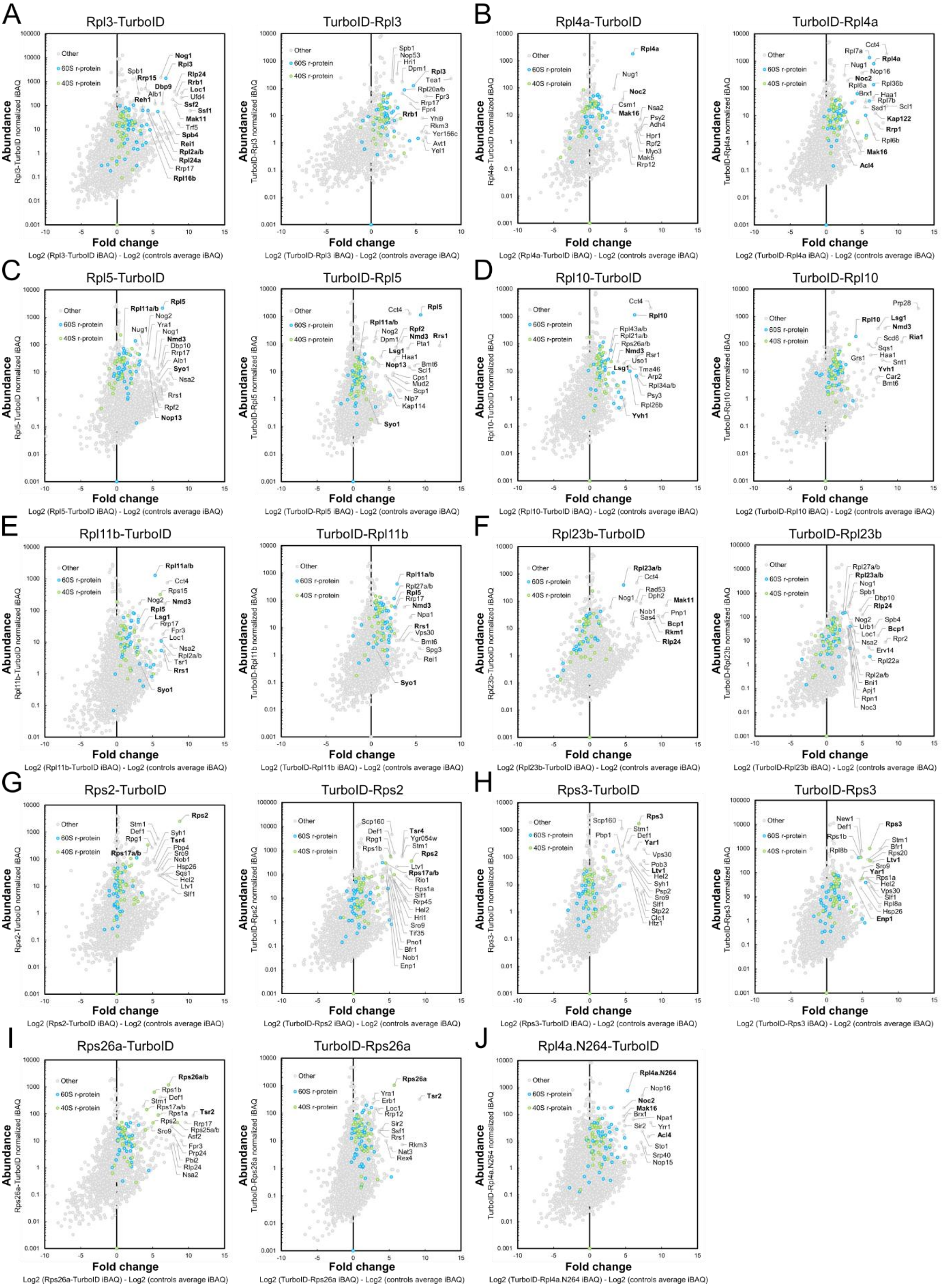


**Supplementary Figure S6. Identification of the known DCs in the proxiOMEs of their r-protein clients. (A-I)** Graphical representation of the TurboID results obtained with N- and C-terminally TurboID-tagged Rpl3 **(A)**, Rpl4a **(B)**, Rpl5 **(C)**, Rpl10 **(D)**, Rpl11b **(E)**, Rpl23b **(F)**, Rps2 **(G)**, Rps3 **(H)**, and Rps26a **(I)**. **(J)** TurboID result obtained with the C-terminally TurboID-tagged Rpl4 variant lacking the C-terminal extension (Rpl4a.N264). The employed bait r-protein is indicated above each graph, and the bait r-protein, its known DC, and selected enriched proteins are written in bold.

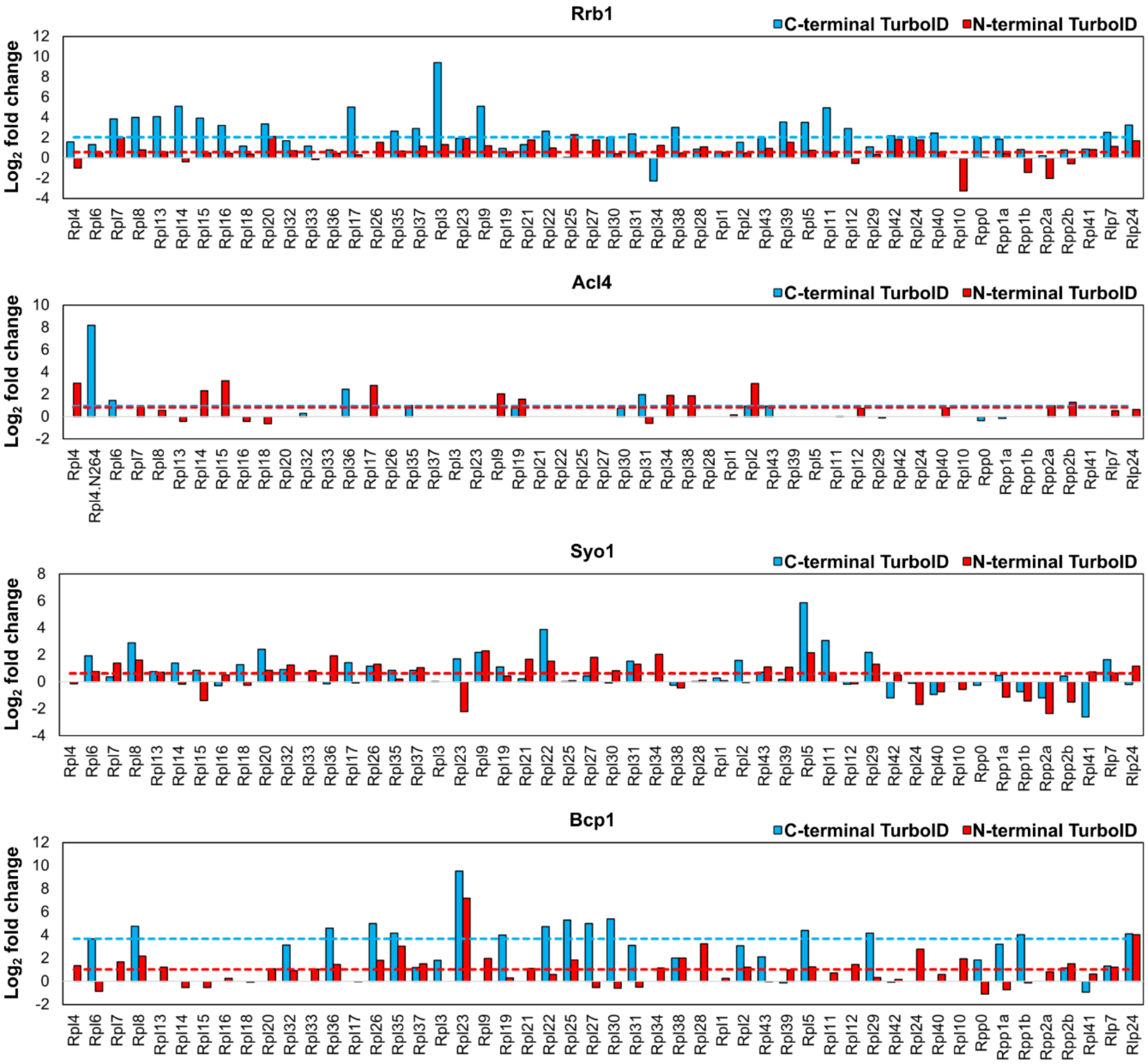


**Supplementary Figure S7. The DCs Rrb1, Acl4, Syo1, and Bcp1 are specifically enriched in the TurboID assays of their r-protein client.** Bar graphs showing the log2 fold enrichment of Rrb1, Acl4, Syo1, and Bcp1 in the TurboID assays of all N- (red bars) and C-terminally (blue bars) TurboID-tagged LSU r-proteins as well as the ones of Rlp7 and Rlp24. In the case of the Rpl3-TurboID bait and the TurboID-Rpl1 bait, their log2 fold enrichment corresponds to the average of the eight or, respectively, six experimental replicates. Dashed lines indicate the median log2 fold enrichment of an AF across all shown TurboID assays. The order of the LSU r-proteins (from left to right) is according to their first visualization in structures of early nucleolar to cytoplasmic pre-60S particles.

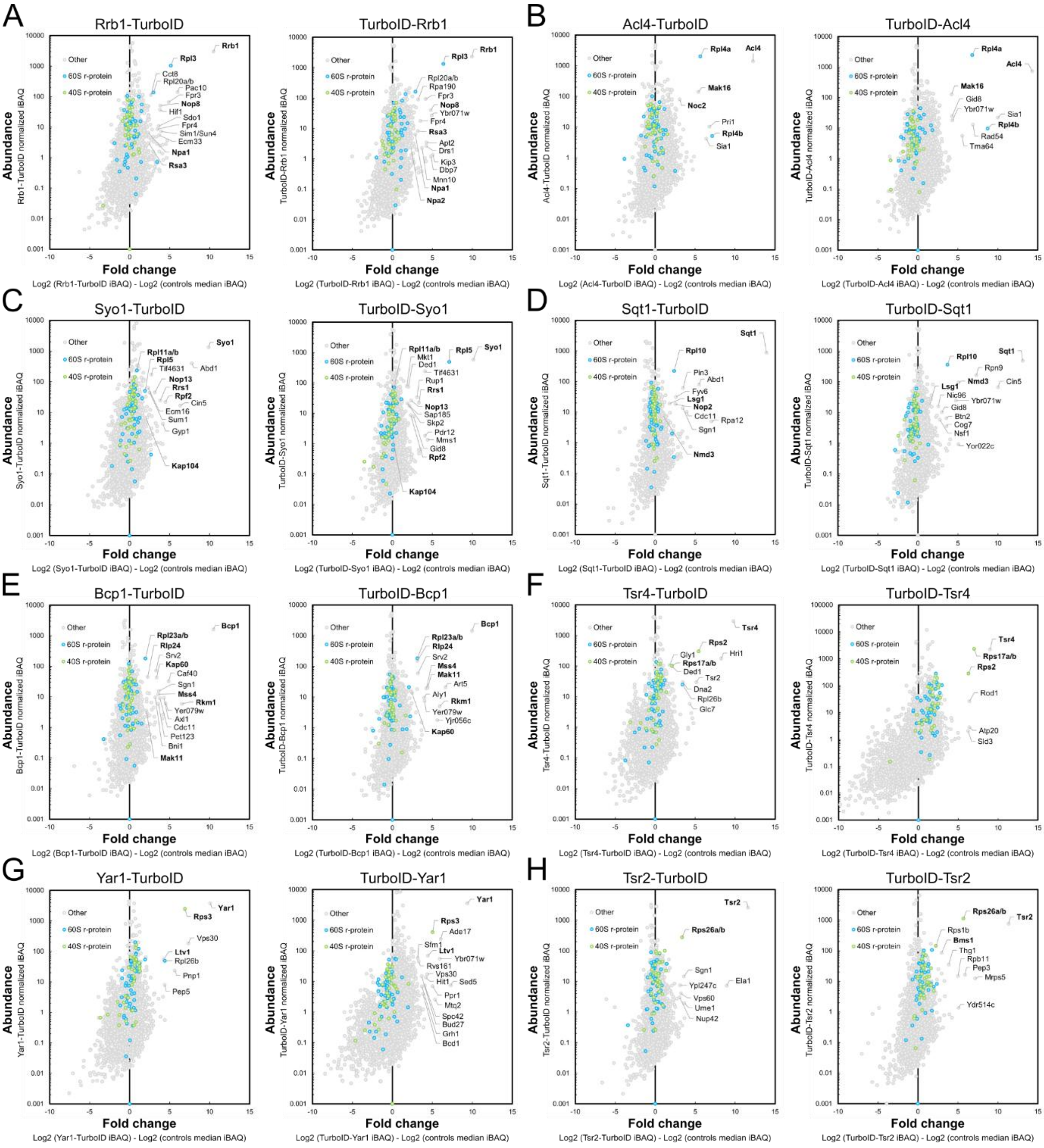


**Supplementary Figure S8. Identification of the known r-protein clients in the proxiOMEs of their DCs. (A-H)** Graphical representation of the TurboID results obtained with N- and C-terminally TurboID-tagged Rrb1 **(A)**, Acl4 **(B)**, Syo1 **(C)**, Sqt1 **(D)**, Bcp1 **(E)**, Tsr4 **(F)**, Yar1 **(G)**, and Tsr2 **(H)**. The abundance of each detected protein is displayed as its normalized intensity-based absolute quantification (iBAQ) value on the vertical axis, and its relative enrichment (fold change) on the horizontal axis as the log2 fold change of its normalized iBAQ abundance between the bait sample and the median of the remaining DC TurboIDs and the two respective control TurboIDs with prior imputation of missing values. The employed bait DC is indicated above each graph, and the bait DC, its known r-protein client(s), and selected enriched proteins are written in bold.

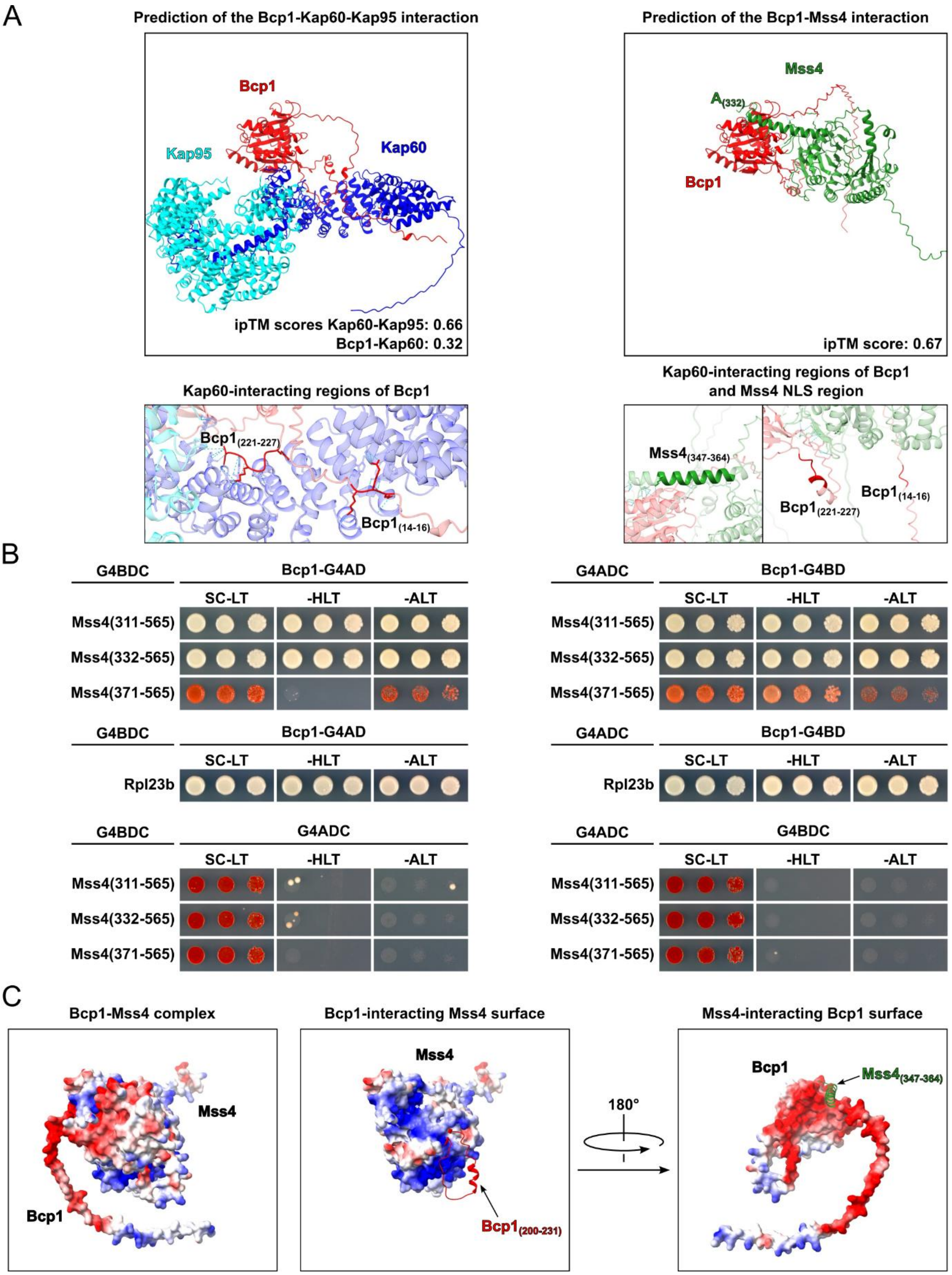


**Supplementary Figure S9. Bcp1 shields all prominent positively charged surfaces of Mss4. (A)** Cartoon representation of the AlphaFold3 model of the Bcp1-Kap60-Kap95 complex (upper left) and the Bcp1-Mss4 complex (upper right; for clarity of the representation, residues 1-331 of Mss4 have been hidden). The ipTM scores of the individual binary interactions are indicated. Close-up view highlighting the Kap60-interacting residues (14-16 and 221-227; dark red) of Bcp1 in contact with

Kap60 (lower left) or in the Bcp1-Mss4 complex (lower right). The location of the functional NLS region of Mss4 (residues 347-364; [7]) within the predicted Bcp1-Mss4 complex is highlighted in dark green. **(B)** Y2H interaction assays between Bcp1 and Rpl23 (positive control) or the indicated Mss4 segments. **(C)** Predicted electrostatic surface potential of the Bcp1-Mss4 complex (AlphaFold3 model; left), the Bcp1-interacting surface of Mss4 with residues 200-231 of Bcp1 in cartoon representation (middle), and the Mss4-interacting surface of Bcp1 with residues 347-364 of Mss4 in cartoon representation (right), revealing the respective negatively (Bcp1) or positively (Mss4) charged interaction interfaces.

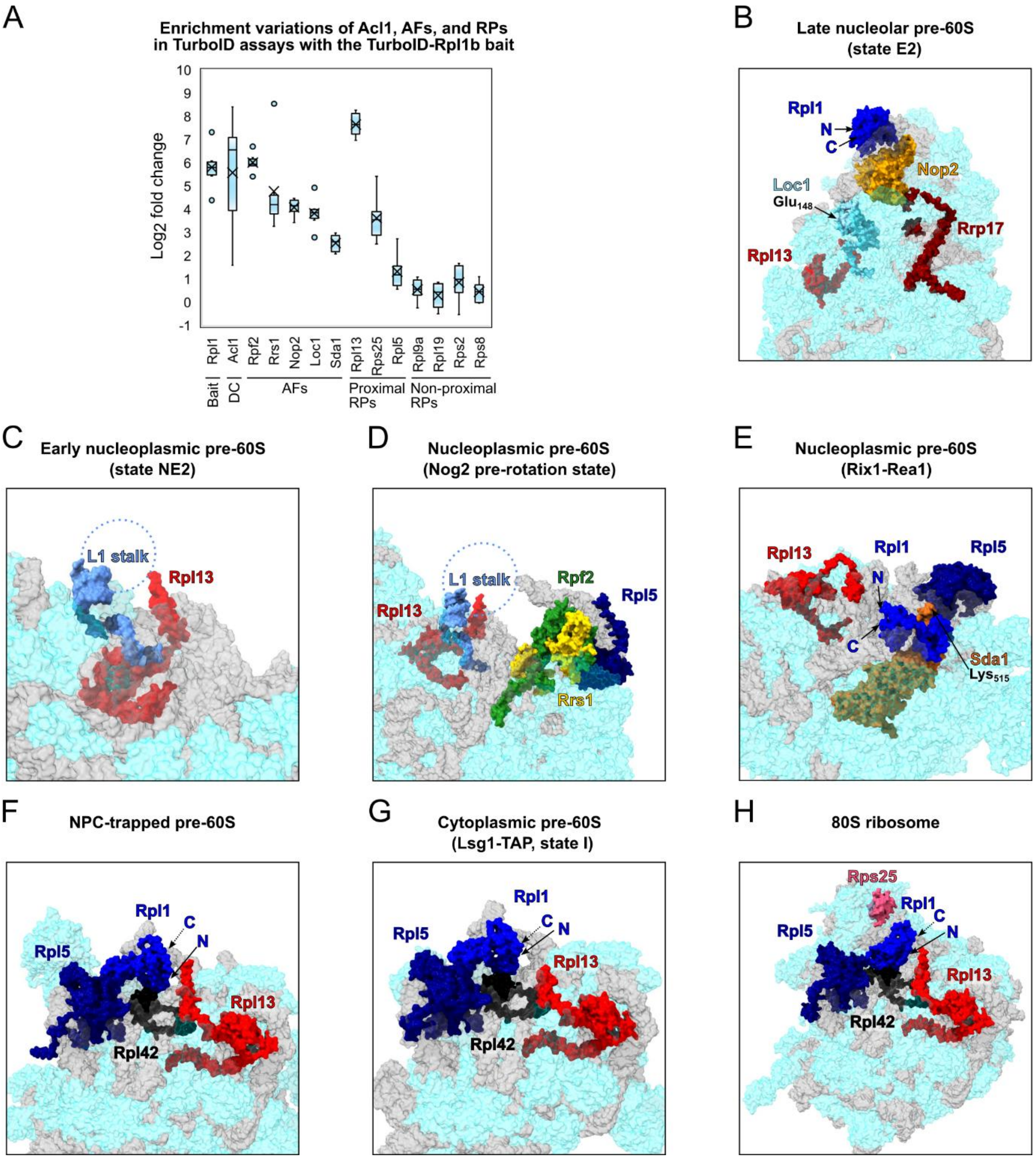


**Supplementary Figure S10. ProxiOME of Rpl1. (A)** Representation of the variation of the enrichment of the Rpl1 bait, its DC Acl1, proximal AFs, and abundantly detected r-proteins between the experimental replicates (n=6); r-proteins that are close to or far away from Rpl1, as indicated by available structures of pre-60S particles, mature 60S subunits, and 80S ribosomes, are referred to as proximal or non-proximal RPs. The variation of the enrichment is represented by boxes highlighting the interquartile range of each data set, while the whiskers indicate the minimal and maximal limits of the distribution; outliers are shown as dots, the median by a line, and the average by a cross. **(B-H)** Location of Rpl1 or the L1 stalk rRNA and their neighbouring proteins on: **(B)** the late nucleolar state E2 pre-60S particle (PDB: 7NAC [1]), **(C)** the early nucleoplasmic state NE2 pre-60S particle (PDB: 6YLY [8]), **(D)** the nucleoplasmic pre-60S intermediate of the Nog2-containing pre-rotation state (PDB: 7UOO [9]), **(E)** the nucleoplasmic Rix1-Rea1 pre-60S particle (PDB: 6YLH [8]), **(F)** the NPC-trapped pre-60S particle (PDB: 8HFR [10]), **(G)** the cytoplasmic Lsg1-TAP state I pre-60S particle

(PDB: 6RZZ [3]), and **(H)** the 80S ribosome (with the Rpl1-bound L1 stalk rRNA added from the PDB 4V7R 80S structure [11] and superposed on the PDB 4V88 80S structure [2]). The indicated proteins have been highlighted in different colours; other r-proteins and AFs are coloured in light cyan and (pre-)rRNAs in light grey.

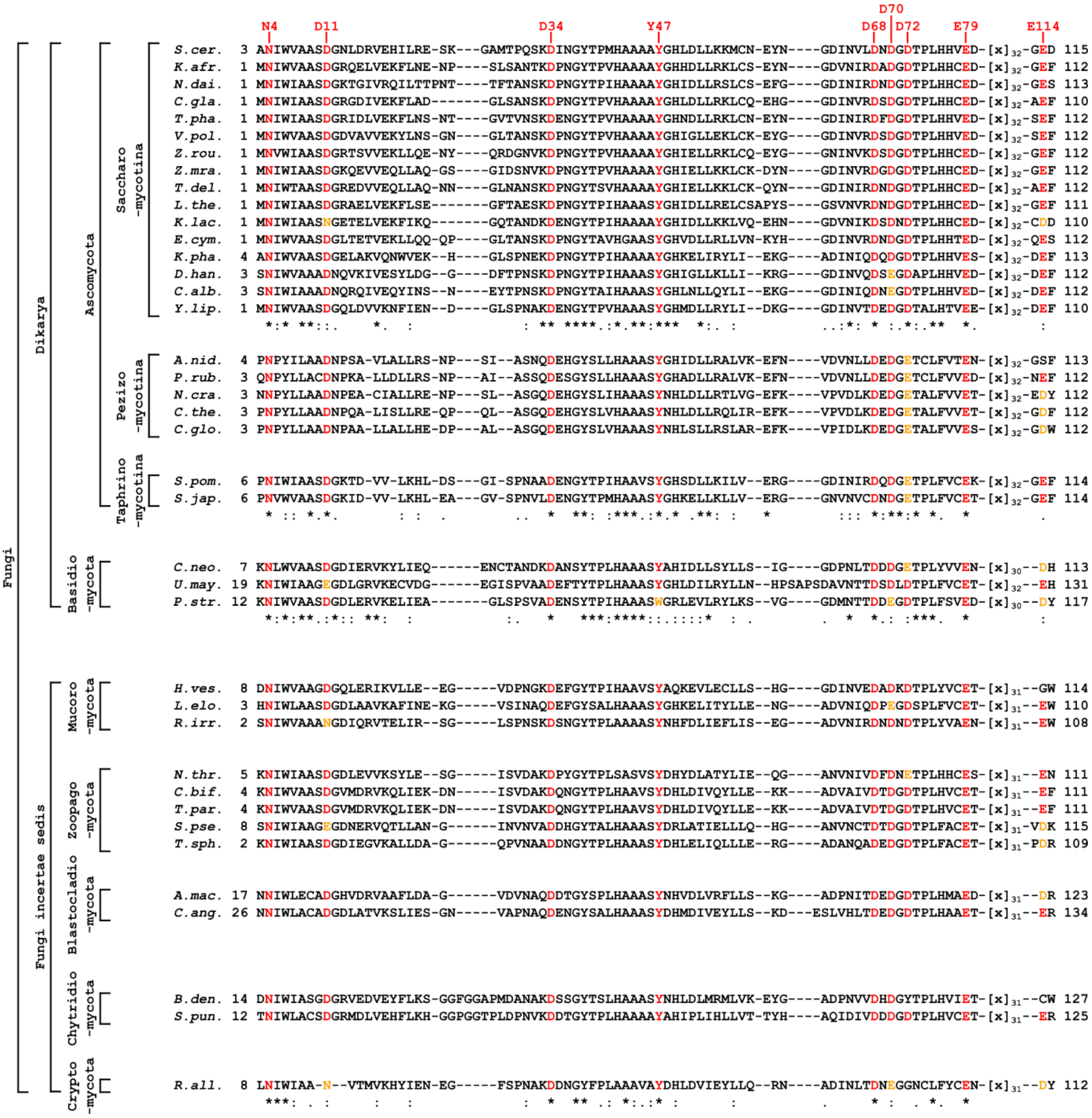


**Supplementary Figure S11. Acl1 and its identified Rpl1-interacting residues are conserved within the Fungi kingdom.** The amino acid sequence of *S. cerevisiae* Acl1 (*S.cer.*, UniProt: P25631) was aligned with the amino acid sequences of selected potential orthologous proteins, identified with the BLAST function of the UniProtKB [12], belonging to a specific subkingdom, phylum, or subphylum of the Fungi kingdom (see Materials and Methods). For clarity, only the alignment parts covering most of the N-terminal ankyrin repeat domain (residues 3-80 and 113-115 of *S. cerevisiae* Acl1) are shown, and interaction-mediating residues of *S. cerevisiae* Acl1 (indicated on top) that are conserved or semi-conserved are coloured in red or orange. Kingdom, subkingdom, phylum, and subphylum names are indicated. Species names are abbreviated as follows (with the UniProtKB or UniParc number of the respective protein referenced between parentheses): *Kazachstania africana*, *K.afr.* (H2B114); *Naumovozyma dairenensis*, *N.dai.* (G0WFQ9); *Candida glabrata*, *C.gla.* (Q6FWT6); *Tetrapisispora phaffii*, *T.pha.* (G8BT41); *Vanderwaltozyma polyspora*, *V.pol.* (A7TG26); *Zygosaccharomyces rouxii*, *Z.rou.* (C5DVQ9); *Zygotorulaspora mrakii*, *Z.mra.* (A0A7H9AWJ3); *Torulaspora delbrueckii*, *T.del.* (G8ZN72); *Lachancea thermotolerans*, *L.the.* (C5DBY1); *Kluyveromyces lactis*, *K.lac.* (Q6CTY4); *Eremothecium cymbalariae*, *E.cym.* (G8JPK3); *Komagataella phaffii*, *K.pha.* (C4R8E0);

*Debaryomyces hansenii*, *D.han.* (Q6BK78); *Candida albicans*, *C.alb.* (A0A1D8PHS4); *Yarrowia lipolytica*, *Y.lip.* (Q6C254); *Aspergillus nidulans*, *A.nid.* (Q5BDT5); *Penicillium rubens*, *P.rub.* (B6H7C4); *Neurospora crassa*, *N.cra.* (Q7RZ65); *Chaetomium thermophilum*, *C.the.* (G0S6G3); *Chaetomium globosum*, *C.glo.* (Q2H182), *Schizosaccharomyces pombe*, *S.pom.* (Q9UU77); *Schizosaccharomyces japonicus*, *S.jap.* (B6JYY9); *Cryptococcus neoformans*, *C.neo.* (Q5KM81); *Ustilago maydis*, *U.may.* (A0A0D1E9K4); *Puccinia striiformis*, *P.str.* (A0A2S4WKI7); *Hesseltinella vesiculosa*, *H.ves.* (A0A1X2GXH0); *Linnemannia elongata*, *L.elo.* (A0A197K2N3); *Rhizophagus irregularis*, *R.irr.* (U9U5S0); *Neoconidiobolus thromboides*, *N.thr.* (UPI002211511C); *Coemansia biformis*, *C.bif.* (A0A9W8CY32); *Tieghemiomyces parasiticus*, *T.par.* (A0A9W8ADQ8); *Syncephalis pseudoplumigaleata*, *S.pse.* (A0A4P9YSW2); *Thamnocephalis sphaerospora*, *T.sph.* (A0A4P9XN32); *Allomyces macrogynus*, *A.mac.* (A0A0L0S8J9); *Catenaria anguillulae*, *C.ang.* (A0A1Y2HCN5); *Batrachochytrium dendrobatidis*, *B.den.* (A0A177W6V9); *Spizellomyces punctatus*, *S.pun.* (A0A0L0HTG5); *Rozella allomycis*, *R.all.* (A0A075AZK2).

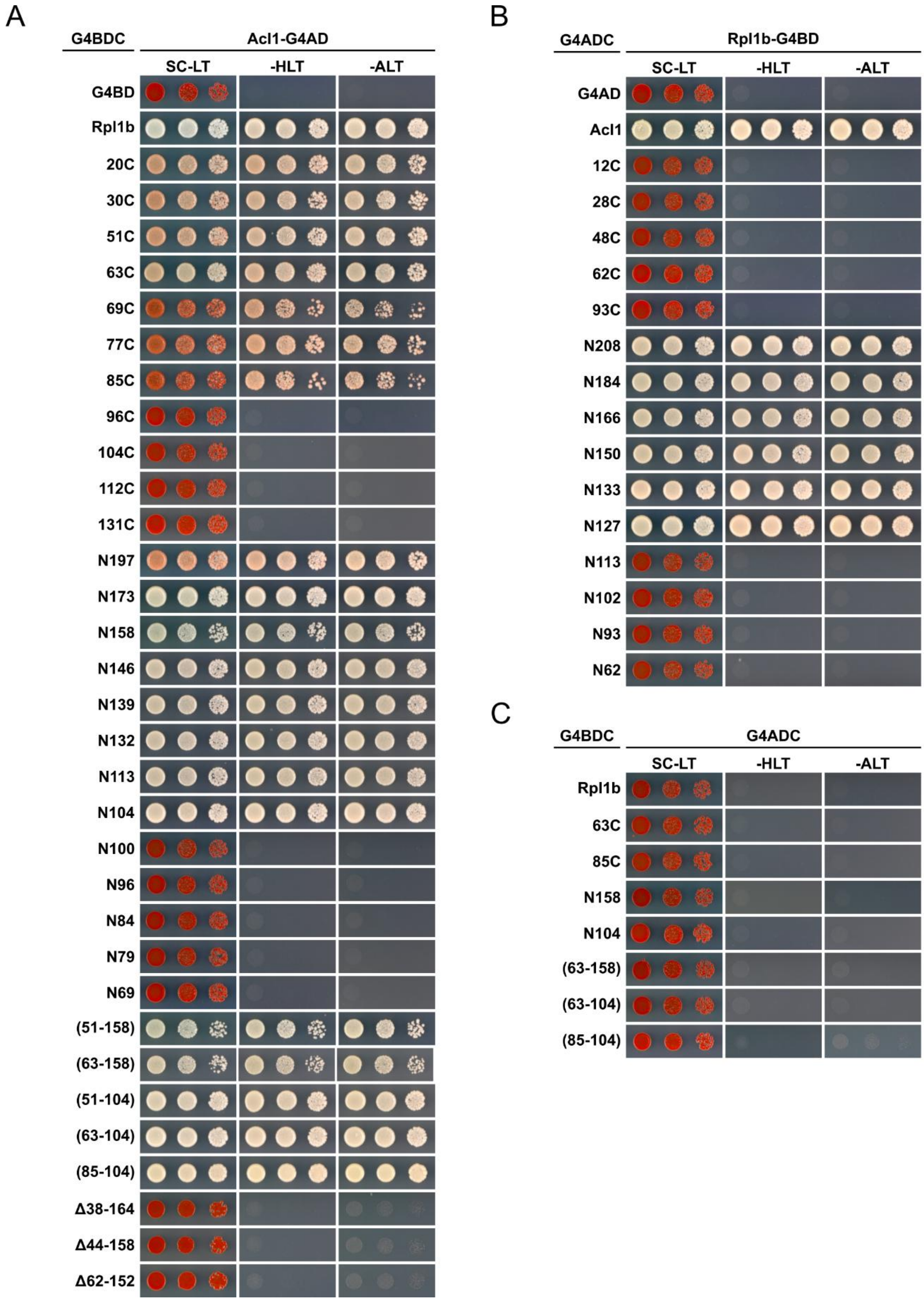


**Supplementary Figure S12. Acl1 interacts via its ankyrin repeat domain with the second domain of Rpl1. (A, B)** Complete data set for the mapping of the minimal respective interaction-mediating regions on Rpl1 **(A)** and Acl1 **(B)** by Y2H assays. **(C)** Negative control Y2Hs showing that the indicated, C-terminally G4BD-tagged Rpl1 variants do not self-activate the reporter genes when assayed together with the non-fused G4AD. Related to the data shown in Fig. 5A and B.

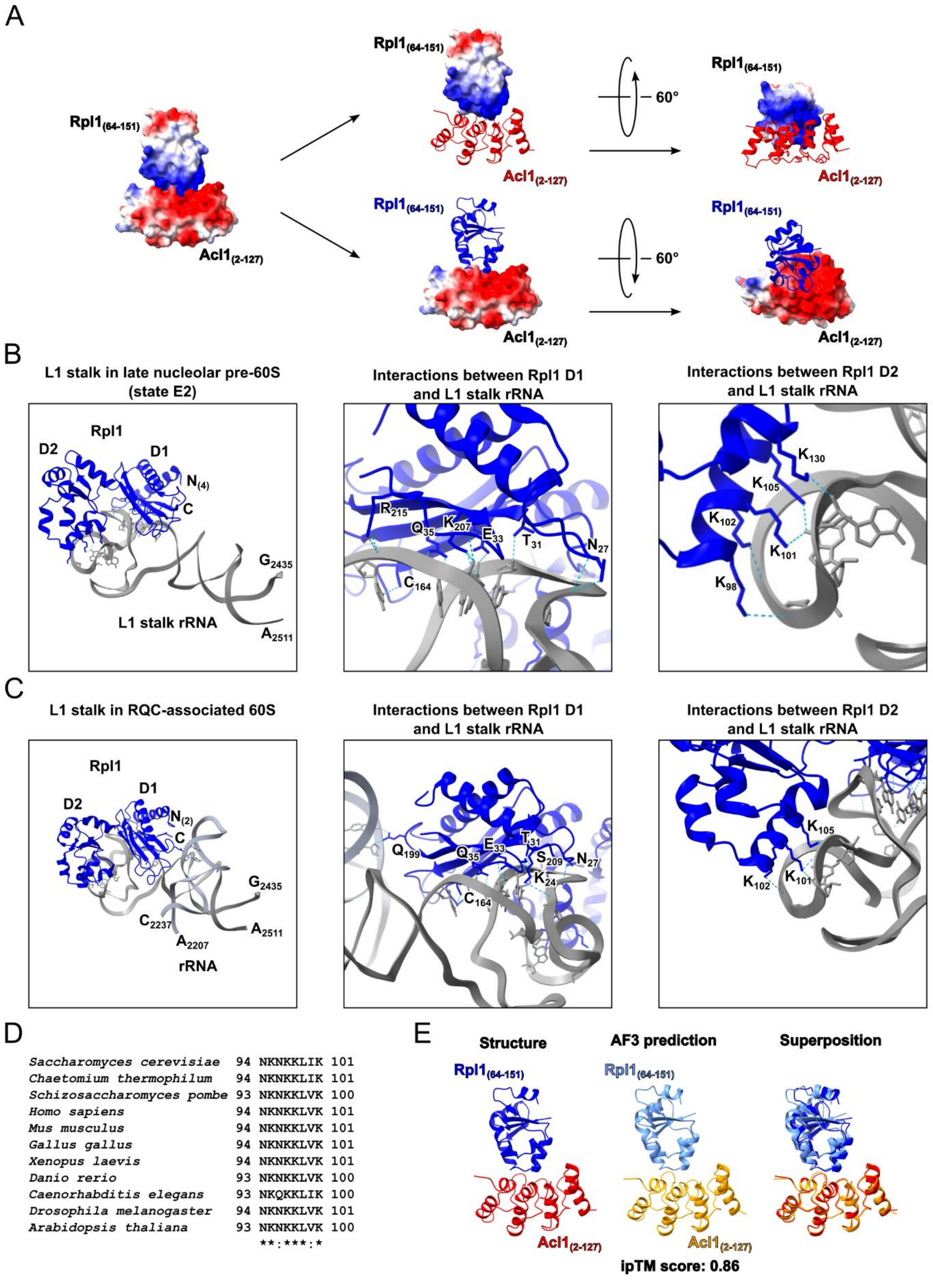


**Supplementary Figure S13. (A)** Representation of the electrostatic surface potential of the X-ray co-structure of the ankyrin repeat domain of Acl1 (residues 2-127) in complex with the second structural domain of Rpl1 (residues 64-151) with either Acl1 (in red) or Rpl1 (in blue) in cartoon representation to reveal the respective negatively (Acl1) or positively (Rpl1) charged interaction interfaces. **(B, C)** Representation of the interactions of Rpl1 with the L1 stalk rRNA as observed in the cryo-EM structure of: **(B)** the late nucleolar state E2 pre-60S particle (PDB: 7R7C [1]) and **(C)** an RQC-associated 60S subunit (PDB: 8AGX [13]). **(D)** The Acl1-interacting residues of Rpl1 are conserved in Rpl1 orthologues in metazoans, plants, and other fungi, as shown here for Rpl1 of *Chaetomium thermophilum* (UniProt: G0S4Z9), *Schizosaccharomyces pombe* (O74836), *Homo sapiens* (P62906), *Mus musculus* (P53026), *Gallus gallus* (F6SU35), *Xenopus laevis* (Q7ZYS8), *Danio rerio* (Q6PC69), *Caenorhabditis elegans* (Q9N4I4), *Drosophila melanogaster* (Q9VTP4), and *Arabidopsis thaliana* (P59230). **(E)** Cartoon representation of the ankyrin repeat domain of Acl1 (residues 2-127) in complex with the second structural domain of Rpl1 (residues 64-151) as observed in our experimental X-ray co-structure or the AlphaFold3 model (ipTM score 0.86). The superposition of the two structures reveals their high similarity.

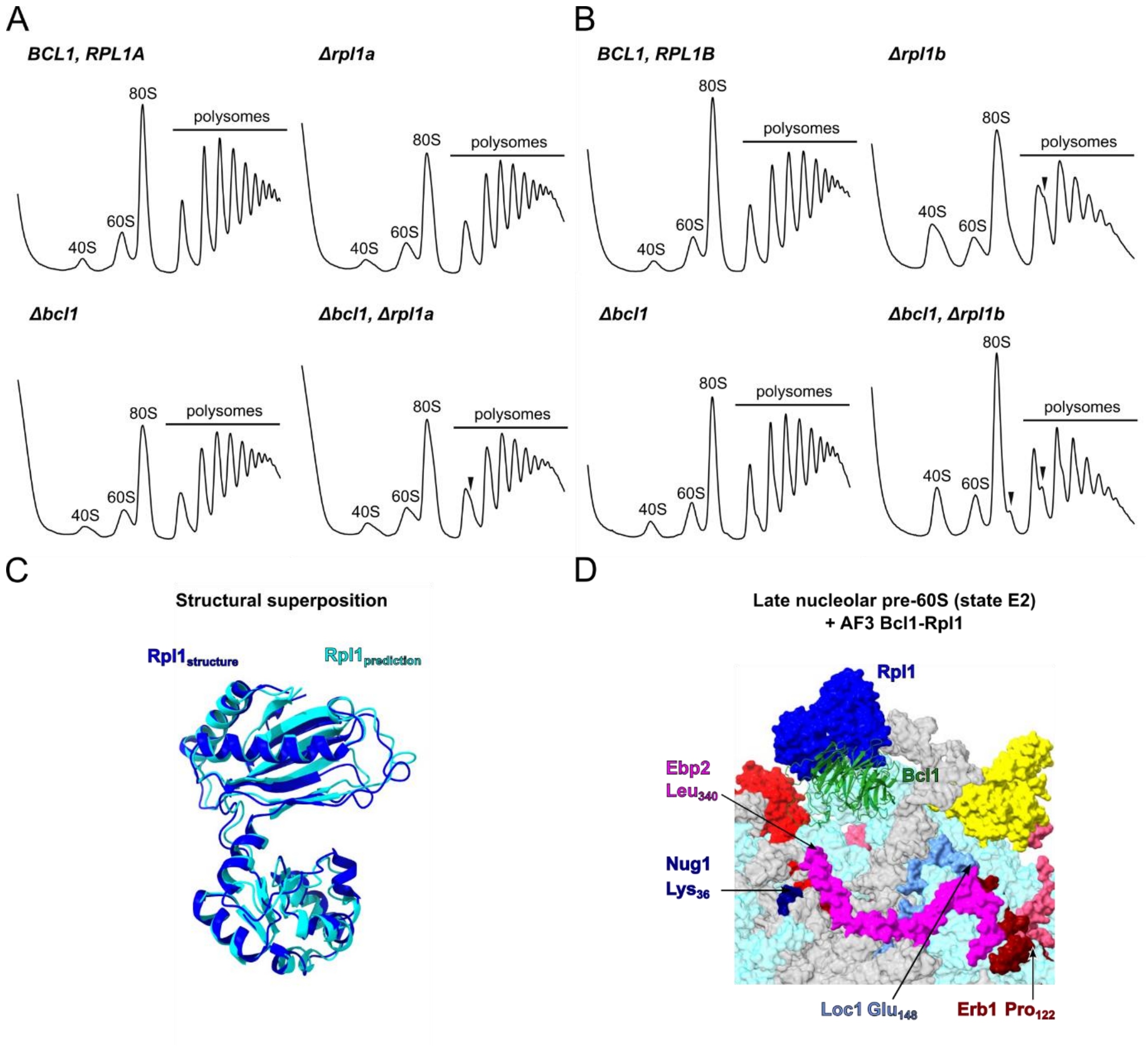


**Supplementary Figure S14. (A, B)** Polysome profiles of wild-type (*BCL1*, *RPL1A*), Δ*rpl1a*, Δ*bcl1*, and Δ*bcl1*/Δ*rpl1a* cells **(A)**, and of wild-type (*BCL1*, *RPL1B*), Δ*rpl1b*, Δ*bcl1*, and Δ*bcl1*/Δ*rpl1b* cells **(B)** grown at 30ºC in YPD medium. **(C)** Cartoon representation of the superposed predicted (extracted from the AlphaFold3 model of the Bcl1-Rpl1 complex) and observed (extracted from the late nucleolar state E2 pre-60S particle (PDB: 7NAC [1]) Rpl1 structures. **(D)** Close-up view of the superposition (shown in Fig. 8D) of the AlphaFold3 model of the Bcl1-Rpl1 complex onto Rpl1 within the late nucleolar state E2 pre-60S particle (PDB: 7NAC [1]), highlighting the first resolved residue of Erb1 and the last resolved residue of Ebp2, Nug1, and Loc1.

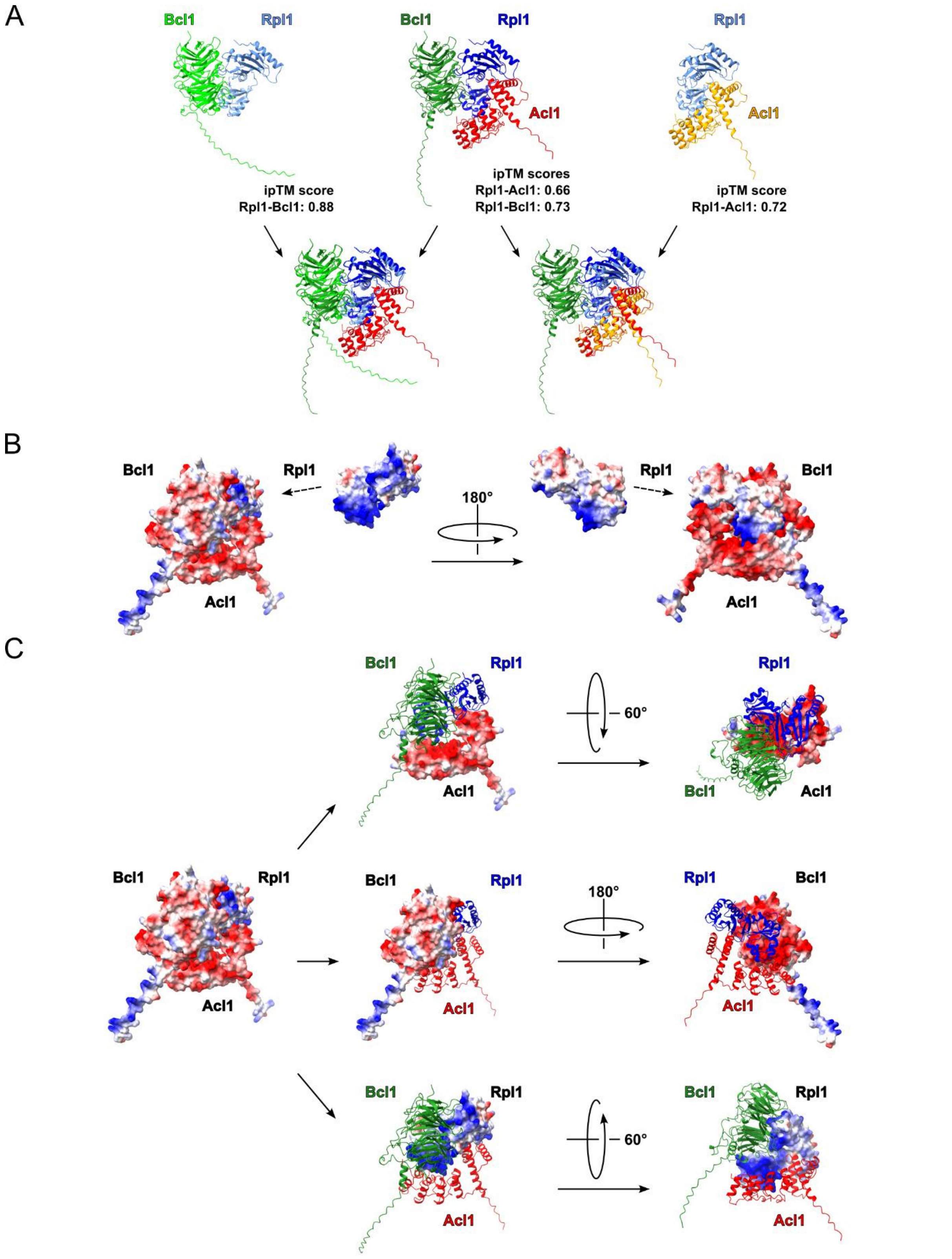


**Supplementary Figure S15.** **(A)** AlphaFold3 models of the Bcl1-Rpl1 (upper left), the Acl1-Rpl1-Bcl1 (upper middle), and the Acl1-Rpl1 (upper right) complexes. The lower panels show the superposition of the predicted Bcl1-Rpl1 or Acl1-Rpl1 complex with the predicted trimeric complex. The ipTM scores of the individual binary interactions are indicated. **(B, C)** Representation of the predicted electrostatic surface potential of the trimeric Acl1-Rpl1-Bcl1 complex (AlphaFold3 model)

**(B)** with either Bcl1 and Rpl1, Acl1 and Rpl1, or Acl1 and Bcl1 in cartoon representation to reveal the respective negatively (Acl1 and Bcl1) or positively (Rpl1) charged interaction interfaces **(C)**.

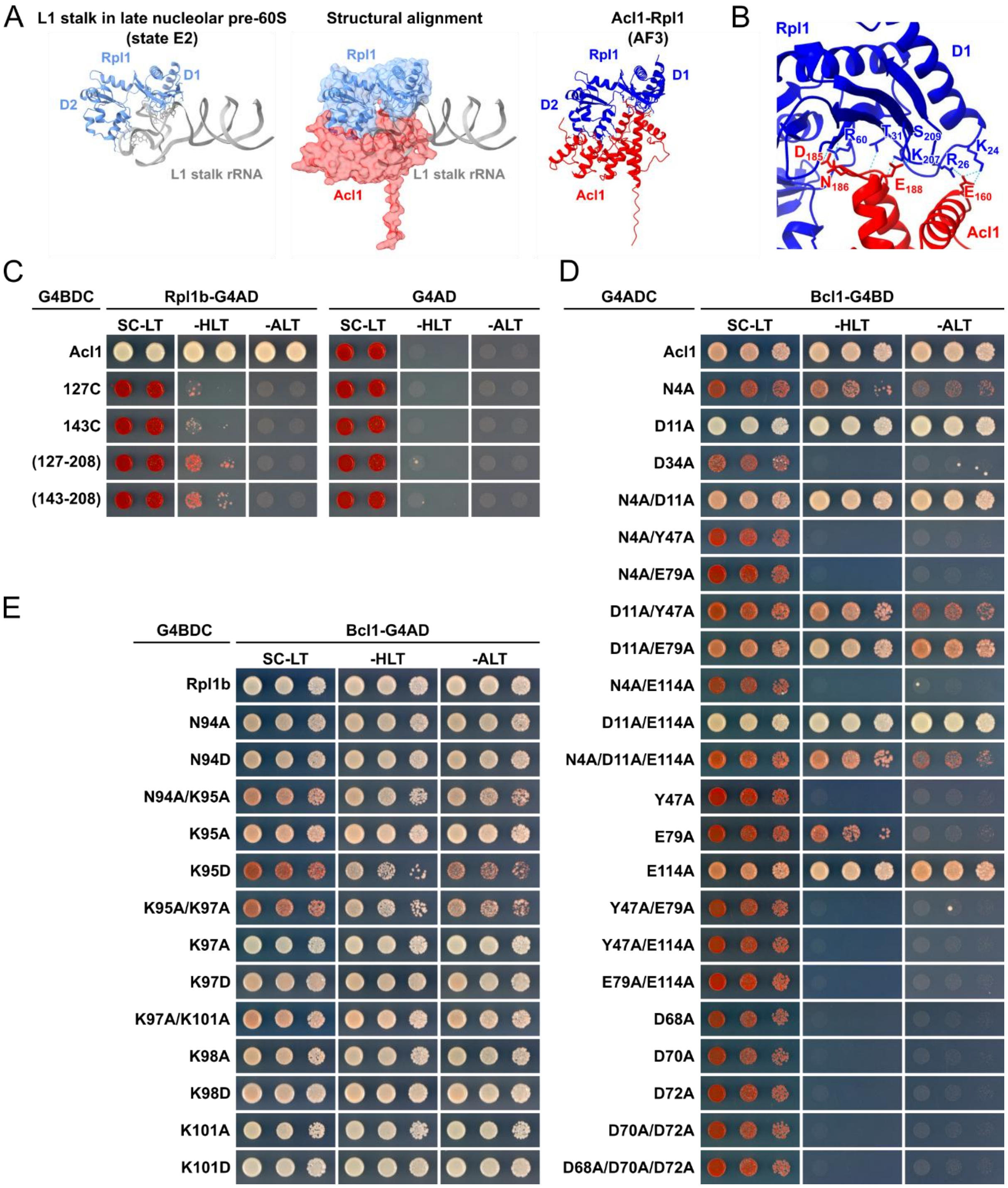


**Supplementary Figure S16. (A)** Interaction between Rpl1 (light blue) and the L1 stalk rRNA (grey) as observed on the late nucleolar state E2 pre-60S particle (left panel; PDB: 7R7C [1]). AlphaFold3 model of the Acl1-Rpl1 complex (right panel). Superposition of the AlphaFold3 model of the Acl1-Rpl1 complex and the Rpl1-bound L1 stalk rRNA (middle panel), revealing that Acl1 shields the rRNA-binding surfaces of Rpl1. **(B)** Close-up view of the predicted interaction between residues of the predicted three-helix fold within Acl1's C-terminal part and the first structural domain (D1) of Rpl1. **(C)** Y2H interaction assays between Rpl1 and full-length Acl1 or the indicated C-terminal segments of Acl1 (127C, residues 127-222; 143C, residues 143-222). The right panel shows the negative control Y2Hs with the non-fused G4AD, revealing that the indicated, C-terminally G4BD-tagged Acl1 variants do not self-activate the reporter genes. **(D, E)** Y2H interaction assays between Bcl1 and the indicated

Acl1 **(D)** or Rpl1 **(E)** mutant variants. Complete data set related to the data shown in Fig. 9F and G. Single-letter abbreviations for the amino acid residues are as follows: A, Ala; D, Asp; E, Glu; K, Lys; N, Asn; Y, Tyr.

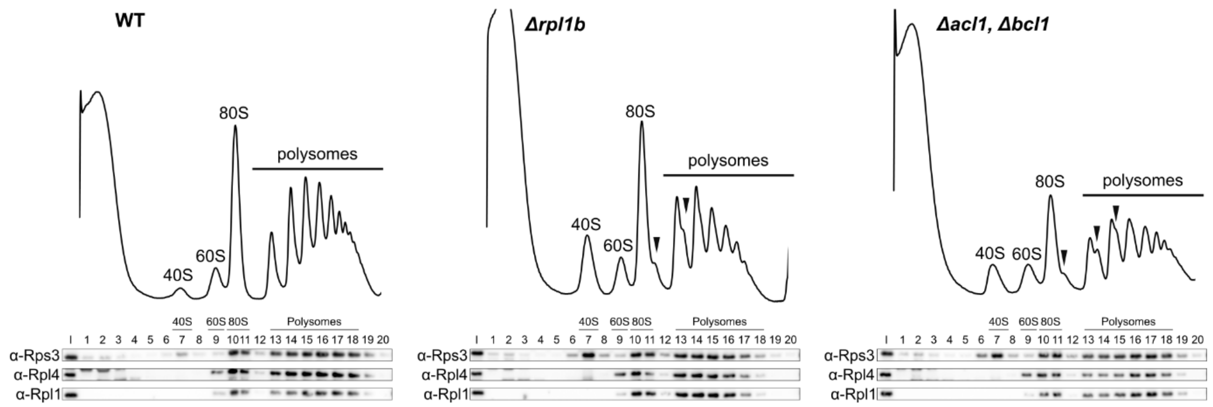


**Supplementary Figure S17.** Second experimental replicate of the sucrose gradient centrifugation and fractionation experiment shown in Fig. 10E that was used for the bar graph showing the average and standard deviation (n=2) of the normalized Rpl1/Rpl4 ratios in the 60S, 80S, and polysomal fractions of wild-type, Δ*rpl1b*, and Δ*acl1*/Δ*bcl1* cells in Fig. 10F.

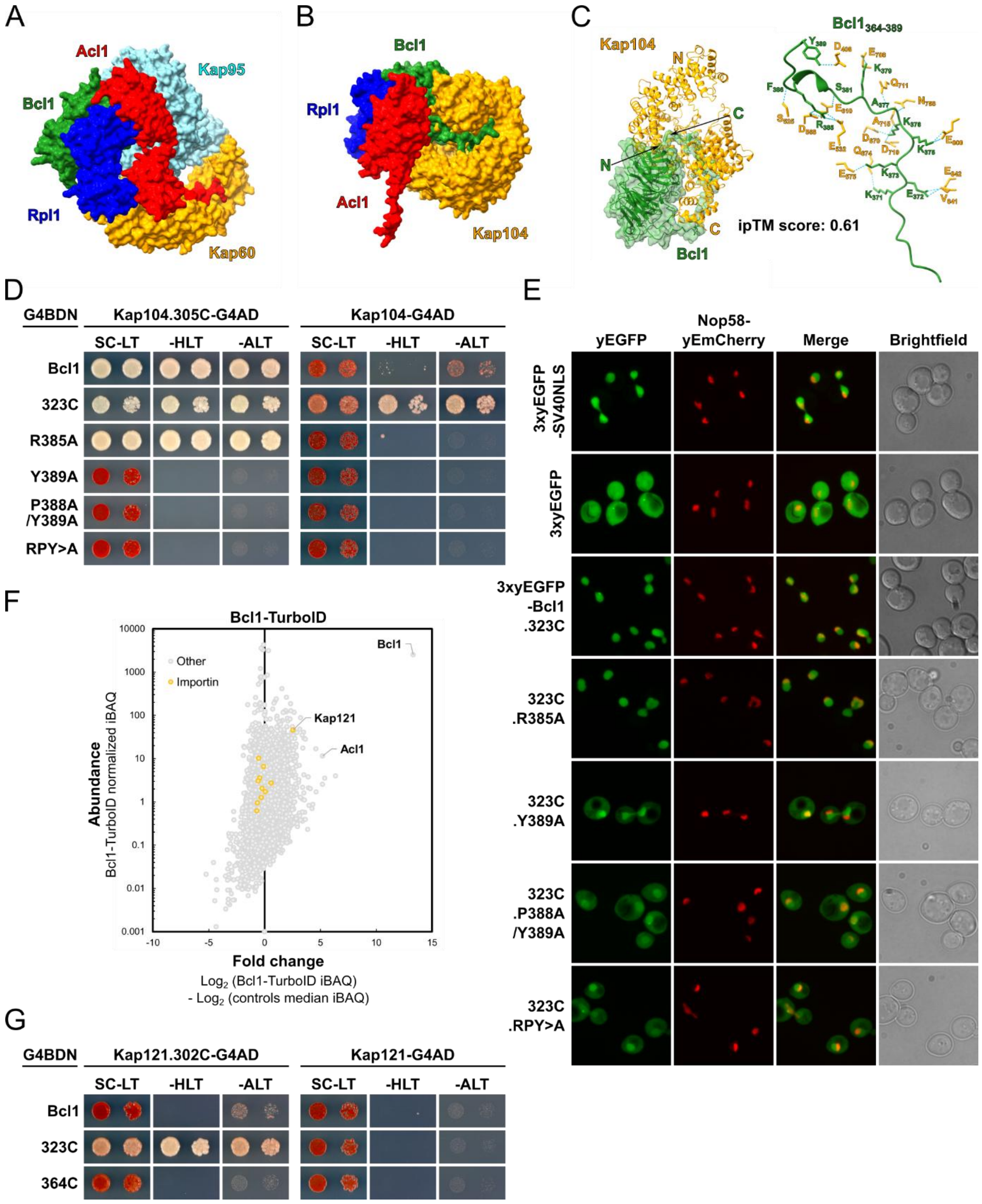


**Supplementary Figure S18**. **Acl1 and Bcl1 contain functional NLSs for the interaction with distinct importins. (A, B)** Surface representation of the AlphaFold3 model of the pentameric Kap95-Kap60-Acl1-Rpl1-Bcl1 complex **(A)** and the tetrameric Kap104-Bcl1-Rpl1-Acl1 complex **(B)**. **(C)** AlphaFold3 model of the binary Kap104-Bcl1 complex in cartoon representation with Bcl1 in semi-transparent surface representation (left). Close-up view showing the molecular details of the predicted interaction between Kap104 and the C-terminal bPY-NLS of Bcl1, with interacting residues being labelled and the observed H-bonds being depicted as dotted lines (right). **(D)** Y2H interaction assays between N-terminally G4BD-tagged (G4BDN) Bcl1, Bcl1.323C, and the indicated mutant variants of Bcl1.323C and C-terminally G4AD-tagged full-length Kap104 (right) or the N-terminally truncated

Kap104.305C variant (left). Single-letter abbreviations for the amino acid residues are as follows: A, Ala; P, Pro; R, Arg; Y, Tyr. **(E)** The nuclear targeting activity of Bcl1.323C and the indicated mutant variants thereof was assessed by fluorescence microscopy in cells grown at 30ºC in SC-Leu medium. The 3xyEGFP and 3xyEGFP-SV40NLS control proteins and the N-terminally 3xyEGFP-tagged Bcl1.323C fusion proteins were expressed from plasmid under the control of the *ADH1* promoter in cells expressing the nucleolar marker protein Nop58-yEmCherry from the genomic locus. **(F)** Graphical representation of the TurboID result obtained with C-terminally TurboID-tagged Bcl1 (Bcl1-TurboID). The Bcl1 bait protein, Acl1, and the enriched importin Kap121 are written in bold; orange dots highlight the location of importins on the graph. **(G)** Y2H interaction assays between N-terminally G4BD-tagged (G4BDN) Bcl1, Bcl1.323C, and Bcl1.364C and C-terminally G4AD-tagged full-length Kap121 (right) or the N-terminally truncated Kap121.302C variant (left).

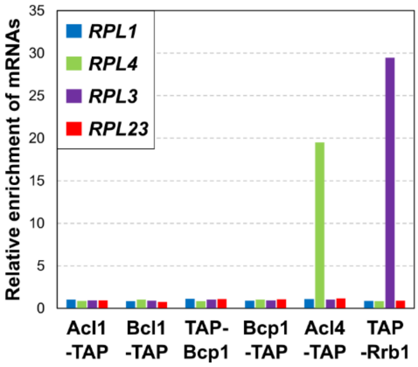


**Supplementary Figure S19. Co-translational capturing assay.** The DCs Acl1 (Acl1-TAP), Bcl1 (Bcl1-TAP), Bcp1 (TAP-Bcp1 and Bcp1-TAP), and, as positive controls, Acl4 (Acl4-TAP) and Rrb1 (TAP-Rrb1) were affinity purified (IgG-Sepharose pull-down) from extracts of cycloheximide-treated cells and the associated RNA was isolated from the TEV eluates. Each of the six DC purifications was assessed for their content of the four RPG mRNAs (*RPL1*, *RPL3*, *RPL4*, and *RPL23*) by real-time qRT-PCR. For each cDNA, real-time qPCRs were performed in triplicates. The bar graph shows the relative enrichment of the four RPG mRNAs in the six DC purifications.

#### SUPPLEMENTARY TABLE

**Supplementary Table S1:** Data collection and refinement statistics

| | **Rpl1(63-158)-Acl1.N127** |
|---|---|
| **Data collection** | |
| X-ray source | ESRF, Grenoble<br>ID30A-3 (MASSIF-3) |
| Detector | Eiger1 X 4M |
| Wavelength (Å) | 0.967697 |
| Space group | $P6_522$ |
| Cell dimensions ($a$, $b$, $c$ (Å)) | 101.65, 101.65, 108.13 |
| Resolution (Å)* | 46.07-2.55 (2.70-2.55) |
| Total reflections | 445071 |
| Multiplicity | 39.52 |
| Unique reflections | 11261 |
| Completeness (%) | 100.0 (100.0) |
| $R_{\mathrm{meas}}$ (%) | 29.5 (222.0) |
| $CC_{1/2}$ | 99.8 (52.3) |
| $I/\sigma(I)$ | 17.87 (2.78) |
| Mosaicity (°) | 0.221 |
| Wilson $B$-factor (Å$^2$) | 57.24 |
| **Refinement** | |
| Resolution (Å) | 46.07-2.55 |
| $R_{\mathrm{work}}$, $R_{\mathrm{free}}$ | 0.195, 0.242 |
| Reflections (working, test set) | 10698, 563 |
| Completeness for range (%) | 100.0 |
| r.m.s.d. from ideal | |
| Bond lengths (Å) | 0.011 |
| Bond angles (°) | 1.190 |
| Total number of atoms | 1663 |
| Mean $B$ value (Å$^2$) | 63.94 |

*Values in parentheses are for highest-resolution shell
